# Structural perspective on the design of selective DYRK1B inhibitors

**DOI:** 10.1101/2022.12.23.521429

**Authors:** Przemyslaw Grygier, Katarzyna Pustelny, Filipe Menezes, Malgorzata Jemiola-Rzeminska, Piotr Suder, Grzegorz Dubin, Anna Czarna

## Abstract

DYRK1B has been recently recognized as a critical therapeutic target in oncology and non-alcoholic fatty liver disease. However, the lack of structural information has constrained the development of selective inhibitors for DYRK1B. Here, we employed recombinant protein production, activity assays, and crystallization to elucidate the structure of DYRK1B. We present a crystal structure of DYRK1B in complex with a known inhibitor, AZ191. For comparative analysis, we provide the crystal structure of the closely related DYRK1A kinase in complex with AZ191. Our analysis identifies the exclusiveness of the binding site in the hinge region of DYRK1B, which is pivotal for selective inhibitor design. Quantum mechanical calculations reveal a notable difference in the accessibility of the catalytic lysine between DYRK1B and DYRK1A, offering avenues for distinguishing binders to these kinases. Our findings mark a significant advancement in the quest for specific DYRK1B inhibitors, potentially offering focused efficacy compared to the current dual-specificity inhibitors targeting both DYRK1B and DYRK1A.

## Introduction

Protein phosphorylation is the most widespread post-translational modification facilitating cellular signal transduction.^1^ Deregulation of kinase signalling underlies multiple human diseases. However, one of the challenges in exploring phosphorylation as a drug target arises in designing selective inhibitors addressing over 500 protein kinases of the human kinome.^2,3^

Aberrant activity of dual-specificity tyrosine phosphorylation-regulated kinases (DYRKs) has been linked to neurological disorders, diabetes, and cancer. DYRKs belong to a larger family of CMGC kinases, which includes cyclin-dependent kinases (C), mitogen-activated protein kinases (M), glycogen synthase kinases (G), and CDC-like kinases (C). DYRK1A and DYRK1B localize mainly in the nucleus, while DYRK2, DYRK3, and DYRK4 reside predominantly in the cytoplasm.^4^ The activity is regulated by autophosphorylation within the Tyr-Xxx-Tyr motif in the activation loop.^5^ The uniqueness of DYRKs is their ability to autophosphorylate on tyrosine residues while phosphorylating other substrates exclusively on serine and threonine.^6^ Within the DYRK family, the DYRK1A and 1B, two closely related paralogs, are of major interest as drug targets. This results from their multifaceted roles in critical cellular processes that are dysregulated in various diseases. Their involvement in cell cycle regulation, tissue regeneration, and neuronal signalling pathways presents a unique opportunity to develop targeted therapies that could address mechanisms underlying cancer, diabetes, or neurodegenerative diseases.

DYRK1B is overexpressed in certain types of cancer where its role is better understood than that of DYRK1A. The overexpression of DYRK1B maintains cellular quiescence^7^ and enhances cell survival by reducing reactive oxygen species (ROS) levels.^8^ It has been demonstrated that genetic depletion of DYRK1B leads to cell cycle re-entry and apoptosis.^9^ Additionally, the DYRK1B protein level increases upon DYRK1A silencing, suggesting a compensatory mechanism.^10^ DYRK1A, with its gene located at the Down Syndrome Critical Region (DSCR) of chromosome 21, is widely considered a potential drug target in neurological disorders, including Down Syndrome (DS), Alzheimer’s, and Parkinson’s disease.^11–14^ In the realm of regenerative medicine, both DYRK1A and 1B are pivotal for human beta cell regeneration, with their inhibition showing promise in treating diabetes by promoting beta cell proliferation.^11,15–17^ Although DYRK1B silencing in human islets had almost no effect on beta-cell proliferation, simultaneous silencing of DYRK1A and 1B resulted in significantly higher rates of proliferation than silencing DYRK1A alone, suggesting that dual inhibition also has its applications. As such, DYRK1B is potentially an interesting drug target for certain cancers. Collectively, these findings position DYRK1A and 1B as versatile targets for drug discovery, with implications across oncology, diabetes management, and neurodegeneration.

Regarding inhibitor development, the conserved catalytic domain of DYRKs is the commonly targeted site. Features defining the active site include a glycine-rich loop and DFG motif involved in ATP binding.^18,19^ The DFG motif adopts an “in” conformation in active kinases where the aspartic acid residue coordinates a Mg^2+^ ion bound to the β- and γ- phosphate groups of ATP. Other characteristic features include a gatekeeper residue (phenylalanine) guarding access to the ATP binding pocket^20^ and the Tyr-Xxx-Tyr motif determining kinase activity.^21^ The catalytic domain is flanked at the N-terminus by the DYRK homology box (DH), vital for autophosphorylation during the kinase maturation^6,22^ and at the C-terminus by the PEST sequence (proline, glutamic acid, serine, and threonine enriched) promoting rapid degradation^23^ (Figure 1). DYRK1A is additionally characterized by a histidine-rich region (HRD) responsible for localization to nuclear speckles, a feature that is absent in DYRK1B. This region is characteristic of several proteins expressed in the nervous system.^24^

**Figure 1.**
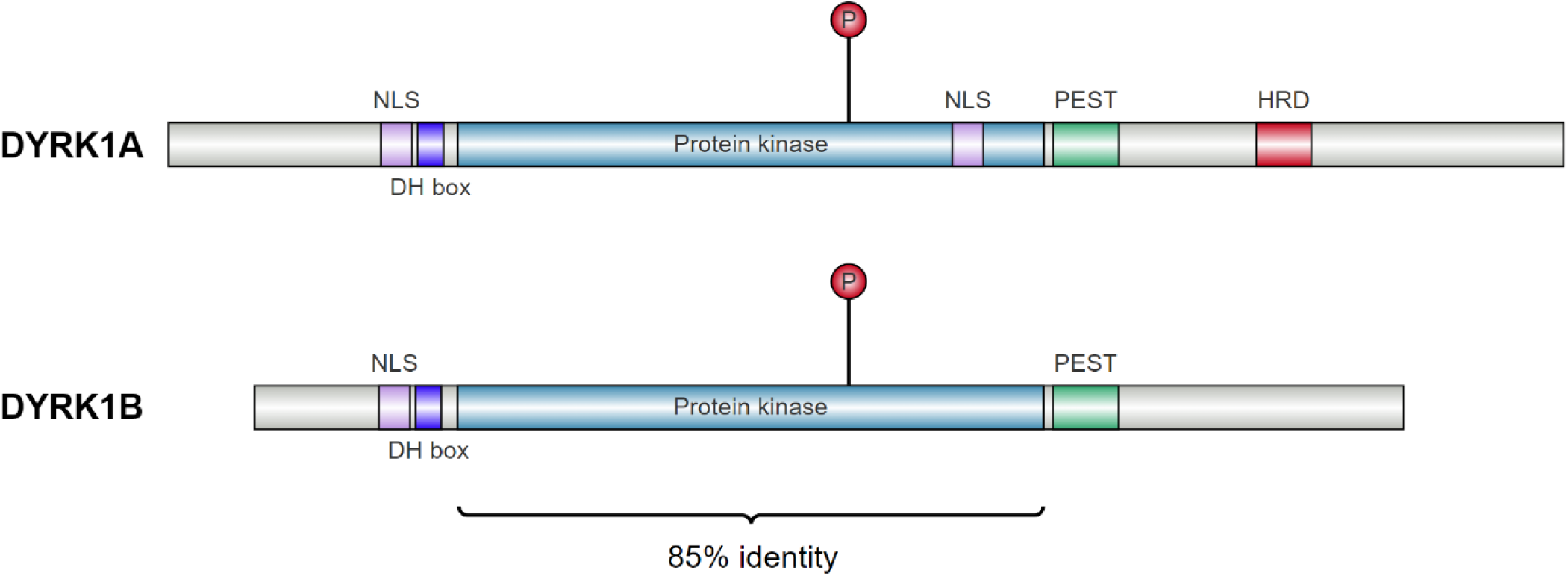
Schematic arrangement of domains in human DYRK1A and DYRK1B.

Harmine is the most common inhibitor used in investigating the roles of DYRK1A and 1B in health and disease. Among others, it was shown that harmine inhibits tau hyperphosphorylation, a process relevant to Alzheimer’s Disease pathology^25^ and exhibits a β-cell pro-proliferative effect.^15,26^ However, the inhibitor is equally effective against both kinases,^27^ hindering the possibility of kinase-specific studies. Various other inhibitors of DYRKs were reported, including AZ191,^28^ but none is specific enough to distinguish the kinases. Selective inhibitors are needed to understand the specific effects of inhibiting each paralogue.

At the same time, selective inhibitors may be of interest in treating DYRK1B-related cancers.^29,30^ However, developing selective inhibitors for closely related molecular targets is challenging, and the lack of a crystal structure of DYRK1B further complicates the effort. In our work, we provide the first crystal structure of DYRK1B in complex with AZ191, which will facilitate the future development of selective inhibitors.

## Results

### The pivotal role of autophosphorylation in maintaining DYRK1B stability

In the process of purifying the kinase domain of DYRK1B, it was observed that a substantial portion of the kinase aggregated in the insoluble fraction after cell lysis. This observation is particularly relevant considering the established importance of tyrosine phosphorylation within the Tyr-Xxx-Tyr motif for the enzyme’s catalytic functionality and structural stability, as documented in prior studies.^22,31,32^ Given this background, we hypothesized that the autophosphorylation of DYRK1B might be suboptimal when expressed in *Escherichia coli*. To investigate this, we specifically assessed the phosphorylation status of Tyr273. This was achieved through mass spectrometry analysis conducted on the whole cell lysate.

The Tyr273 was clearly visible during cell lysate analysis, mainly in the unphosphorylated form. Specifically, peptides: [253-277], [257-277], [157-280], and [270-280] (Uniprot ID Q9Y463) were identified based on their numerous MS/MS spectra (details in supplement: „identification report”). We detected Tyr273 phosphorylation in some of these peptides; however, the ratio between phosphopeptide and their unphosphorylated forms appeared to be low. To estimate the level of Tyr273 phosphorylation, we compared peak areas of the corresponding peptides in both forms. We focused on the ions at m/z = 730.3836^4+^ (unphosphorylated [253-277]) and at m/z = 750.3749^4+^ (phosphorylated [253-277]). The identities of both peptides were verified by their fragmentation spectra to confirm the sequences and phospho-residue (Figures S1 and S2). Based on the area under the corresponding extracted ion chromatograms, we calculated that the phosphorylation efficiency was close to 14% (Figure S3). Notably, the purified DYRK1B exhibited complete phosphorylation at Tyr273, strongly suggesting that the unphosphorylated pool is unstable and thus lost during the purification process.

### AZ191, a potent and selective DYRK1B inhibitor, enables paralog differentiation

The thermal-shift assay revealed that the catalytic domain of DYRK1A and DYRK1B has a markedly distinct thermal profile despite the high sequence identity (over 85%). The kinase domain of DYRK1A (124-490) was more stable with a melting point of 56 °C (T_m_), while the thermal denaturation of the kinase domain of DYRK1B (78-442) was observed at 47 °C (T_m_) (Figure 2B). This difference in melting temperatures aligns with previous observations and supports the distinct stability profiles of these two proteins^33^. Both DYRK1s’ stability was improved, and their melting temperature was raised by binding of harmine or AZ191. Notably, AZ191 at 10 μM induced a remarkable 12 °C shift in T_m_ of DYRK1B, indicating a strong stabilizing effect. AZ191 had a less pronounced impact on DYRK1A, with only a 4 °C shift observed. On the other hand, Harmine exhibited a moderate stabilizing effect, causing a 1.5 °C and 2.5 °C increase in melting temperatures of DYRK1A and DYRK1B, respectively (Figure 2B). To provide a more comprehensive context for our findings regarding the significant stabilization of DYRK1B by the AZ191 inhibitor, we conducted a Cellular Thermal Shift Assay (CETSA). The thermal stability was assessed for the full-length DYRK1B expressed in HEK293 cells and treated with the inhibitor (10 µM) or DMSO. For DYRK1B and DYRK1B-AZ191, denaturation occurred gradually over a wide temperature interval. However, an increased thermal stability of the full-length DYRK1B was observed following treatment with the AZ191 inhibitor (Figure S4). This provides evidence that AZ191 binds to and enhances the stability of both the isolated kinase domain and full-length DYRK1B.

**Figure 2.**
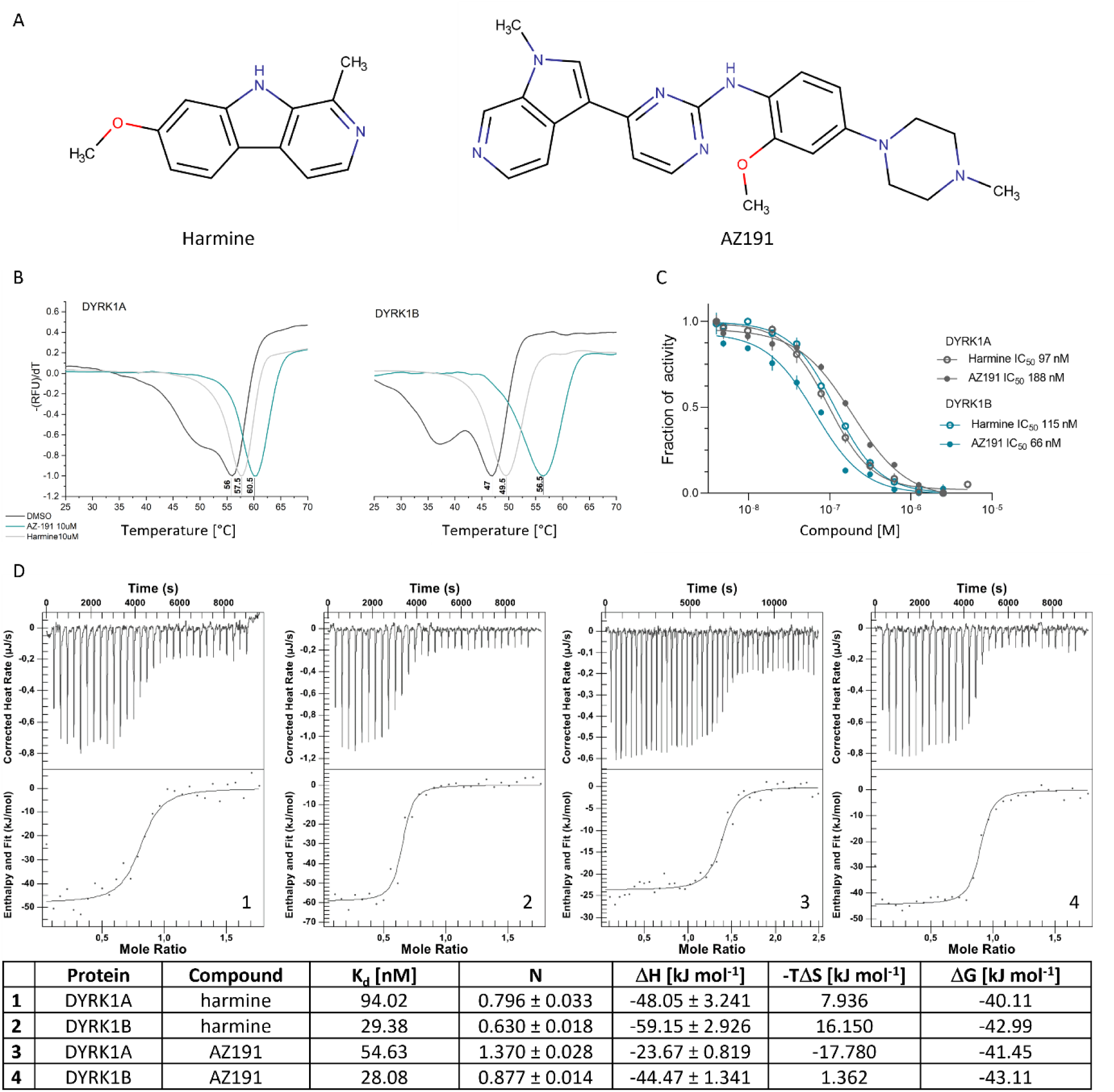
Biochemical analysis of DYRK1’s interactions with harmine and AZ191. (A) Structure of harmine and AZ191. (B) Thermal denaturation curves (first derivative) of DYRK1A (left panel) and DYRK1B (right panel) kinase domains after treatment with harmine, AZ191, or DMSO as determined in the Sypro Orange dye monitored thermal shift assay. (C) Inhibitory activity of AZ191 and harmine against DYRK1A and DYRK1B determined in ADP-Glo kinase activity assay and (D) ITC.

The AZ191-induced thermal stabilization of DYRK1A and DYRK1B kinases prompted us to investigate the impact of the inhibitor on the kinase activity profile. We employed the ADP-Glo kinase activity assay to quantify the inhibitory potency of AZ191 (IC_50_) against DYRK1A and DYRK1B. In the assay, purified kinase domains of DYRK1A or DYRK1B (5 ng/µl) were titrated with the inhibitor (5 nM – 5 µM) in the presence of ATP (100 µM) and a synthetic peptide substrate (400 µM). The ATP concentration was chosen based on the experimentally determined K_M_ for DYRK1A and DYRK1B of 118 μM and 105 μM, respectively. AZ191 strongly inhibited DYRK1B and DYRK1A, with IC_50_ in the nanomolar range (Figure 2C). Notably, the inhibitory effect was more pronounced for DYRK1B, with an IC_50_ of 66 nM, whereas for DYRK1A, the IC_50_ was 188 nM. These results are consistent with previously reported data, IC_50_(DYRK1B) of 18 nM, IC_50_(DYRK1A) of199 nM and IC_50_(DYRK1B) of 83 nM, IC_50_(DYRK1A) of191 nM in radiometric assay and proximity assay, respectively^27^. For comparison, IC_50_ values were determined for harmine using the same assay conditions. Harmine inhibits DYRK1A and DYRK1B with IC_50_ of 97 and 115 nM, respectively.

The binding affinities of harmine and AZ191 to DYRK1A and DYRK1B were also evaluated by isothermal titration calorimetry (ITC) (Figure 2D). AZ191 demonstrated a binding affinity of 54 nM to DYRK1A and a stronger binding affinity of 28 nM to DYRK1B, suggesting a slight preference in binding to DYRK1B over DYRK1A. The study also indicated that harmine exhibited a binding affinity of 94 nM to DYRK1A and 29 nM to DYRK1B. This differential binding may reflect potential selectivity, which could be further explored for therapeutic implications, especially in designing selective inhibitors for DYRK1B.

The CETSA results confirmed the cellular interaction between DYRK1B and AZ191, as evidenced by the thermal stabilization of the protein. However, these results did not directly demonstrate the inhibition of DYRK1B kinase activity. To address this, we decided to investigate the effect of the AZ-191 inhibitor on the DYRK1B-controlled calcineurin/NFAT (Nuclear Factor of Activated T Cells) pathway. This signaling pathway is crucial in various biological processes, including the immune response and neuronal development. DYRK1B-mediated phosphorylation of NFAT prevents its translocation from the cytoplasm to the nucleus, thereby inhibiting gene transcription. The cellular localization of NFAT, which depends on its phosphorylation status, is a valuable indicator of DYRK1B kinase activity.

We utilized a cellular assay to track the translocation of EGFP–NFATc1 between the cytoplasm and the nucleus. In cells co-expressing mCherry-DYRK1B and EGFP–NFATc1, we observed predominant cytoplasmic localization of EGFP–NFATc1 (Figure 3A). This localization persisted even upon stimulation with ionomycin, a known activator of calcineurin, indicating that mCherry-DYRK1B effectively restrained the nuclear translocation of EGFP-NFATc1 (Figure 3B). However, upon treatment with 10 μM of AZ191, a marked translocation of EGFP–NFATc1 to the nucleus was observed despite the presence of mCherry-DYRK1B (Figure 3C). This demonstrates that a concentration of 10 μM of AZ191 inhibits DYRK1B activity and thereby restores the functionality of the calcineurin/NFAT pathway in our cellular model.

**Figure 3.**
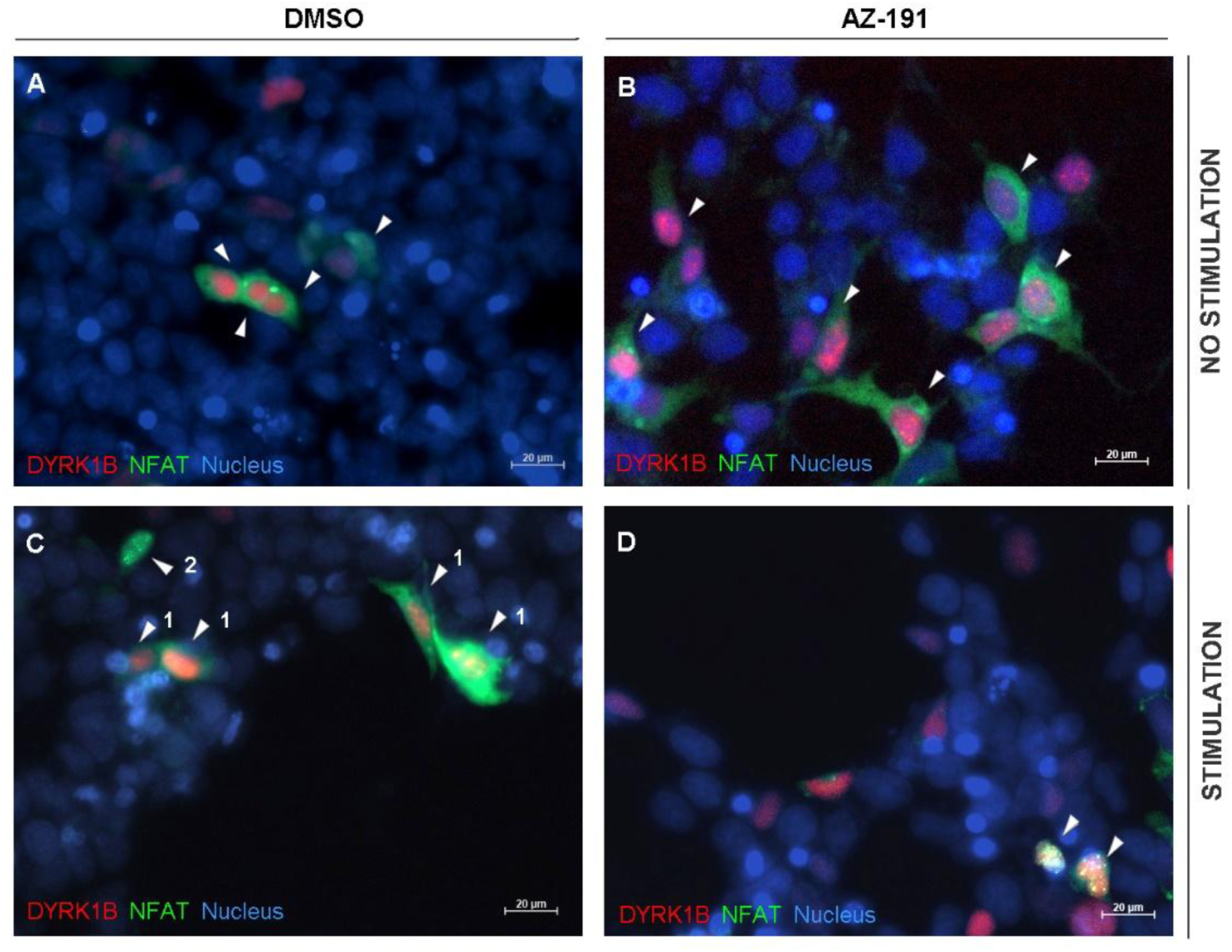
AZ191 treatment restores NFAT signalling by blocking DYRK1B kinase activity. The EGFP–NFATc1 (green) and mCherry-DYRK1B (red) were co-expressed in HEK293T cells. Cells pre-treated for 3 h with AZ191 (10 μM) (B,D) or DMSO (A,C) and stimulated for 1 h with ionomycin (IM; 6 μM) (C,D) or left untreated (A, B). The representative fluorescent images are shown with a scale bar 20 μm.

### Crystal structure of DYRK1B in complex with AZ191

Alexandrov et al.^29^ reported a pipeline for optimizing selective inhibitors for DYRK1B. However, the key challenge in this and other structure-based design approaches was the lack of an experimental structure for DYRK1B. Addressing this gap, we successfully expressed, purified, and crystallized the kinase domain of DYRK1B (residues 78 to 442; Uniprot ID Q9Y463) in complex with the inhibitor AZ191 (PDB ID 8C2Z). The kinase crystallized in the C222_1_ space group, featuring a single molecule per asymmetric unit. The structure, resolved at a 1.91 Å resolution, clearly delineates most of the molecule, except for portions of the CMGC kinase-specific insert (residues 382 - 394). Furthermore, residues 118 - 124 in the glycine-rich loop, residues 248 - 253, residues 266 - 273 within the activation loop and the regions encompassing additional parts of the CMGC insert (residues 357 - 369, 378 - 381 and 395 - 400) exhibit higher temperature factors compared to the rest of the molecule. This observation suggests that these areas possess structural flexibility, which could be crucial for understanding the kinase’s functional dynamics and designing more effective inhibitors.

The overall structure of DYRK1B resembles that of protein kinases of the CMGC group, with a bilobal fold and the active site located in a cleft formed between these lobes. The structure is seen in the active DFG-in conformation. The smaller N-terminal lobe mostly comprises β-sheets and contributes the catalytic lysine (Lys140). The larger C-terminal lobe is composed primarily of α-helices and contributes the activating tyrosine residue within the Tyr271-Xxx-Tyr273 motif. A clearly defined electron density indicates Tyr273 phosphorylation (Figure S6). Since the kinase was expressed in bacteria, where posttranslational modifications are relatively rare compared to eucaryotes, it is likely that the modification results from autophosphorylation. This is in good agreement with previous findings suggesting that DYRK1B undergoes autophosphorylation.^31^ Overall, the crystal structure of the DYRK1B kinase domain determined in this study overlays well with DYRK2 (PDB ID 4AZF) and DYRK3 (PDB ID 5Y86). The RMSDs are 0.85Å for 74.6% of C_α_ atoms and 0.77Å for 73.1% of C_α_ atoms for DYRK2 and DYRK3, demonstrating the high structural similarity.

AZ191 exemplifies an ATP mimetic kinase inhibitor and, accordingly, is located at the putative ATP binding site of DYRK1B (Figures 4A and B). The pyrimidine ring contributes a canonical interaction with the backbone amine of Leu193 within the hinge region. The methylpyrrolopyridine group penetrates deep into the pocket toward the catalytic Lys140. This moiety is stabilized by a direct hydrogen bond with the side chain aminium of Lys140, π-stacking interactions with the sidechains of the gatekeeper residues Phe190 and Phe122, and hydrophobic interactions contributed primarily by the sidechains of Val125 and Val258. The methylpiperazine moiety of AZ191, commonly found in several kinase inhibitors, generally protrudes out of the pocket into the solvent, which is also observed in our structure. However, what is characteristic of AZ191, however, is that the methylpiperazine contributes to affinity by forming a hydrogen bond with the sidechain carboxyl of Asp199. Additionally, this methylpiperazine contributes a water-mediated hydrogen bond with the main chain of Ile117. The methoxyphenyl linker contributes a π-lone pair interaction with the backbone carbonyl of Ser194, a hydrogen bond with Leu193, and hydrophobic interactions with the sidechain of Ile117. The methoxy substituent stabilizes the rotation of the moiety at the binding site.

**Figure 4.**
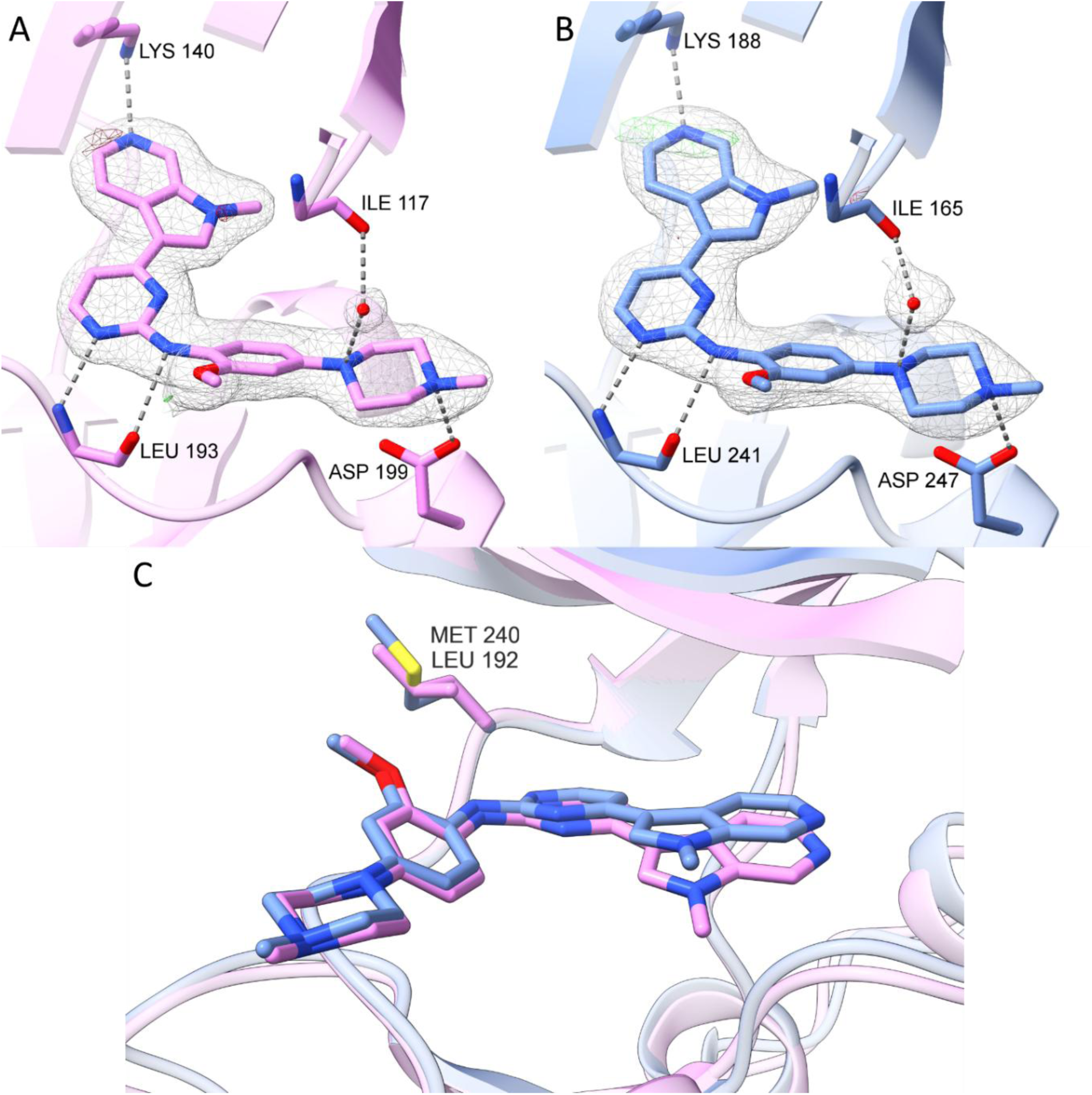
Crystal structures of DYRK1B (pink) and DYRK1A (blue) in complex with AZ191. (A) the binding mode of AZ191 in the ATP-binding pocket of DYRK1B (PDB ID 8C2Z), (B) binding mode of AZ191 in the ATP-binding pocket of DYRK1A (PDB ID 8C3G), (C) superimposition of DYRK1B and DYRK1A ATP-binding pockets with indicated single amino acid difference between their active sites. Hydrogen bonds are shown as dashed lines (dark grey) and the electron density is shown as a mesh, 2Fo-fc: +1.0σ (light grey); Fo-Fc omit map: +3.0σ (green); Fo-Fc omit map: −3.0σ (red).

Phosphorylation of Tyr273 provides an anchor point for a network of hydrogen bonds stabilizing the position of the Tyr273 sidechain at the surface of the kinase. These include salt bridges with the guanidine moieties contributed by Arg277 and Arg280. A water-mediated hydrogen bond with the sidechain of Ser276 further stabilizes the sidechain of Tyr273 and, thereby, the orientation of the Leu266-Ser276 linker. This region constitutes the peptide substrate binding site and is unstructured in related unphosphorylated kinases. Thus, phosphorylation regulates the activity by stabilizing the substrate binding cleft, as previously described for other kinases of the CMGC group.

### Crystal structure of DYRK1A in complex with AZ191

DYRK1B and DYRK1A kinases share a sequence identity of 85% among the catalytic domain residues. The identity is even higher for residues composing the active site. At such high evolutionary relatedness, the design of selective inhibitors is highly challenging, and success relies on subtle differences at the active site. A total of 83 crystal structures of DYRK1A in complex with inhibitors have been solved and reported. Collectively, this data allows us to compare our DYRK1B structure in search of specificity anchor points, but minor differences are best indicated when using structures with identical inhibitors. Consequently, we determined the structure of the DYRK1A kinase domain (residues 126-489, Uniprot ID Q13627) in complex with AZ191 (PDB ID 8C3G).

The DYRK1A in complex with AZ191 crystalized in the P2_1_2_1_2_1_ space group, and the asymmetric unit comprised 4 molecules. The crystals diffracted to 2.08 Å resolution. A well-defined electron density documents phosphorylation at the Tyr321 residue (Figure S6 B-E). Accordingly, the structure reflects that of a catalytically competent enzyme. Each kinase molecule in the asymmetric unit contains a clearly interpretable electron density describing the inhibitor (Figure S6 B-E). The protein molecules in the asymmetric unit are virtually identical, and the inhibitor dispositions at the active site of each molecule in the asymmetric unit do not differ. Thus, only model A is considered in further analysis.

The interactions guiding the inhibitor affinity are virtually identical to those observed in DYRK1B (Figure 4A and C). Hydrogen bonds contributed by the sidechains of the catalytic Lys188, Asp247, and the mainchain of Leu241 in the hinge region (equivalent to Lys140, Asp199, and Leu193 of DYRK1B, respectively) anchor AZ191 to the active site of DYRK1A. Further, the methylpiperazine moiety contributes a water-mediated hydrogen bond with the main chain of Ile165 (an equivalent interaction is found with Ile117 in DYRK1B). However, the said water is found only in molecules A and C. Further, the inhibitor is stabilized by major hydrophobic interactions contributed by Ile165, Phe170, Val173, Phe238, Leu294, and Val306 (equivalent to Ile117, Phe122, Val125, Phe190, Leu246 and Val258 of DYRK1B, respectively).

When comparing the AZ191 inhibitor in the co-crystal structure with DYRK1B to that with DYRK1A, different orientations of the methylpyrrolopyridine moiety are observed (Figure 5). In the structure of DYRK1B, the aromatic group is positioned closer to the C-lobe region. The difference, however, is not expected to be related to the fine details distinguishing the kinase active sites but rather reflects the different crystallization conditions, which, in the case of DYRK1B, further contained Mn^2+^ ions. In the DYRK1B structure, the manganese ion coordinates the side chains of Asp239 and Asp259, reorienting them away from the ATP binding pocket. This affords additional space for methylpyrrolopyridine reorientation when compared to DYRK1A. In the structure with DYRK1A, the absence of manganese ion allows the side chain of Asp307 (corresponding to the Asp259 in DYRK1B) to displace the methylpyrrolopyridine group of AZ191 towards the N-lobe. Apart from the influence of the ion, the binding modes of AZ191 at the active site of DYRK1B and DYRK1A are virtually identical, reflecting the comparable affinity of the inhibitor to both kinases (Figure 2C).

**Figure 5.**
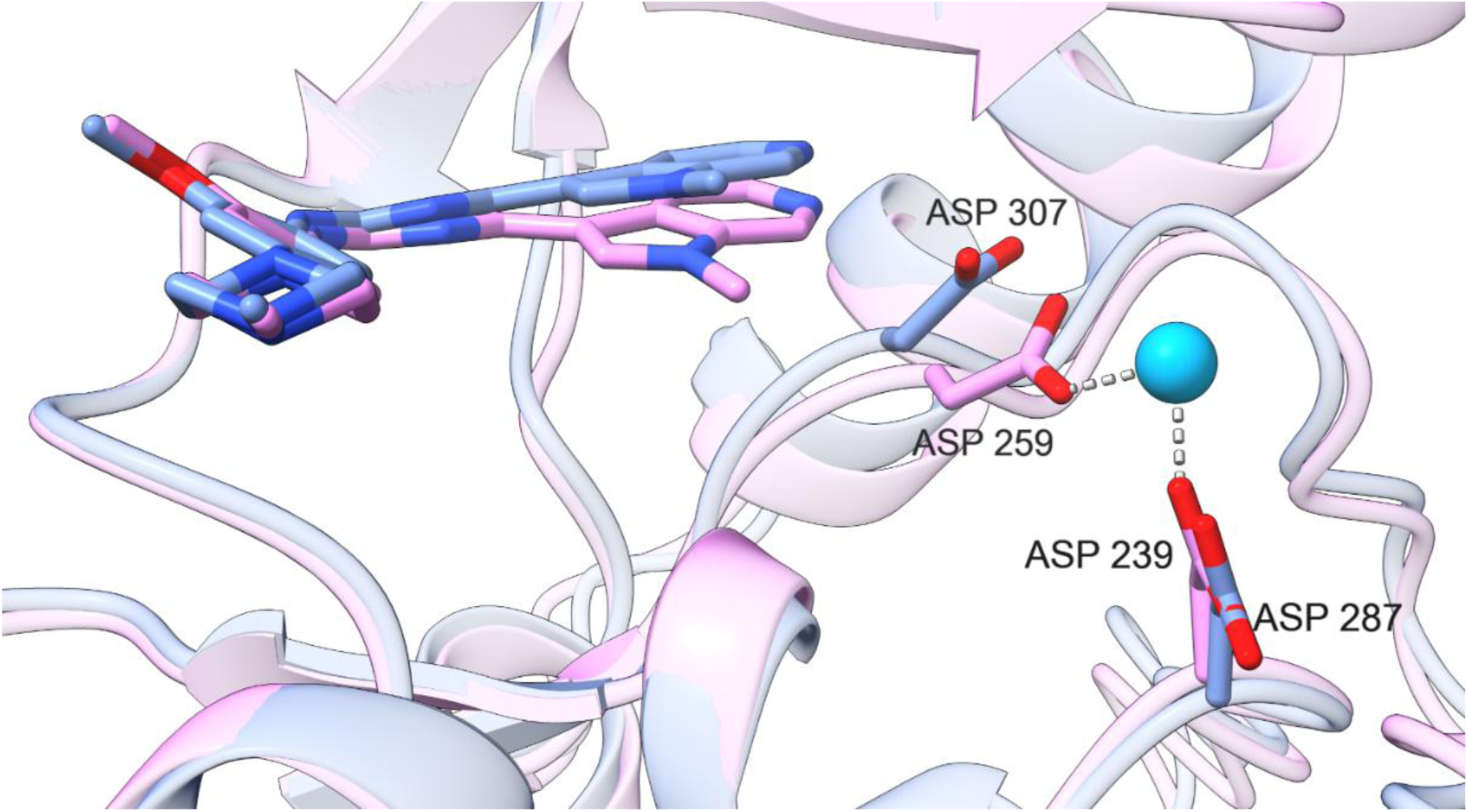
Superimposition of DYRK1B (pink) and DYRK1A (blue) in complex with AZ191 (PDB ID 8C2Z and 8C3G, respectively) with indicated Mn^2+^ ion bound to DYRK1B.

Despite the overall identity of the binding sites and inhibitor stabilizing interactions, the direct overlay of DYRK1B and DYRK1A crystal structures in complex with AZ191 points to a notable difference in the overall structures of the kinases. The active site of DYRK1B is narrower than that of DYRK1A, which is related to the closing of the N- and C-terminal lobes around the hinge region. The N-lobes of DYRK1B (residues 88-190) and DYRK1A (124-238) show a high degree of structural similarity, with RMSD of 0.34 Å for 75.7% of C_α_ atoms (Figure S5A). The C-lobes (residues 197-434 and 245-490 for DYRK1B and DYRK1A, respectively) also display a similar pattern, with RMSD of 0.39 Å for 72.3% of C_α_ atoms (Figure S5B). However, when comparing the overall structures of the kinase domains, the RMSD is 0.89 Å for 80.5% of Cα atoms. The increase in RMSD suggests that the N- and C-lobes are oriented differently relative to each other in the two kinases. To determine whether the differences in the relative orientation of the N- and C-lobes are characteristic of DYRK1B, we compared its structure against all available crystal structures of DYRK1A. From this structural evaluation, we concluded that this scissor-like relative relocation of the lobes around the hinge is characteristic of DYRK1A, and our structure with AZ191 represents one of the multiple relative orientations of the lobes. Other structures of DYRK1A present this reorientation of the lobes almost identical to our structure of DYRK1B (*e.g.*, PDB ID 2VX3; RMSD of 0.49 Å for 75.4% of C_α_ atoms). The apparent relative motions of the N- and C- lobes are partly related to crystal packing. The structure of DYRK1A closest to our present one in complex with AZ191 was determined in the same space group (*e.g.*, PDB ID 7FHT; RMSD of 0.30 Å for 89.8% of C_α_ atoms). On the other hand, the most distal structures with respect to the currently presented structure of DYRK1A with AZ191 were obtained in different space groups. Overlaying our crystal structure of DYRK1B with the most closely related structure of DYRK1A shows overall negligible differences.

Apart from the differences induced by crystal packing and crystallization conditions, the ATP binding pockets of DYRK1B and DYRK1A are practically indistinguishable, except for a single residue at the hinge region (Figure 4C). In the structure of DYRK1B, the sidechain of Leu192 is close to the methoxy group in the inhibitor, though it does not contribute to significant interactions. The equivalent residue in DYRK1A, Met240, has a slightly longer sidechain, which comes closer to the inhibitor. Convenient exploration and optimization of these interactions by modifications of substituents at the phenyl ring of AZ191 should allow for tackling the specificity of this class of inhibitors. Other scaffolds could also explore this pocket to achieve selectivity.

### Quantum Mechanical Binding Analysis

Despite the overall structural similarity and an almost exact identity in the binding pocket, DYRK1A and DYRK1B show different binding affinities (ΔG = RT log[K_d_]) and selectivity towards inhibitors. The binding affinities towards AZ191 differ by a factor of two (ΔΔG ∼ 0.41 kcal/mol). Differences in other thermodynamic data are more pronounced, like binding enthalpies, indicating subtle differences between the two kinases warrant further exploration. We employed our Energy Decomposition and Deconvolutional Analysis algorithm (EDDA) to better understand the underlying mechanisms behind such fine selectivity.^34^ EDDA is a quantum mechanical-based algorithm that partitions interaction energies into different contributions. These components include electrostatics, polarization, dispersion (lipophilicity), repulsion (shape complementarity), solvation, and charge transfer. The EDDA algorithm allows the determination of the nature of protein-ligand interactions, allowing the quantification of the main forces that lead to binding. For a detailed algorithm description, please refer to the Supporting Information (SI) section 2 and the original paper^34^. Figure 6 collects the electrostatics, dispersion (lipophilicity), repulsion, and total interaction maps for the inhibitor AZ191 in DYRK1A and DYRK1B. Figure 6 also contains a column plot of decomposed energies. Additional maps and the respective calculated values are given in supplemental information (Figures S7-S9 and Table S2).

**Figure 6.**
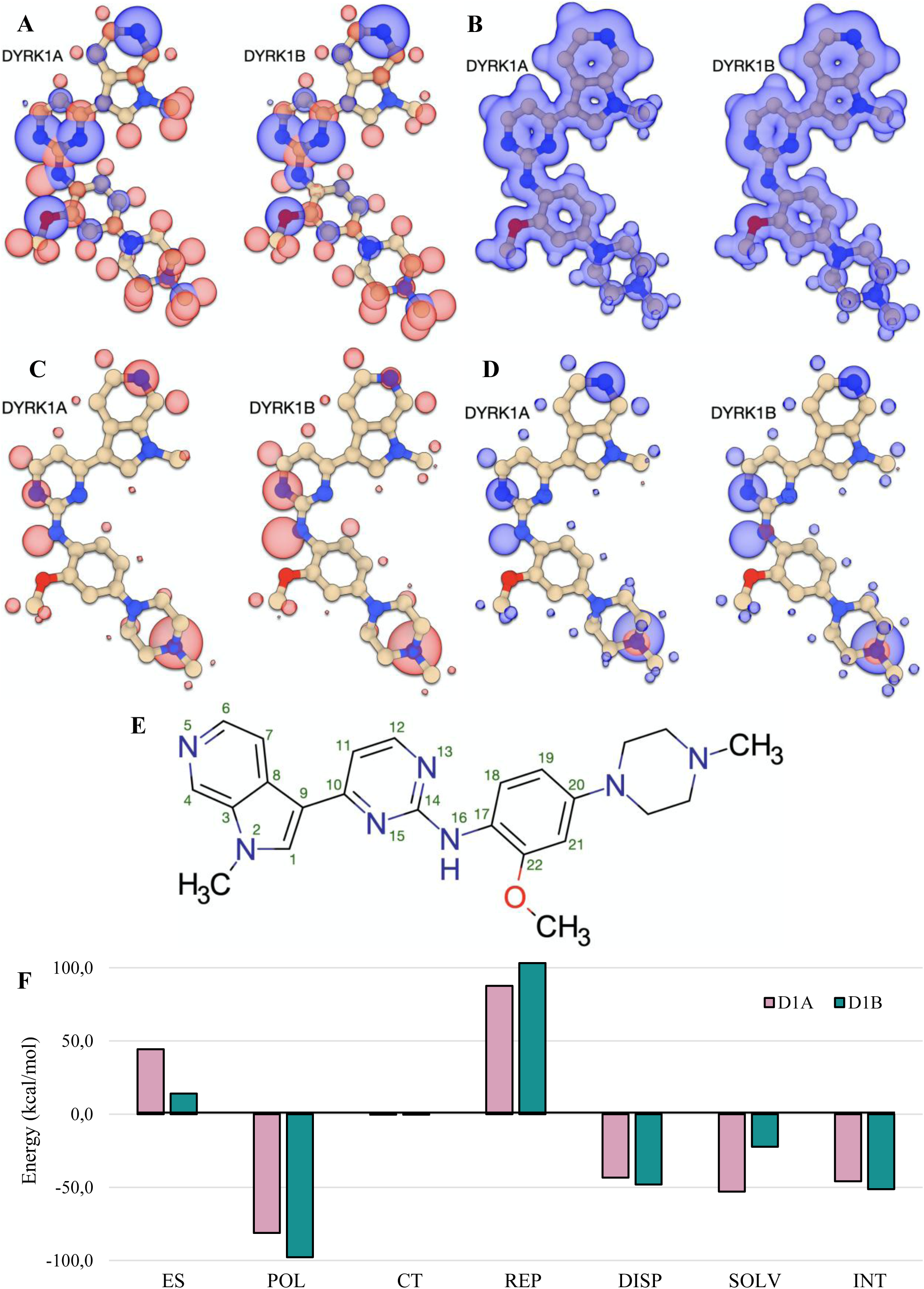
The decomposition analysis of the binding energy of AZ191 in DYRK1A and DYRK1B. (A) The electrostatics (ES), (B) dispersion (DISP), (C) repulsion (REP), and (D) total interaction imaps for AZ191 in the kinases. (E) The structure of the ligand and the atomic numbering used in the discussion. (F) Decomposed energies.

The initial observation when analysing the maps for AZ191 in each kinase is that the landscape of interactions is virtually indistinguishable between the two DYRKs analysed (Figure 6 A-D). This reflects the sequence homology around the ATP binding pocket. The ligand’s main foci of interactions are N5, which binds to the catalytic lysine; N15 and N16 atoms, which interact with the hinge region; and the external nitrogen from the piperazine moiety. Despite the similarity, the total interaction maps (Figure 6D) reveal stronger binding to the hinge of DYRK1B than to DYRK1A. The total interaction energies, comparable to enthalpies of binding, differ by 5.4 kcal/mol (in favour of DYRK1B), which correlates with the 5.0 kcal/mol measured by ITC. This establishes a strong agreement between our experimental and computational data. The electrostatics maps (Figure 6A) show more polar interactions in the case of DYRK1A, which, when combined with the repulsion and charge transfer maps, indicate a lower covalent character for the hydrogen bonds in this kinase. It is particularly interesting in the case of the linker N16, where the differences in maps are pronounced. Interestingly, the distance between N15 and N16 to the hinge binding atoms is shorter in DYRK1B by almost 0.2 Å for each bond. Repulsion maps (Figure 6C), which measure the overlap of electronic densities between ligand and protein, indicate stronger contacts with DYRK1B. This evidences that the higher covalent character in the hydrogen bonds between AZ191 and the kinase’s hinge is primarily an effect of atomic proximity and is less dependent on the protein’s charge distribution.

The interaction of N5 with the catalytic lysine is weaker in the case of DYRK1B. While not apparent from the viewpoint of electrostatics, this difference is evident in the repulsion and total interaction maps. Again, the effect seems to be primarily distance-related, as in DYRK1A, this hydrogen bond is shorter by almost 0.4 Å. Lastly, the interactions between the piperazine group and the proteins are identical.

To understand whether the different interactions with the hinge region and the catalytic lysine are significant from the binding perspective or whether these result from crystal artefacts (*e.g.*, lacking the dynamics of the protein-ligand complex), we analysed the total repulsion energy for each complex. Overall, repulsion is larger in DYRK1B by 15 kcal/mol (Figure 6F), indicating a stronger overlap of protein and ligand electronic densities. This difference can also be observed in the respective maps (Figure 6C), which show more intense contacts between DYRK1B and the ligand. The only exception is N5, the atom involved in interacting with the catalytic lysine, where the interaction is noticeably stronger in DYRK1A. Overall, the calculations indicate that AZ191 captures more effectively DYRK1B by the hinge region, whereas in DYRK1A, the interactions with the hinge and the catalytic lysine are more balanced. Indirectly, this hints that the accessibility of the catalytic lysine of DYRK1B is lower and could be a useful feature for achieving selectivity.

### A Role for the CMGC Insert

The CMGC insert is one of the most variable regions between DYRK1A and DYRK1B, with a sequence identity of 66% (comparing to 85% for kinase domains). However, only one of these differences alters the amino acid’s charge state. Since the CMGC region was excluded from previous calculations, we reran our EDDA analysis using the full, experimentally determined kinase constructs.

Figure 7A compares the DYRK1A-DYRK1B EDDAs for two system reconstructions: the large kinase domains analyzed in the previous section versus the full kinase domains obtained from X-ray crystallography experiments. The total relative binding energies remain similar, further reinforcing the robustness and convergence of the previously calculated values. However, the absolute binding energy values vary due to changes in the protein’s total charge. Notably, in the full constructs, the protein component of the complex becomes more positively charged, leading to an overall weakening of the total interaction. This effect is furthermore strongly reflected in the relative electrostatic and solvation contributions, as shown in Figure XA. Specifically, the electrostatic repulsion between DYRK1A and AZ191 increases when considering the full kinase domain, including the CMGC insert. In contrast, electrostatic interactions between DYRK1B and AZ191 become predominantly attractive, indicating an inversion of the electrostatic interaction pattern in the case of DYRK1B. Interestingly, the solvation contributions to binding in DYRK1A remain unchanged when using the full protein system in the calculations. On the other hand, for DYRK1B, the solvation terms become repulsive. These findings align with our previous observations regarding electrostatics. While differences exist in other binding forces (CT-POL, REP, DISP), they are less significant than the effects of electrostatic interactions and solvation.

**Figure 7.**
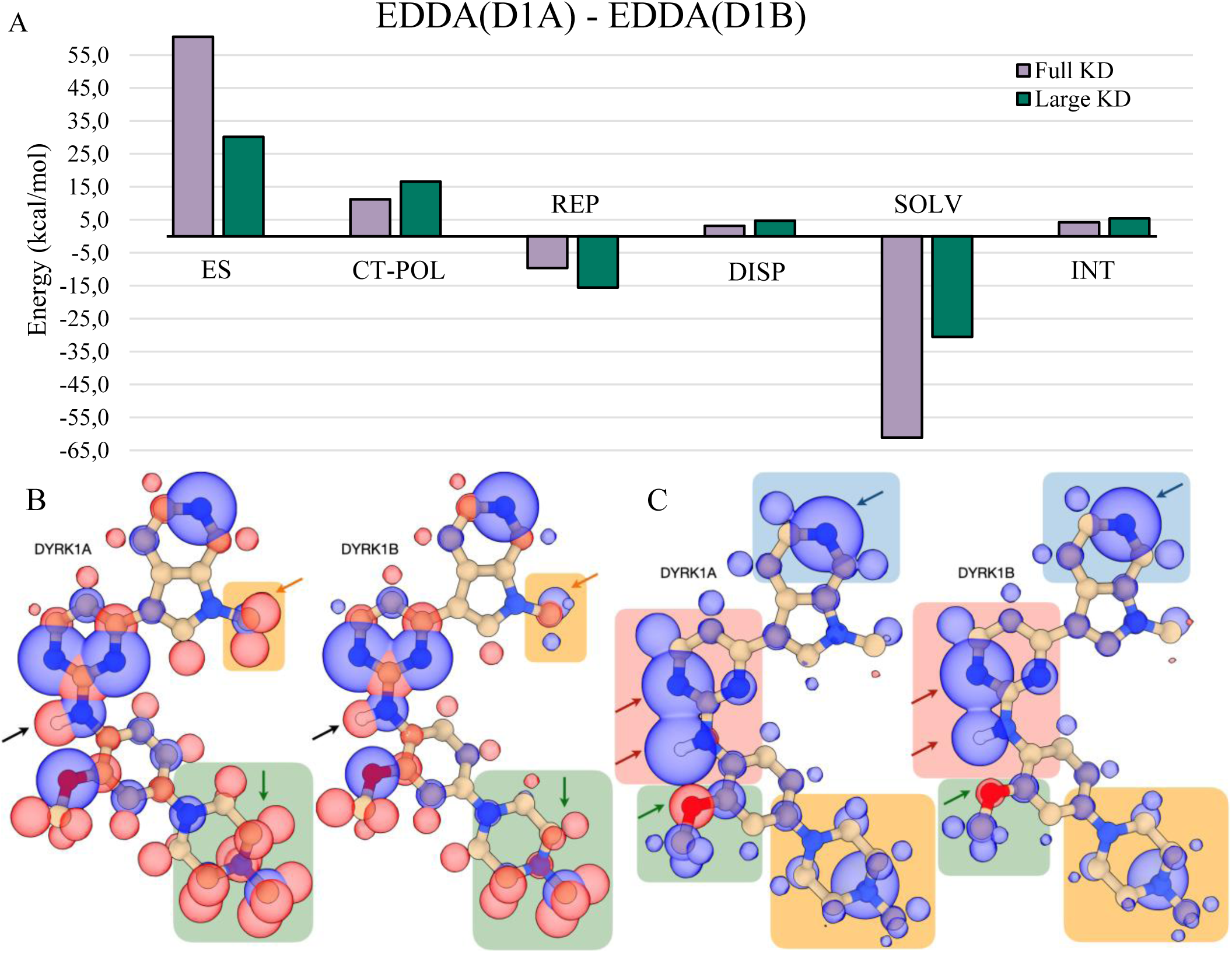
The binding energy decomposition analysis of AZ191 in the full kinase domains of DYRK1A and DYRK1B, compared against the large pocket previously analysed in Figure 6. We coupled the POL and CT interactions because the underlying algorithm needed to split these terms has poor convergence properties with system size. A) The differential EDDA plot, where each interaction contribution in the DYRK1B-AZ191 complex is subtracted from the respective contribution in the DYRK1A-AZ191 EDDA. KD in the legend stands for Kinase Domain. B) The ES maps for AZ191 in each of the DYRK1 kinases using the full kinase domains. The main regions where the maps differ are highlighted with yellow and green backgrounds. C) The total interaction maps (INT) for AZ191 in each of the DYRK1 kinases, using the full kinase domain. The main regions where the maps differ are highlighted with yellow, green, blue, and red backgrounds.

At first glance, the EDDA interaction maps calculated with the full kinase domains show no significant changes in interaction patterns (Figure 7B and 7C), with the overall map composition largely conserved compared to our previous calculations (Figure 6A and 6D). This shows that the first amino acid shells are the primary drivers of protein-ligand interactions and that expanding from the large kinase domains used in the previous section to the full kinase domains in the new calculations only incorporates long-range interactions. However, we do observe differences in the relative contributions of ligand functional groups involved in non-specific and non-directional interactions. These may influence kinase selectivity based on ligand electronic properties. For example, the electrostatic interaction pattern differs around AZ191’s methyl-pyrrolo-pyridine moiety (yellow highlighted region in Figure 7B). The increased repulsive interactions in the DYRK1A complex suggest that modifying the electronic properties of this exposed functional group could provide valuable structure-activity relationship (SAR) insights. Similarly, our calculations indicate a comparable effect for AZ191’s piperazine group (green highlighted region in Figure 7B). Although this piperazine moiety is in proximity to an Asp residue (Asp247 in DYRK1A, Asp199 in DYRK1B), it does not fully disrupt the interaction between this Asp’s carboxylate and a nearby Arg residue (Arg250 in DYRK1A, Arg202 in DYRK1B). Similar observations are made when analyzing the total interaction maps (INT) around this group (Figure 7C). The INT maps highlight additional ligand modifications that could be of interest for SAR exploration. AZ191’s interactions with DYRK1A’s catalytic lysine are strengthened compared to those with DYRK1B (blue highlighted region). Differences are also observed in the interaction patterns around the hinge region of each kinase (red highlighted region), suggesting that variations in the hydrogen-bonding properties of hinge-binding groups may yield valuable SAR insights. This was also reflected at the ES-interaction level, as seen by the black arrows in Figure 7B. A closer look at the maps indicates that, energetically, the interaction with the hinge has a stronger bidentate character for DYRK1A. Potentially, designing mono-dentate hinge binders will have less impact for DYRK1B inhibiton than it does for DYRK1A. Similar observations were already noted in the previous section’s analysis. A comparable trend is seen for the methoxy substituent on the aromatic ring linking the hinge-binding motif to the solvent-exposed piperazine. While our analysis is based on only two structures, limiting its reliability, we believe that further exploration of the interaction imbalance at the kinase hinge and the accessibility of the catalytic lysine could guide the development of the first DYRK1-selective ligands.

### The Dynamics of DYRK1A and DYRK1B

To better understand the behaviour of DYRK1A and DYRK1B, apo and inhibitor complexes, a series of molecular dynamics (MD) simulations were conducted. MD and EDDA results on the MD traces are presented in Figure 8 (see supplemental Tables S3-S6 for additional data).

**Figure 8.**
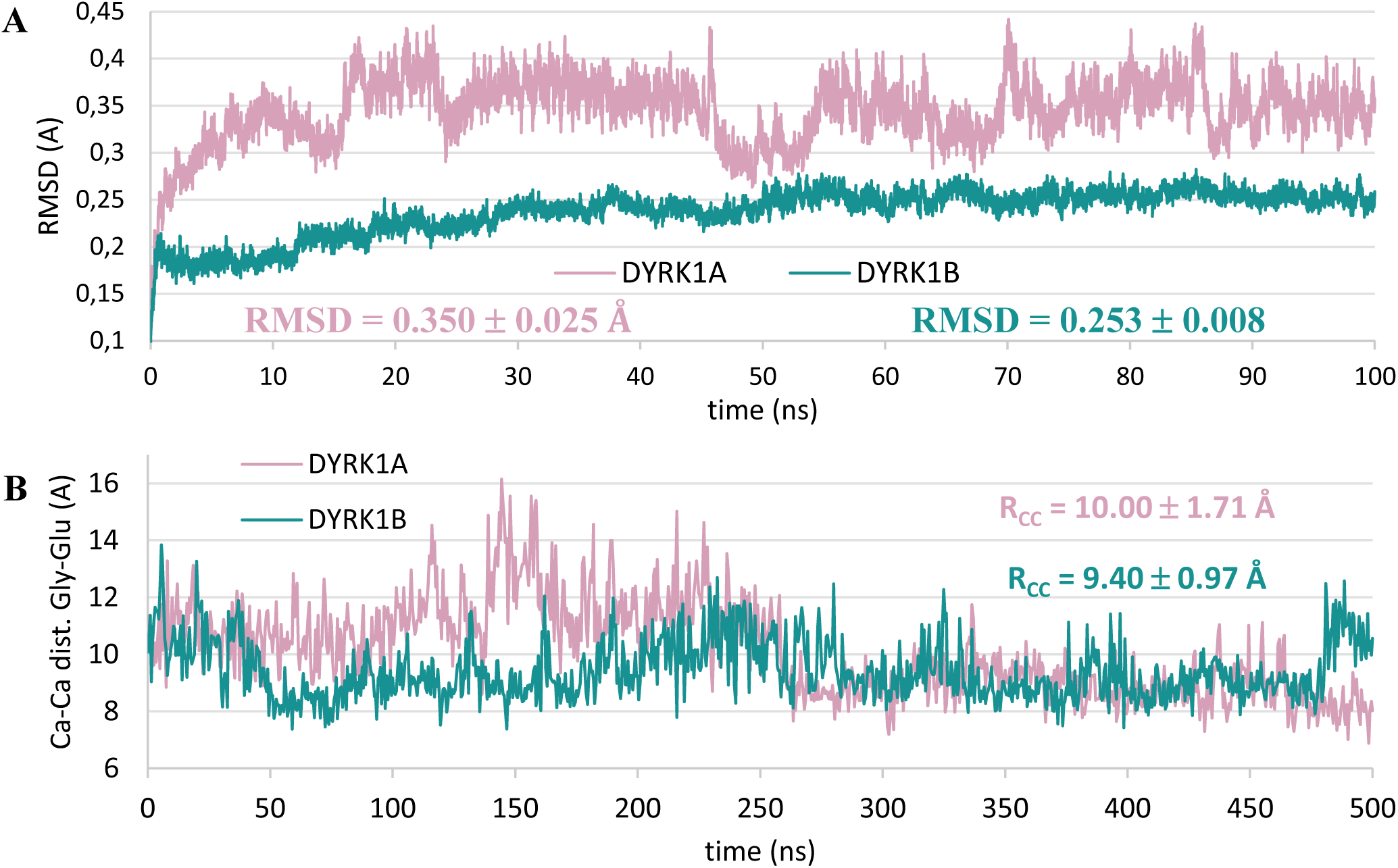
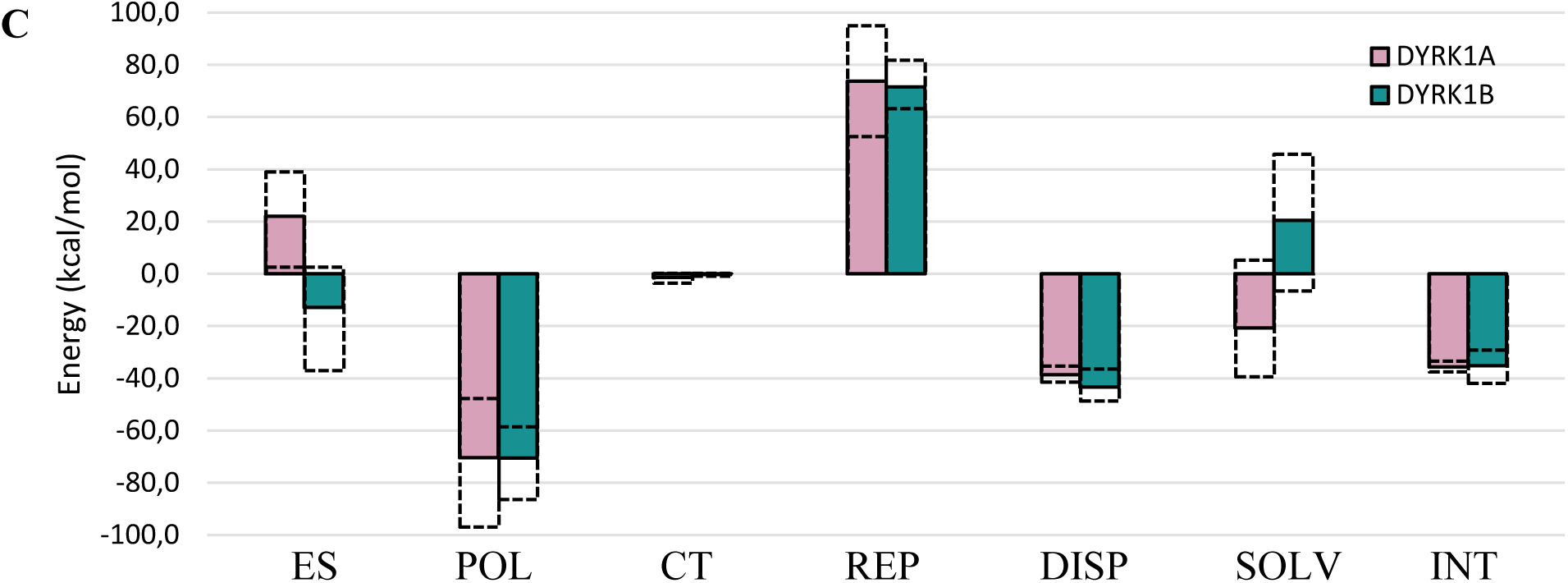
The dynamics of the apo and AZ191 bound DYRKs. (A) RMSD of pocket residues as a function of time. Average RMSDs and the respective standard deviations that are given in the plot were obtained for the last 40 ns of simulation time. (B) The distance between the Cα in the Gly-Glu residues of each DYRK (for DYRK1A: Gly168-Glu291; for DYRK1B: Gly120-Glu243) as a function of time. Average distance (and standard deviation) calculated after the first 100 ns of simulation time. (C) Time-averaged EDDA on DYRK1A and DYRK1B with AZ191 and the respective limits observed for each contribution. EDDA conducted on the larger subsystem B.

The most direct observation from the apoprotein simulations is the larger flexibility of DYRK1A, as seen in the RMSD fluctuations of the dynamical traces (Figure 8A). On average, the N- and C-lobes are more distal in DYRK1A, with wider amplitude movements (Figure 8B). This is, however, a simulation artefact due to the extended time required for DYRK1A to relax from the initial structure with the ligand’s imprint. After about 250 ns, DYRK1A closes the pocket, remaining in this state until the end of the simulation. This we identify as the resting state of the kinase, which is identical for both paralogs (Figure S11).

Figure 8C shows the time-averaged EDDA on both protein-ligand complexes. Interestingly, all interactions decrease in magnitude compared to crystal structure calculations. Most contradictory, the average shape complementarity becomes larger in DYRK1A. The average binding energy also favours DYRK1A, which contradicts experimental data. Though DYRK1B shows larger variations in binding energy, the EDDA binding energy as a function of time tendentially decreases for DYRK1A and increases for DYRK1B. These trends and the simulation snapshots indicate the inability of AZ191 to reach DYRK1B’s catalytic lysine. This is reflected in the distances between the ligand’s N5 and the NZ of the catalytic lysine (Figure S12): on average, 3.17 Å for DYRK1A and 3.51 Å for DYRK1B, differ significantly from the experimental values of 2.92 and 2.97 Å, respectively. Though this imbalance warrants caution when analysing MD results, we believe it underscores the previously noted differences in the accessibility of the catalytic lysine. This raises the question: how can two kinases that show such remarkable sequence and structural similarities exhibit such disparate behaviour on their ATP-binding pockets, allowing biological systems to selectively exploit kinase activity? Answer to this question, the long-range effect of distal residues, is corroborated by calculations (Table S7): as the number of protein atoms included in calculations increases, electrostatics become less repulsive. The impact of distal residues extends, however, to other interactions. The repulsion energy between the inhibitor and the protein increases with the pocket size. By including larger protein domains in the calculations, fine changes in the charge distribution and electronic populations occur, affecting how the two molecules interact. Note that because of the exponential decay of repulsion with distance, the 9 kcal/mol difference in repulsion implies only punctual modifications to the electronic densities. Overall, the calculations reveal a clear atomic mechanism underlying selectivity between the two kinases, which could only be uncovered by incorporating detailed electronic effects into the analysis.

## Discussion

DYRK1A’s involvement in neurological disorders and diabetes has been well-documented, with significant progress made in inhibitor design and development. In contrast, DYRK1B has only recently been recognized as a potential target in cancer and non-alcoholic fatty liver disease.^35^ The challenge in targeting DYRK1B lies in the poor selectivity of existing probes against related CMGC group kinases and the near absence of selectivity among DYRK inhibitors. AZ191 inhibits over 10 other kinases^28^. While some researchers advocate for selective inhibition of DYRK kinases to mitigate toxicity, concrete evidence supporting this claim is limited.

For example, currently available DYRK1A and 1B inhibitors also affect Homeodomain-Interacting Protein Kinases (HIPKs), crucial for eye, wing, nervous system, and muscle development in *Drosophila melanogaster*.^36,37^ The implications of HIPK inhibition in mammals are yet to be fully understood, but the findings in Drosophila suggest the need for thorough testing. Other possible off-targets are Cdc-like kinases (CLKs), with CLK1 involved in cell cycle-related splicing and CLK2 disruptions causing cell cycle defects and proliferation loss.^38,39^ Therefore, monitoring the long-term effects of DYRK1A/1B inhibitors is essential. Selective inhibition of DYRK kinases over related CMGC kinases may offer higher efficacy and help define regulated substrates and downstream effects.

Selectively targeting specific DYRK kinases is also advantageous. DYRK1B activity is implicated in certain cancers and non-alcoholic fatty liver disease. DYRK1B upregulation in ovarian, pancreatic, and non-small cell lung cancers and its inhibition in certain ovarian cancer cell lines leads to increased ROS levels and apoptosis. In contrast, similar treatment in colon cancer cells expressing only DYRK1A showed no effect.^30^ Elevated DYRK1B levels in non-alcoholic steatohepatitis (NASH) patients correlate with hypertriglyceridemia, fatty liver, and hepatic insulin resistance.^35^ However, concurrent inhibition of DYRK1A is undesirable due to the risk of elevated homocysteine levels known for their neurotoxic effects^40^, as demonstrated in mice.^41^ These findings underscore the need for *in vivo* testing of the benefits of selective DYRK kinase inhibition.

Our structural characterization of DYRK1B, alongside the earlier DYRK1A structures, sets the stage for the rational design of the inhibition module. The enzyme-inhibitor interactions follow Koshland’s induced fit model.^42^ Thus, to minimize the influence of inhibitor variability in our comparative analysis, we also determined the structure of DYRK1A in complex with the same inhibitor, AZ191. Following the high sequence homology, the ATP pockets of DYRK1A and DYRK1B are almost identical, with only a single residue differentiating the two. This site, addressed by the methoxy moiety of AZ191, suggests a convenient mechanism for enhancing specificity. Bulkier substituents are expected to cause a steric clash in DYRK1A due to the ramification in the leucine sidechain, a hypothesis that remains to be experimentally verified. Thus, optimizing interactions at the Leu/Met site appears to be one of the rational approaches for enhancing the selectivity of small molecule ATP mimetic inhibitors based on AZ191 or other scaffolds. The quantum mechanical calculations revealed two additional factors that could be investigated to explore selectivity. Firstly, the ATP-binding pocket’s electrostatic environment, influenced by distal residues, differs significantly between the kinases. However, leveraging electrostatics for selectivity is complex due to environmental factors like salt concentrations, which vary between *in vitro* and *in vivo* conditions. Secondly, simulations showed reduced accessibility to the catalytic lysine in DYRK1B, defined by a narrower ATP-binding pocket. This suggests a more stable and environment-independent approach for inhibitor design. Our model hypothesises that DYRK1B’s restricted access to the catalytic lysine renders binding to this kinase less sensitive to the bulkiness of the lysine-binding groups. On the other hand, binding to DYRK1A should favour smaller and more compact lysine binding groups. To test this, we examined published data on ligands structurally similar to AZ191, such as Abemaciclib and Cirtuvivint.^27^ The interaction with the catalytic lysine is key to both the inhibitors and involves a benzimidazole ring in Abemaciclib and a pyrazole ring in Cirtuvivint. Abemaciclib, which contains a bulky isopropyl group in its benzimidazole ring, shows a 5-fold higher affinity for DYRK1B than DYRK1A. Conversely, Cirtuvivint, with a smaller methyl group near the compact pyrazole ring, exhibits a 5-fold lower affinity for DYRK1B than DYRK1A. Additional evidence comes from ARN25067 and TCMDC-135051,^27^ both with bulky substituents near the lysine-binding groups, resulting in a preference for DYRK1B (factor of 8 for ARN25067 and factor of 2 for TCMDC-135051). Similarly, compounds SM77 and SM330^27^ lack bulky substituents and bind more strongly to DYRK1A (factor of 2-3). EHT-1610, with its non-bulky methyl acetimidate group, also shows stronger binding to DYRK1A by a factor of 3. Our model also explains the slight preference of AZ191 for DYRK1B, given its indole ring’s size in capturing the catalytic lysine.

In conclusion, our crystal structure of DYRK1B kinase opens the door for developing selective inhibitors. The almost identical ATP pockets of DYRK1A and DYRK1B underscore the challenge of this endeavour, but the Leu/Met site and the accessibility of the catalytic lysin offer potential anchor points. Future development of selective inhibitors will enable assessing whether selective targeting is more beneficial than the current inhibitors targeting both kinases with similar potency.

## Experimental Section

### Compound Purity

All compounds are >95% pure by HPLC. Harmine, acquired from Sigma-Aldrich, catalog number 286044, purity of 98%; AZ191, acquired from MedChemExpress, catalog number HY-12277, purity of 99.87%.

### Plasmid construction

The gene encoding the kinase domain of DYRK1B (78-442; Uniprot ID Q13627) was codon optimized, synthesized by Genscript and cloned into a pET24a plasmid. The kinase domain of DYRK1B was expressed with a C-terminal tobacco etch virus (TEV) protease cleavage site followed by His-Tag (ENLYFQ*GHHHHHH).

The gene fragment encoding the kinase domain of DYRK1A (126-490; Uniprot ID Q9Y463) was PCR amplified from plasmid pEXP17-DYRK1A^43^ and subcloned into pET24a plasmid. The kinase domain of DYRK1A was expressed along with a non-cleavable C-terminal His-Tag.

### Protein expression and purification

DYRK1B was expressed in *E. coli* LOBSTR strain (Kerafast) in TB medium supplemented with kanamycin (50 µg/mL) at 17 °C. 16 hours after the induction the cells were harvested by centrifugation and the pellet was resuspended in lysis buffer (20 mM HEPES pH 7.5, 300 mM NaCl, 5% glycerol, 15 mM imidazole, 5 mM MgCl_2_, and 5 mM 2-mercaptoethanol) and lysed by sonication. Clarified lysate was incubated with HisPur™ Cobalt resin (ThermoFisher Scientific, Waltham, MA, United States) for 1 hour at 4 °C, and DYRK1B was eluted with increments of imidazole concentration (50–300 mM). The elution fractions were pulled and dialyzed against 20 mM HEPES pH 7.5 containing 150 mM NaCl, 5mM MgCl_2_, 5 mM 2-mercaptoethanol and TEV protease. Released His-tag was removed by reverse chromatography on HisPur™ Cobalt resin. Final purification was obtained on HiLoad 16/600 Superdex 75 pg column (Cytiva) in 20 mM HEPES pH 7.5 containing 150 mM NaCl, 5 mM MgCl_2_ and 1 mM TCEP.

The expression and initial purification steps for DYRK1A were identical to DYRK1B. Following cobalt affinity chromatography, the kinase was dialysed into 20 mM HEPES pH 7.5 containing 50 mM NaCl, 5 mM MgCl_2_ and 5 mM 2-mercaptoethanol. Further purification was obtained by ion-exchange chromatography on HiTrap Q FF column (Cytiva) followed by size exclusion chromatography as for DYRK1B.

Purified proteins were flash-frozen in liquid nitrogen and stored at −80 °C for further analysis.

### Dye-based Thermal Shift Assay

DYRK1A and DYRK1B kinase domain stability in the presence of harmine and AZ191 were analysed by the proteins’ melting temperatures determination using Thermal Shift Assay (TSA) as described previously.^44^ Both proteins (1.5 mg/ml) were incubated with 1:200 diluted Sypro Orange dye 20 mM HEPES pH 8.0, 100 mM KCl, 10 mM MgCl_2_, 1 mM 2-Mercaptoethanol and compound (10 μM) or DMSO. The fluorescence signal of Sypro Orange was determined as a function of temperature between 5 and 95 °C in increments of 0.5 °C min^-1^ (λ_ex_ 492 nm, λ_em_ 610 nm). The melting temperature was calculated as the inflection point of the fluorescence as a function of temperature. The experiment was carried out in triplicates repeated two times with similar results.

### Cellular Thermal Shift Assay (CETSA)

The interaction between AZ191 and full-length DYRK1B kinase was analyzed by cellular thermal shift assay in complex cell lysate. Samples were prepared from control and inhibitor-exposed cell lysates. For each set, 2 × 10^7^ cells were seeded in a 10-cm cultured dish. After 24 h of culturing, the cells were transfected with plasmid encoding Flag-DYRK1B using PEI Prime (Sigma-Aldrich), then 4 h later medium was changed and cells were left for 48 h. Then cells were treated with AZ191 (10 µM) or DMSO. 24 h later the cells were washed with PBS, treated with trypsin and collected by centrifugation at 1200 rpm for 5 min at room temperature. The pellets were gently resuspended in 800 µl of RIPA lysis buffer (Sigma-Aldrich) supplemented with Protease Inhibitor Cocktail (Roche) and disintegrated by sonication. The samples were centrifuged at 20 000 rpm for 30 min at 4 °C, and the supernatants were transferred to new Eppendorf tubes. Pairs consisting of one control aliquot and one experimental aliquot were heated at 40 °C, 45 °C, 50 °C, 55 °C, 60 °C, 65 °C or 70 °C for 5 min. Next the samples were cooled on ice, centrifuged at 20 000 rpm for 30 min at 4 °C and supernatants were analyzed by Western Blot using monoclonal ANTI-FLAG M2-Peroxidase (HRP) antibody (Sigma-Aldrich, A8592).

### ADP-Glo kinase activity assay

The inhibitory potency (IC_50_) of harmine and AZ191 was determined with the ADP-Glo kinase activity assay (Promega). The synthetic peptide DYRKtide (RRRFRPASPLRGPPK, Caslo ApS) was used as a substrate for DYRK1A and DYRK1B. 20 µl of kinase in reaction buffer (100 mM MOPS pH 6.8, 100 mM KCl, 10 mM MgCl_2_) and 10 µl of compound were incubated for 15 minutes at room temperature. The reaction was started by simultaneous addition of 20 µl of substrates mix (ATP and DYRKtide) and continued for 30 minutes at room temperature. The 5 µl of reaction mixture was transformed to HTRF 96 well low volume plates (Cisbio) and the consumption of ATP was measured with ADP-Glo Kinase Assay kit (Promega Corporation) according to manufacturer protocol. The final composition of reaction reagents was - kinase (DYRK1A, DYRK1B) 5 ng/µl, compound (harmine, AZ191) 5 nM – 5 µM, ATP 100 µM and DYRKtide 400 µM. All measurements were done in triplicate and repeated two times. The IC_50_ was determined using GraphPad Prism software.

### Isothermal titration calorimetry measurements

ITC was performed with Nano ITC 2G (TA Instruments). To measure the DYRK1A interaction with harmine and AZ191, the proteins were dialyzed overnight at 4 °C against the ITC buffer (20 mM HEPES, 150 mM NaCl, 1 mM TCEP and 0.2% DMSO). The reaction cell was filled with a respective protein solution (10 μM) whereas harmine or AZ191 was prepared in the same ITC buffer at 100 μM (from 50 mM stock solution) and loaded into a dosing syringe. All samples were degassed using an accessory degassing system (TA Instruments) prior to being placed in the ITC to minimize the possibility of gas bubble formation during the run by pulling a vacuum of 0.3–0.5 atm for 10–15 min. Each injection (5 μl) was performed in 300 s intervals. All experiments were conducted at 25 °C with a stirring rate of 250 rpm. The results were analyzed using NanoAnalyze software (TA instruments). The experiment was carried out once.

### NFAT translocation assay

HEK293T cells were grown on μ-Slide 8 well (IBIDI) to 50%–70% confluency. The plasmids expressing mCherry-DYRK1B and eGFP-NFATc1 were transiently co-transfected for 48 h with PEI Prime (Sigma-Aldrich). Cells were pre-treated with inhibitor AZ191 (10 μM) or DMSO for 3 h and then stimulated with ionomycin (Thermo Fisher Scientific) for 1 h. Cells were washed with 1 ml PBS and the nuclei were stained with Hoechst 33258 (ThermoScientific) for 10 min at 37 °C and fixed with 4% paraformaldehyde in phosphate-buffered saline (PBS) for 10 min at 25 °C. Images were collected with Zeiss Axio Observer 3 fluorescence microscope and analysed in ZEN Blue edition software.

### Mass Spectrometry

Samples for MS were taken during first step of protein purification (after sonication) and final purified protein. Bottom-up protein identification with phospho-modified residues was done according to the typically used protocols. Briefly, sample in the form of Coomassie Brillant Blue stained band was excised and destained (ammonia-bicarbonate/AMBIC buffer 50 mM, pH=8.0/acetonitrile). Then the band was rehydrated in the AMBIC buffer with following routine reduction (5 mM DTT, 10 min, 80°C, with shaking) and cysteine alkylation (5 mM iodoacetamide, 10 min, 80°C, with shaking) procedure. Peptides were eluted from the gel, using water/acetonitrile/formic acid mixture (9/10/1 v/v/v), then freeze-dried and dissolved in the 60 μl of 4% acetontrile/water solution supplemented with 0.1% formic acid. 5 μl were introduced into the nanoLC-MS/MS system. Equipment used: nanoLC: Ultimate 3000 (Thermo, Bremen, Germany) in the set-up containing trap cartridge (Acclaim Pepmap 100, C18, 1 mm ID, 5 mm length, particle size 5 μm) and analytical column (Acclaim Pepmap 100, C18, 75 μm ID, 150 mm length, particle size 3 μm). Mass spectrometer: Exploris 240 (Thermo, Bremen, Germany) equipped with Nanopray FLEX ion source. Peptides separation was done using following solvents: A: water+0.1% formic acid, B: acetonitrile + 0.1% formic acid. Gradient settings: T=0 min 5%B, T=5 min 5%B, T=20 min 35%B, T=22 min 75%B, T=23 min 75%B, T=25 min 5%B, T=30 min 5%B. Mass spectrometer settings: MS spectrum range: 350-1600 m/z, resolution 30000, MS/MS spectrum range: 200-2000 m/z, resolution 60000. Up to 6 fragmentation scans between MS scans. Ion exclusion for 20 sec after 2 MS/MS scans.

Data analysis was done as follows: conversion from raw to mgf files was done with the aid of RawConverter software, ver. 1.1.0.23 64-bit (from: http://fields.scripps.edu/rawconv/). Mgf files were searched against the database containing the single sequence of the protein-of-interest (DYR1B_HUMAN). Search engine was the local Mascot server, ver. 2.8.0 (Matrixscience, UK). Mascot settings were as follows: enzyme: trypsin, up to 2 missed cleavages; fixed modifications: carbamidomethylation; variable modifications: Met-oxidation, phospho (ST), phospho (Y); MS and MS/MS tolerances: 100 ppm; #13C = 1; peptide charge: 2+, 3+ and 4+; instrument: ESI-FTICR.

### Protein crystallization, data collection and structure determination

DYRK1B was concentrated to 10-13 mg/ml and incubated overnight with 5-molar excess of AZ191 at 4 °C. For crystallization condition screening the preparation was mixed 1:1 (v/v) with crystallization solutions. The experiments were carried out at 20 °C and crystals appeared within 1-3 days. The best crystals were obtained using sitting drop vapor diffusion setup in 0.1 M Bis-Tris pH 5.5 containing 25% PEG 3350, 0.2 M magnesium chloride and 0.1 M manganese chloride. Crystals of DYRK1A in complex with AZ191 were obtained in a manner similar to those of DYRK1B, but from 0.1 M HEPES pH 7.5 containing 10% PEG 8000, 8% ethylene glycol and 0.1 M ammonium sulfate. Crystals were cryoprotected in mother liquor containing 25% ethylene glycol and flash-frozen in liquid nitrogen.

X-ray diffraction data were collected at BESSY (Berlin) and DESY (Hamburg). The diffraction data was indexed and integrated using XDS.^45^ Data was scaled in AIMLESS^46^ from the CCP4 software package.^47^ Following steps were performed in Phenix.^48^ The structures of DYRK1B and DYRK1A were solved by molecular replacement using PHASER^49^ and respective search models: processed AlphaFold model AF-Q9Y463-F1 and PDB ID 7A4O. Models were refined by interchanging cycles of automated refinement using phenix.refine^50^ and manual building in Coot.^51^ Restraints for the inhibitor were created in Grade Web Server.^52^

### Computational Methods

Energy Decomposition and Deconvolutional Analysis (EDDA) calculations were performed using our in-house quantum chemical package, ULYSSES.^53^ Details of the EDDA method were published earlier.^34,54^ The quantum chemical method of choice was GFN2-xTB^55^ with ALPB aqueous solvation.^56^ Molecular Dynamics simulations were conducted using GROMACS^57^ with OPLS^58–60^ and CHARMM36^61^ force fields. Explicit water was simulated using the TIP3P model.^62,63^ Simulations were conducted following standard protocols^64^ in the NPT ensemble. Temperature and density were controlled over all stages to ensure stability of the simulations. Apo protein simulations were run for 500 ns, whereas kinase and ligand systems were simulated for 100 ns. Preparation of the protein for EDDA calculations was done using Chimera.^65^

## Supporting information

MS identification report

## Figures

Figures were prepared in ChimeraX (ver. 1.7),^66,67^ IBS 2.0^68^ and GIMP (ver. 2.10.36).

## Supporting Information

MS/MS spectra of DYRK1B (ZIP)

DYRK1B MS/MS data, crystallographic data, CETSA assay, quantum chemical calculations, additional interaction maps, additional molecular modelling data (PDF)

## PDB ID Codes

Crystal structures of DYRK1B and DYRK1A in complex with AZ191 described in this manuscript are available in PDB (Protein Data Bank; https://www.rcsb.org/) with ID’s 8C2Z and 8C3G, respectively.

## Author contributions

Conceptualization: PG, KP, FM, AC

Methodology: PG, KP, FM, MJR, PS, AC

Investigation: PG, KP, FM, MJR, PS, AC

Visualization: PG, KP, FM, MJR, PS

Funding acquisition: KP, AC

Project administration: AC

Supervision: AC

Writing—original draft: PG, KP, FM, AC

Writing—review and editing: PG, KP, FM, GD, AC

## Acknowledgements

This work was supported by a grant from National Science Centre (UMO-2019/34/E/NZ1/00467 and 2022/06/X/NZ1/01418) to AC and KP, respectively, and by NAWA Polish Returns 2018 (PPN/PPO/2018/1/00046/U/00001) to AC. We extend our gratitude to Grzegorz Popowicz (Institute of Structural Biology, Helmholtz Zentrum Munchen, Neuherberg, Germany) for his invaluable support, guidance, and scholarly discussions throughout the project. We acknowledge the MCB Structural Biology Core Facility (supported by the TEAM TECH CORE FACILITY/2017-4/6 grant from the Foundation for Polish Science) for valuable support. X-ray data were collected at the HZB BESSY II 14.1 and P11 at DESY. We thank HZB for the allocation of neutron/synchrotron radiation beamtime. We acknowledge DESY (Hamburg, Germany), a member of the Helmholtz Association HGF, for the provision of experimental facilities. Parts of this research were carried out at PETRA III and we would like to thank Johanna Hakanpää for assistance in using beamline P11. Beamtime was allocated for proposal I-20220342 EC.

