## Supplementary figures and tables for "Structural perspective on the design of selective DYRK1B inhibitors"

### Supplemental Material

### Section 1: Experimental data

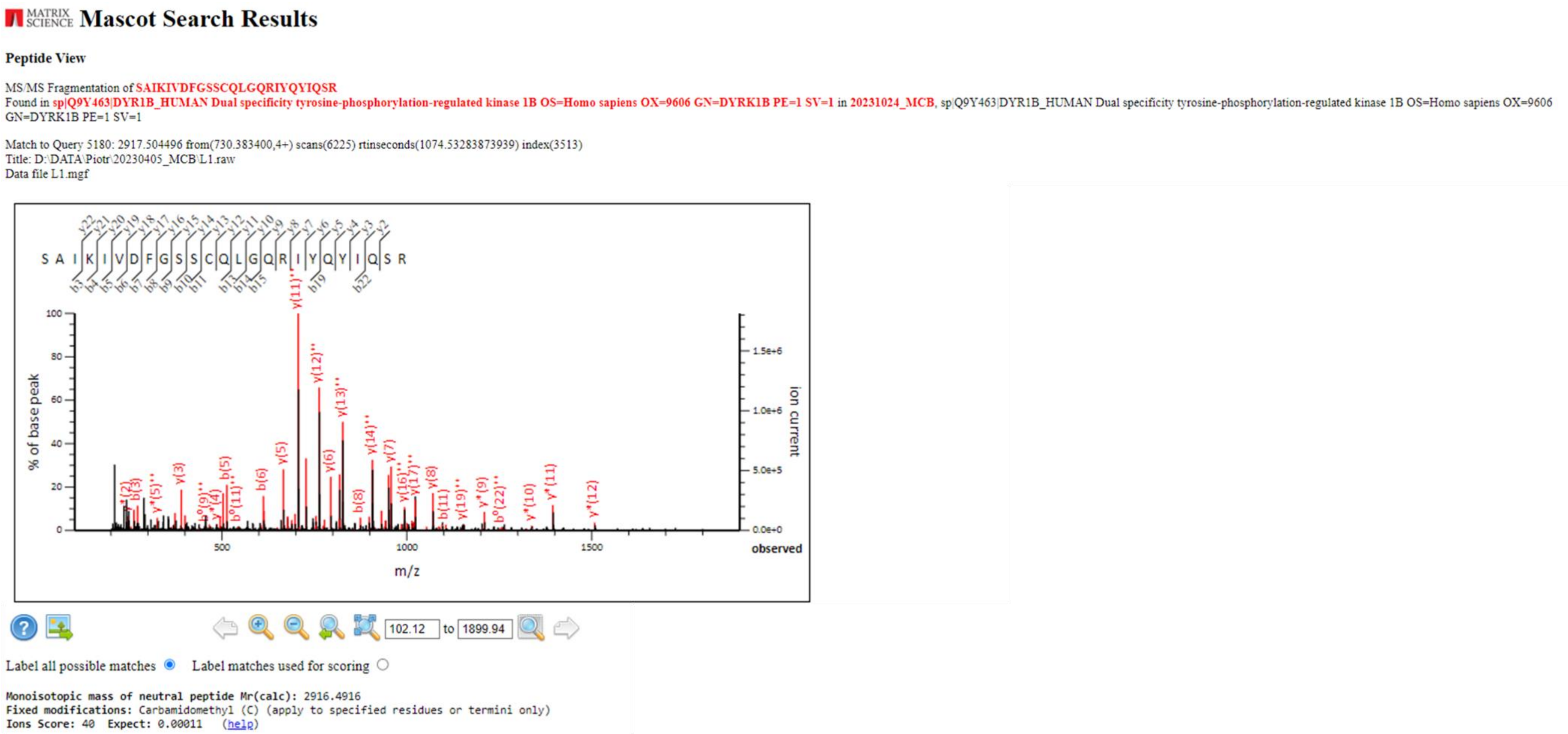


**Figure S1.** Screenshot from the Mascot-based interpretation of the MS/MS spectrum of the ion 730.384+. Almost complete y-ion series and about half of the b-ions series can be observed, according to the Roepstorff's and Fohlmann's nomenclature.

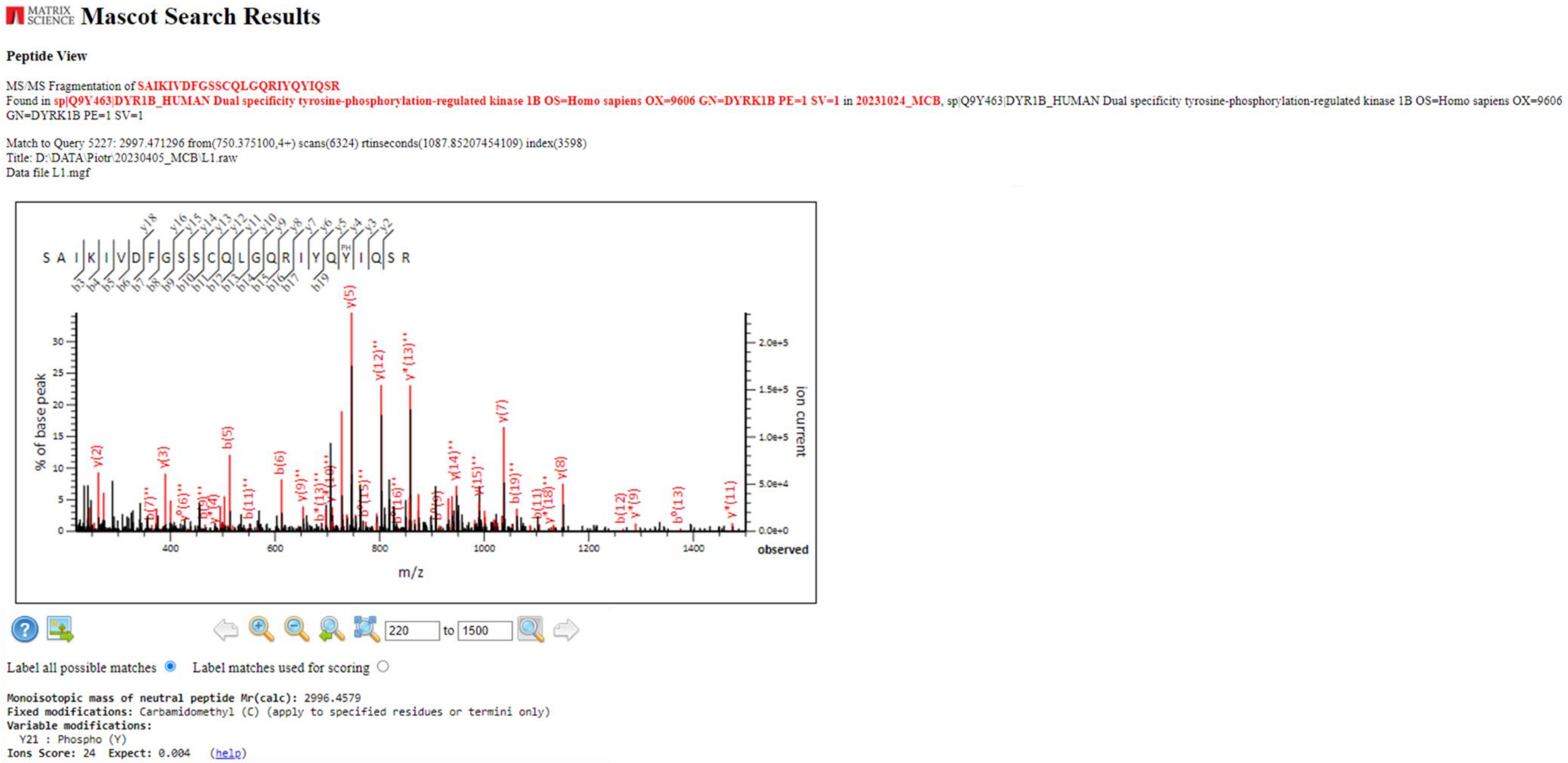


**Figure S2.** Screenshot from the Mascot-based interpretation of the MS/MS spectrum of the ion 750.384+. Almost complete y-ion series and about half of the b-ions series can be observed, according to the Roepstorff's and Fohlmann's nomenclature. There is also visible phosphoserine residue: y18-y5 ions confirm phospho- group presence in the observed peptide.

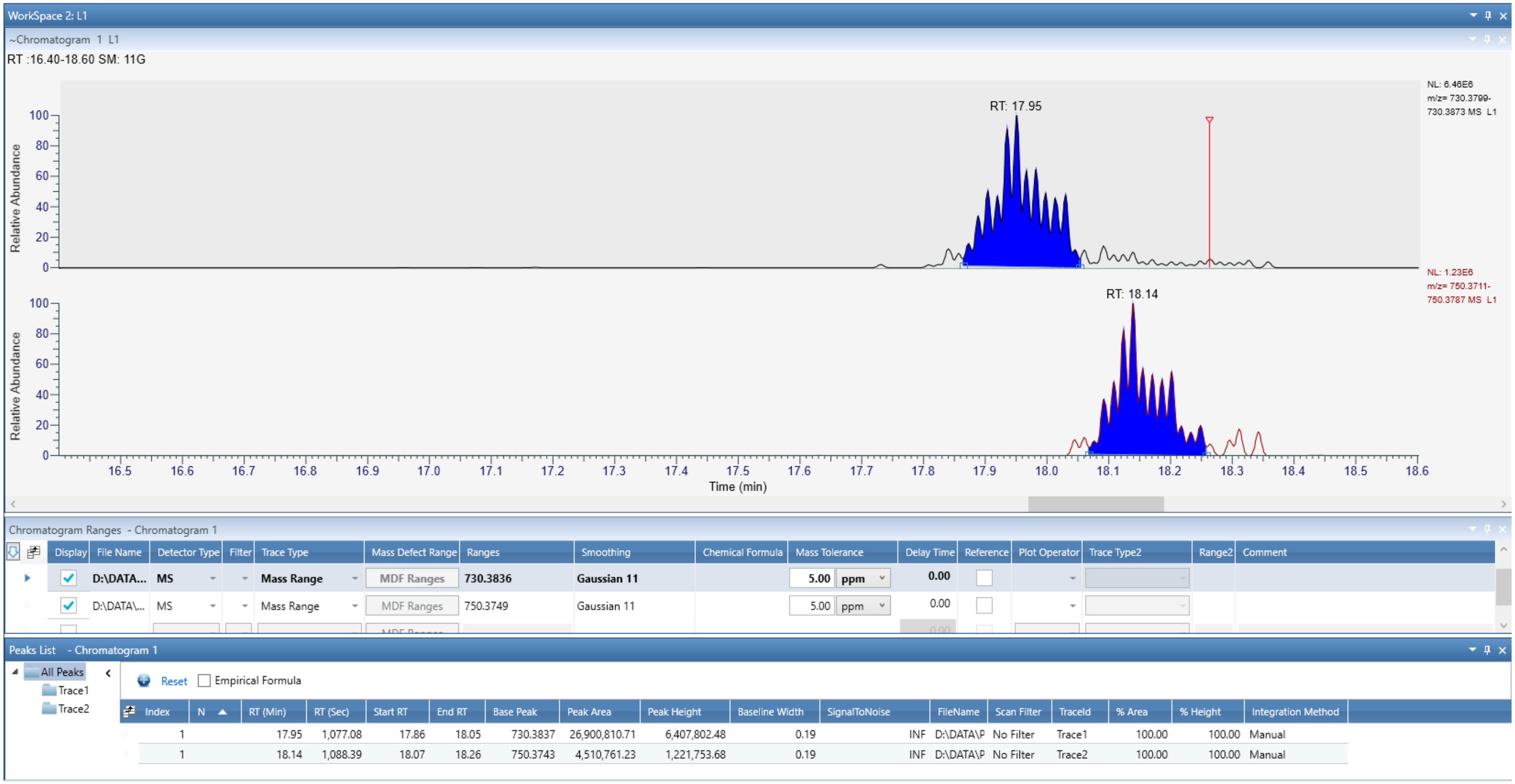


**Figure S3.** Screenshot from the Freestyle software (Thermo, Bremen, Germany) allowing for a simple quantitative comparison of the area under extracted ion chromatograms for both ions. Upper chromatogram: non-phosphorylated peptide, lower chromatogram: phosphorylated peptide, middle panel: additional information about both chromatograms (detailed m/z values, gaussian smoothing cycles and mass tolerance is presented), lower panel: quantitative data („peak area" for both chromatograms) are given.

Please note that the quantitation can be treated estimatively only as the analysis combined MS and MS/MS spectra. The gaussian smoothing cannot exclude gaps between MS spectra devoted for fragmentation scans, however both chromatograms were prepared according to the same procedure, giving reasonable results for quantitation.

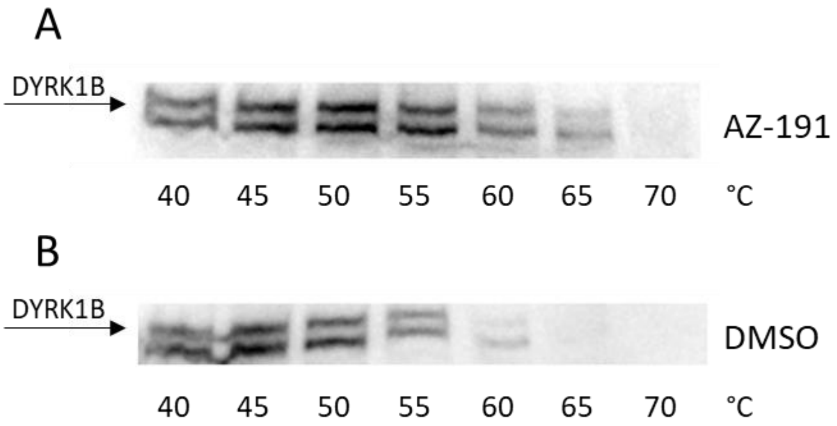


**Figure S4.** CETSA analysis of cell lysates of HEK293T overexpressing FLAG-DYRK1B treated with (A) AZ191 or (B) DMSO. The representative images are shown.

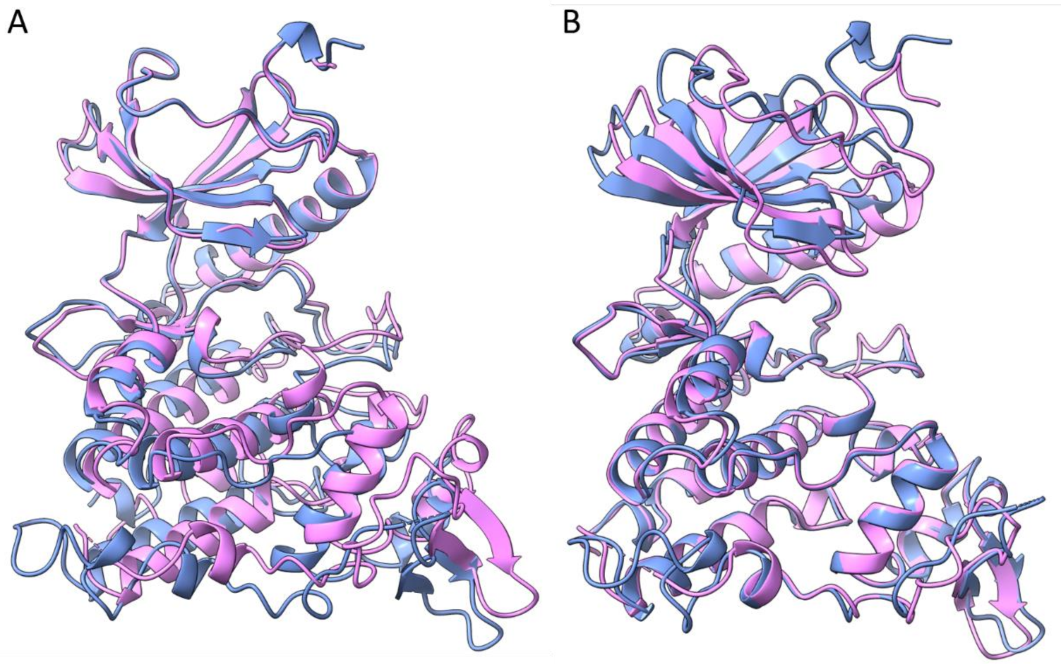


**Figure S5.** Superimposition of DYRK1B (pink) (PDB ID 8C2Z) and DYRK1A (blue) (PDB ID 8C3G). (A) Superimposition of N-lobes, (B) Superimposition of C-lobes.

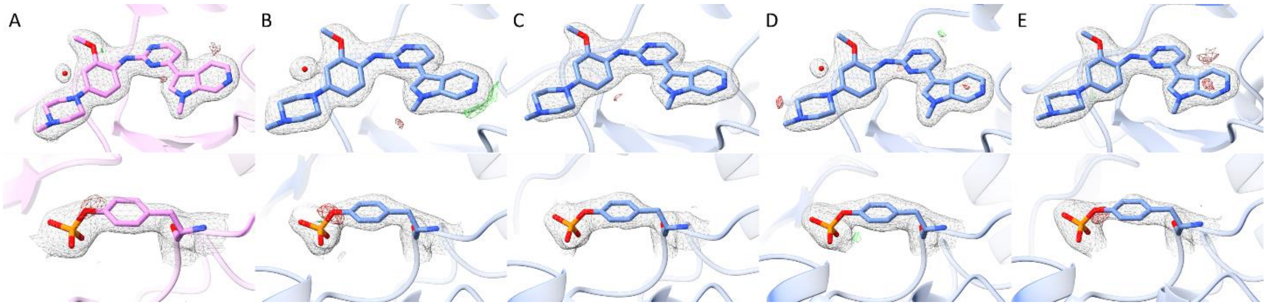


**Figure S6.** Electron density map for AZ191 and phosphorylated tyrosine within YxY motif for DYRK1B and DYRK1A. (A) AZ191 and Tyr273 in DYRK1B/AZ191 crystal structure (PDB ID 8C2Z), (B – E) AZ191 and Tyr321 in DYRK1A/AZ191 crystal structure (PDB ID 8C3G) for models A-D found in the asymmetric unit. Electron density shown as mesh, 2Fo-fc: +1.0σ (light grey); Fo-Fc omit-map: +3.0σ (green); Fo-Fc omit-map: -3.0σ (red).

**Table S1.** Data collection and refinement statistics.

| | DYRK1B AZ191 | DYRK1A AZ191 |
|---|---|---|
| **PDB ID** | 8C2Z | 8C3G |
| **Wavelength** | 1.033 | 0.9184 |
| **Resolution range** | 41.93 - 1.91 (1.95 - 1.91) | 42.77 - 2.08 (2.12 - 2.08) |
| **Space group** | C 2 2 21 | P 21 21 21 |
| **Unit cell** | 115.896 121.509 54.021 (Å) 90 90 90 (°) | 87.064 87.445 228.888 (Å) 90 90 90 (°) |
| **Total reflections** | 242154 (16608) | 536933 (27325) |
| **Unique reflections** | 29064 (1945) | 104770 (5138) |
| **Multiplicity** | 8.3 (8.5) | 5.1 (5.3) |
| **Completeness (%)** | 97.1 (97.8) | 99.3 (99.8) |
| **Mean I/sigma(I)** | 13.8 (1.3) | 9.0 (1.3) |
| **Wilson B-factor** | 38.16 | 35.43 |
| **R-merge** | 0.074 (1.906) | 0.102 (1.251) |
| **R-meas** | 0.084 (2.164) | 0.125 (1.523) |
| **CC1/2** | 0.999 (0.453) | 0.998 (0.464) |
| **Reflections used in refinement** | 29054 (1941) | 104646 (5125) |
| **Reflections used for R-free** | 1488 (98) | 5255 (266) |
| **R-work** | 0.1770 (0.3423) | 0.1941 (0.3031) |
| **R-free** | 0.2202 (0.3701) | 0.2373 (0.3367) |
| **Overall number of atoms** | 2884 | 11918 |
| **In macromolecules** | 2664 | 10848 |
| **In ligands** | 35 | 266 |
| **In waters** | 185 | 804 |
| **Protein residues** | 334 | 1355 |
| **RMSD (bonds)** | 0.013 | 0.008 |
| **RMSD (angles)** | 1.31 | 0.95 |
| **Ramachandran favored (%)** | 95.72 | 96.67 |
| **Ramachandran allowed (%)** | 4.28 | 3.33 |
| **Ramachandran outliers (%)** | 0.00 | 0.00 |
| **Rotamer outliers (%)** | 0.00 | 1.07 |
| **Clashscore** | 3.57 | 4.94 |
| **Average B-factor** | 49.03 | 49.32 |
| **For macromolecules** | 48.92 | 49.32 |
| **For ligands** | 38.79 | 46.36 |
| **For solvent** | 52.54 | 50.34 |

*Data for the highest resolution shell are shown in parentheses.

**Section 2: Computational data**

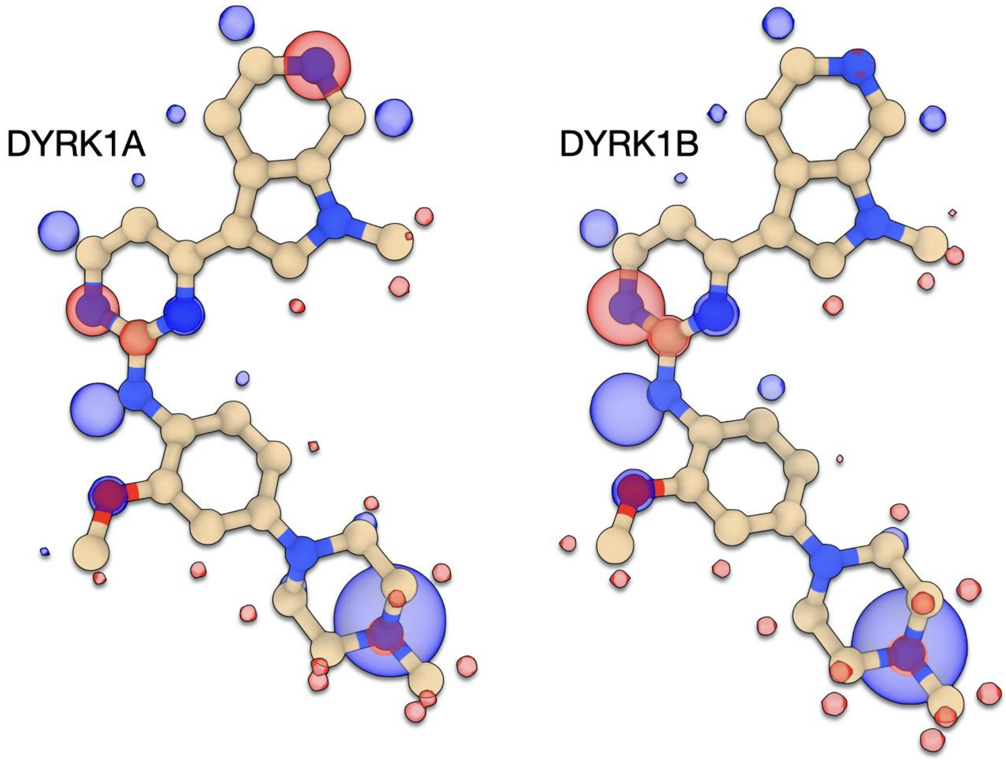


**Figure S7.** Charge Transfer imaps for AZ191 in DYRK1A and DYRK1B.

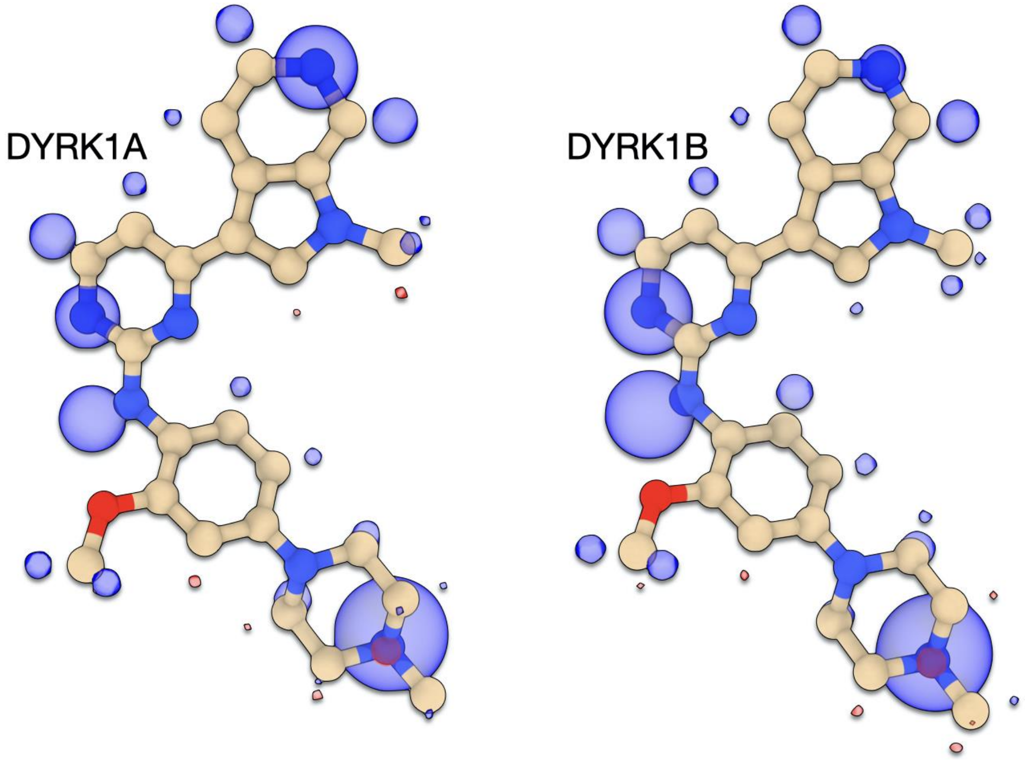


**Figure S8.** Polarization imaps for AZ191 in DYRK1A and DYRK1B.

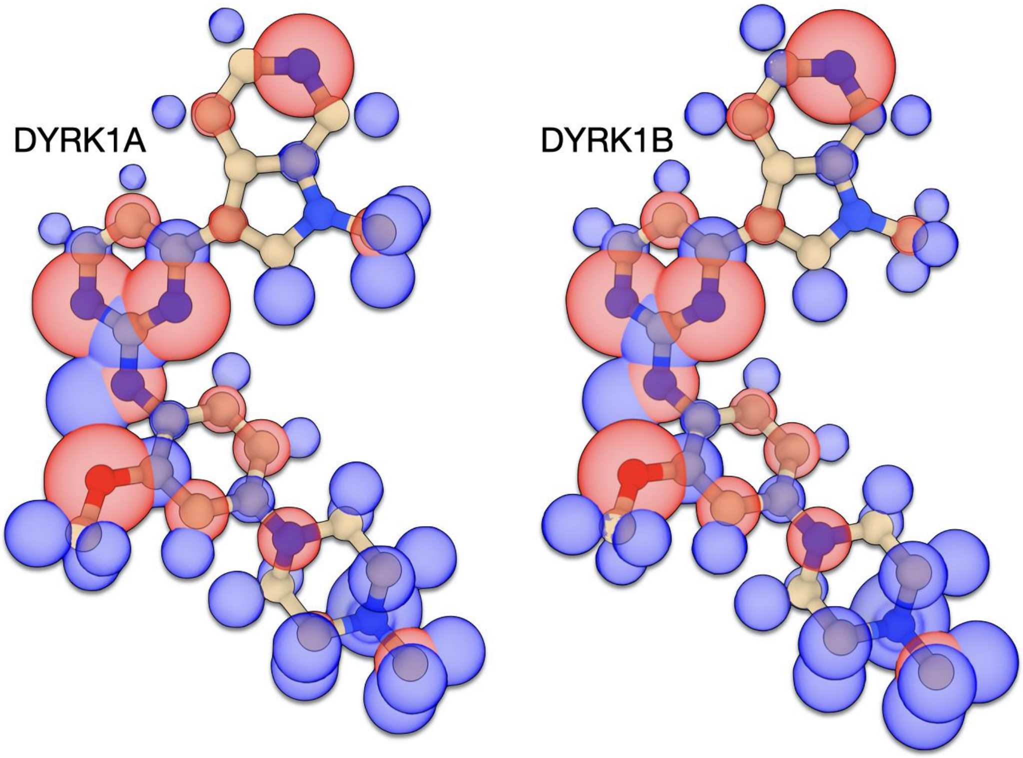


**Figure S9.** Solvation imaps for AZ191 in DYRK1A and DYRK1B.

**Table S2.** Decomposed energies for AZ191 in DYRK1B and DYRK1A. The kinase domain used for the calculations corresponds to subregion B (blue) depicted in Figure S11.

| | DYRK1A | DYRK1B |
|---|---|---|
| ES | 44.25 | 14.06 |
| POL | -81.14 | -97.70 |
| CT | -0.42 | -0.42 |
| REP | 87.69 | 103.24 |
| DISP | -43.35 | -48.09 |
| SOLV | -52.92 | -22.38 |
| INT | -45.90 | -51.29 |

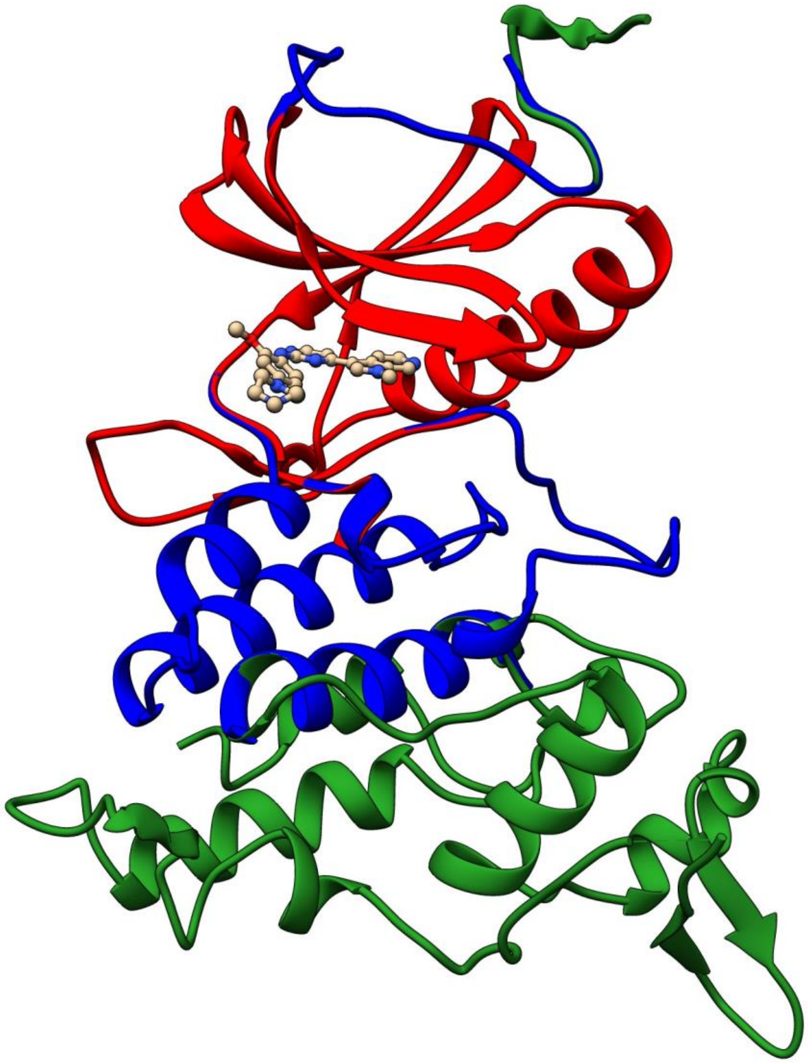

**Figure S10.** The subdomains used for EDDA calculations. The red region corresponds to the smaller subsystem A. The union of red and blue regions corresponds to the subsystem B, which depicts the system used in the calculations. Including the region in green builds the whole kinase as obtained by crystallization.

**Table S3.** Decomposed energies for AZ191 in DYRK1A as a function of time. Calculations using the smaller kinase subsystem A.

| | t (ns) | | | | | |
|---|---|---|---|---|---|---|
| | 0 | 10 | 20 | 30 | 40 | 50 |
| ES | 80.35 | 13.92 | 12.09 | 16.27 | 18.06 | 23.83 |
| POL | -25.86 | -76.87 | -79.93 | -79.74 | -72.58 | -90.62 |
| CT | -0.19 | -1.51 | -0.41 | -0.62 | 0.08 | 0.00 |
| REP | 27.82 | 72.18 | 75.29 | 79.80 | 70.90 | 88.62 |
| DISP | -35.93 | -34.39 | -39.46 | -38.17 | -38.38 | -39.66 |
| SOLV | -72.07 | -4.54 | -2.14 | -16.85 | -17.29 | -22.22 |
| BND | -25.88 | -31.21 | -34.57 | -39.31 | -39.22 | -40.06 |
| RNLys (A) | 3.295 | 3.002 | 3.234 | 2.953 | 3.096 | 2.925 |

| | t (ns) | | | | |
|---|---|---|---|---|---|
| | 60 | 70 | 80 | 90 | 100 |
| ES | 32.18 | 24.33 | 46.27 | 41.70 | 25.22 |
| POL | -85.67 | -83.90 | -58.02 | -63.35 | -98.38 |
| CT | -2.37 | -0.23 | -3.29 | -0.19 | -2.05 |
| REP | 83.59 | 83.24 | 59.74 | 64.68 | 94.55 |
| DISP | -38.09 | -39.43 | -35.26 | -38.06 | -34.23 |
| SOLV | -26.23 | -23.41 | -43.97 | -36.85 | -21.52 |
| BND | -36.59 | -39.40 | -34.54 | -32.06 | -36.41 |
| RNLys (A) | 3.234 | 4.066 | 3.012 | 3.026 | 3.044 |

**Table S4.** Decomposed energies for AZ191 in DYRK1A as a function of time. Calculations using the larger kinase subsystem B.

| | t (ns) | | | | | |
|---|---|---|---|---|---|---|
| | 0 | 20 | 40 | 60 | 80 | 100 |
| ES | 24.59 | 2.47 | 26.73 | 38.88 | 33.63 | 5.58 |
| POL | -47.83 | -78.25 | -65.06 | -78.70 | -55.56 | -96.90 |
| CT | -1.23 | -0.24 | 0.07 | -2.26 | -3.60 | -1.31 |
| REP | 52.48 | 76.42 | 73.17 | 85.28 | 59.79 | 94.91 |
| DISP | -39.84 | -41.54 | -39.72 | -39.50 | -35.39 | -35.90 |
| SOLV | -22.20 | 5.15 | -31.92 | -39.38 | -32.45 | -3.98 |
| INT | -34.02 | -35.99 | -36.74 | -35.68 | -33.59 | -37.60 |

**Table S5.** Decomposed energies for AZ191 in DYRK1B as a function of time. Calculations using the smaller kinase subsystem A.

| | t (ns) | | | | | |
|---|---|---|---|---|---|---|
| | 0 | 10 | 20 | 30 | 40 | 50 |
| ES | 64.65 | 48.50 | 40.40 | 18.03 | 43.45 | 17.95 |
| POL | -50.48 | -63.23 | -45.26 | -45.90 | -57.83 | -59.92 |
| CT | -0.33 | 0.04 | -1.43 | -0.17 | -0.15 | -0.86 |
| REP | 59.37 | 63.56 | 43.35 | 41.66 | 59.86 | 57.70 |
| DISP | -44.81 | -37.64 | -33.36 | -33.70 | -41.96 | -42.56 |
| SOLV | -63.03 | -41.94 | -30.17 | -9.20 | -35.44 | -11.23 |
| INT | -34.63 | -30.70 | -26.46 | -29.27 | -32.06 | -38.92 |
| RNLys (A) | 3.516 | 3.378 | 3.89 | 3.321 | 3.192 | 3.003 |

| | t (ns) | | | | |
|---|---|---|---|---|---|
| | 60 | 70 | 80 | 90 | 100 |
| ES | 21.29 | 32.38 | 50.13 | 22.05 | 43.37 |
| POL | -74.05 | -66.87 | -58.08 | -59.42 | -65.28 |
| CT | -0.06 | -0.22 | -1.15 | -0.02 | -0.33 |
| REP | 75.09 | 68.13 | 59.08 | 55.90 | 66.20 |
| DISP | -44.25 | -38.31 | -39.71 | -38.38 | -41.93 |
| SOLV | -14.25 | -24.14 | -41.97 | -13.04 | -37.53 |
| INT | -36.23 | -29.02 | -31.70 | -32.90 | -35.50 |
| RNLys (A) | 3.61 | 4.067 | 3.692 | 3.991 | 2.942 |

**Table S6.** Decomposed energies for AZ191 in DYRK1B as a function of time. Calculations using the larger kinase subsystem B.

| | t (ns) | | | | | |
|---|---|---|---|---|---|---|
| | 0 | 20 | 40 | 60 | 80 | 100 |
| ES | 2.01 | -28.05 | 2.47 | -37.06 | -2.99 | -13.95 |
| POL | -70.29 | -86.46 | -58.61 | -77.09 | -63.28 | -67.60 |
| CT | 0.00 | -0.75 | 0.03 | -0.05 | -0.88 | -0.27 |
| REP | 81.64 | 78.76 | 63.20 | 76.32 | 63.18 | 66.13 |
| DISP | -48.73 | -36.55 | -44.55 | -45.68 | -41.97 | -42.78 |
| SOLV | -6.56 | 36.46 | 3.89 | 45.71 | 16.73 | 26.29 |
| INT | -41.93 | -36.58 | -33.57 | -37.86 | -29.22 | -32.18 |

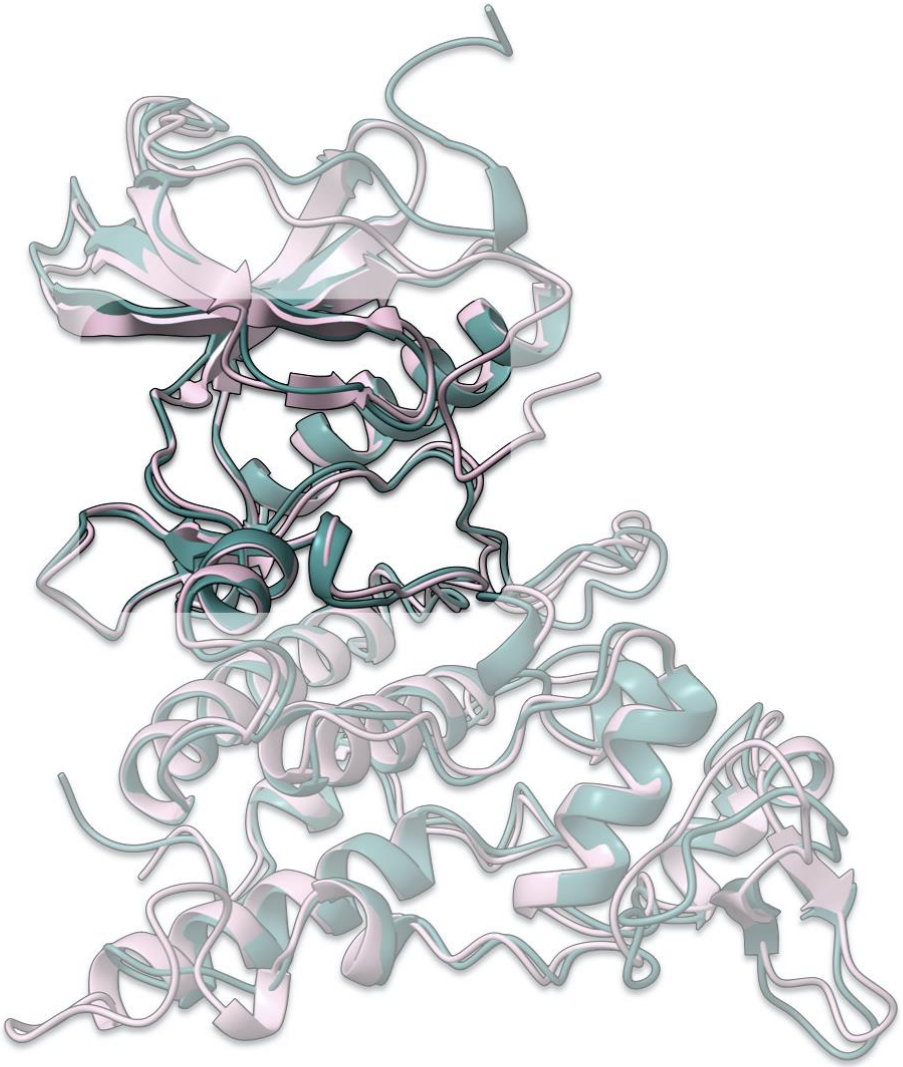

**Figure S11.** Overlay of the structures of DYRK1A (pink) and DYRK1B (green) at 380 ns, where the pocket-narrowness is equivalent.

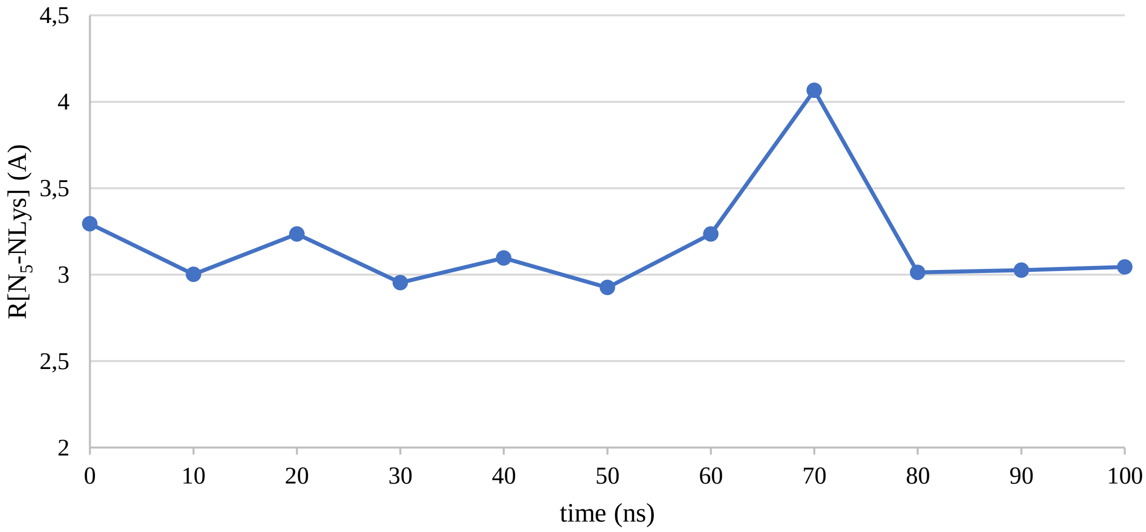

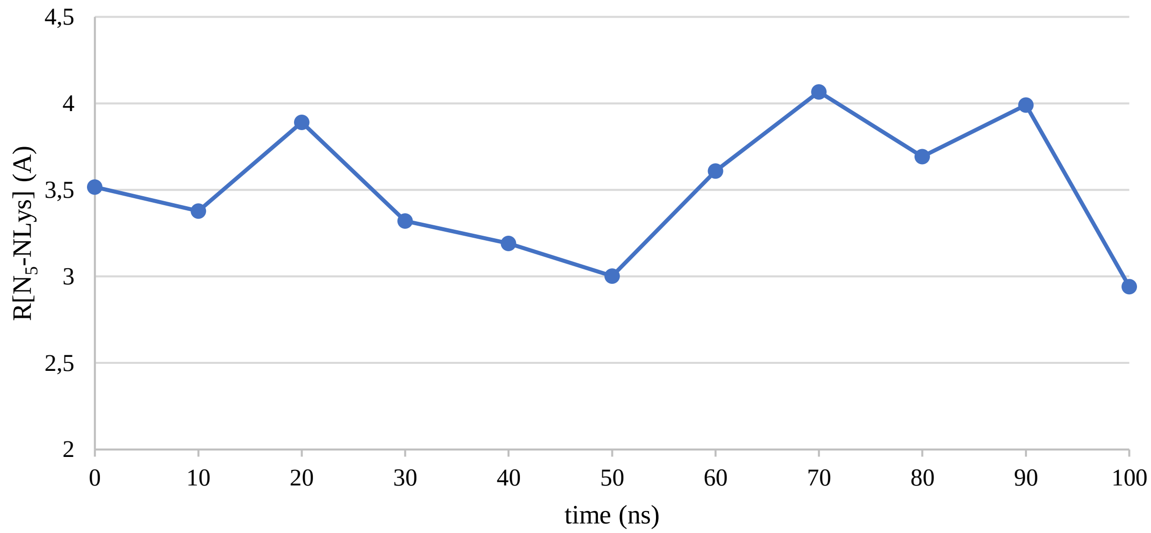


**Figure S12.** $N_5$-$N_{LYS}$ distance between the methylpyrrolepyridine group of the ligand and the catalytic lysin of DYRK1A (up) and DYRK1B (down) as a function of time.

**Table S7.** Decomposed energies on the smaller kinase subdomain A.

| | DYRK1A | DYRK1B |
|---|---|---|
| ES | 65.60 | 58.83 |
| POL | -56.55 | -58.82 |
| CT | -1.30 | -2.71 |
| REP | 56.60 | 62.75 |
| DISP | -37.40 | -40.98 |
| SOLV | -64.61 | -61.53 |
| INT | -37.67 | -42.45 |
