## Supplementary material for "Structural perspective on the design of selective DYRK1B inhibitors": MS identification report: Mascot Search Results_ sp_Q9Y463_DYR1B_HUMAN Dual specificity tyrosine-phosphorylation-regulated kinase 1B OS=Homo sapiens OX=9606 GN=DYRK1B PE=1 SV=1.html

### MASCOT Search Results

#### Protein View: sp|Q9Y463|DYR1B\_HUMAN Dual specificity tyrosine-phosphorylation-regulated kinase 1B OS=Homo sapiens OX=9606 GN=DYRK1B PE=1 SV=1

##### sp|Q9Y463|DYR1B\_HUMAN Dual specificity tyrosine-phosphorylation-regulated kinase 1B OS=Homo sapiens OX=9606 GN=DYRK1B PE=1 SV=1

|  |  |
| --- | --- |
| Database: | 20231024\_MCB |
| Score: | 13672 |
| Monoisotopic mass (M_r_): | 69725 |
| Calculated pI: | 9.25 |

Sequence similarity is available as an NCBI BLAST search of sp|Q9Y463|DYR1B\_HUMAN Dual specificity tyrosine-phosphorylation-regulated kinase 1B OS=Homo sapiens OX=9606 GN=DYRK1B PE=1 SV=1 against nr.

##### Search parameters

|  |  |
| --- | --- |
| MS data file: | `L1.mgf` |
| Enzyme: | Trypsin: cuts C-term side of KR unless next residue is P. |
| Fixed modifications: | Carbamidomethyl (C) |
| Variable modifications: | Phospho (ST), Oxidation (M), Phospho (Y) |

##### Protein sequence coverage: 38%

Matched peptides shown in ***bold red***.

|  |  |  |  |  |  |
| --- | --- | --- | --- | --- | --- |
| `1` | `MAVPPGHGPF` | `SGFPGPQEHT` | `QVLPDVRLLP` | `RRLPLAFRDA` | `TSAPLRKLSV` |
| `51` | `DLIKTYKHIN` | `EVYYAKKKRR` | `AQQAPPQDSS` | `NKKEKKVLNH` | `GYDDDNHDYI` |
| `101` | `VRSGERWLER` | `YEIDSLIGKG` | `SFGQVVKAYD` | `HQTQELVAIK` | `IIKNKKAFLN` |
| `151` | `QAQIELRLLE` | `LMNQHDTEMK` | `YYIVHLKRHF` | `MFRNHLCLVF` | `ELLSYNLYDL` |
| `201` | `LRNTHFRGVS` | `LNLTRKLAQQ` | `LCTALLFLAT` | `PELSIIHCDL` | `KPENILLCNP` |
| `251` | `KRSAIKIVDF` | `GSSCQLGQRI` | `YQYIQSRFYR` | `SPEVLLGTPY` | `DLAIDMWSLG` |
| `301` | `CILVEMHTGE` | `PLFSGSNEVD` | `QMNRIVEVLG` | `IPPAAMLDQA` | `PKARKYFERL` |
| `351` | `PGGGWTLRRT` | `KELRKDYQGP` | `GTRRLQEVLG` | `VQTGGPGGRR` | `AGEPGHSPAD` |
| `401` | `YLRFQDLVLR` | `MLEYEPAARI` | `SPLGALQHGF` | `FRRTADEATN` | `TGPAGSSAST` |
| `451` | `SPAPLDTCPS` | `SSTASSISSS` | `GGSSGSSSDN` | `RTYRYSNRYC` | `GGPGPPITDC` |
| `501` | `EMNSPQVPPS` | `QPLRPWAGGD` | `VPHKTHQAPA` | `SASSLPGTGA` | `QLPPQPRYLG` |
| `551` | `RPPSPTSPPP` | `PELMDVSLVG` | `GPADCSPPHP` | `APAPQHPAAS` | `ALRTRMTGGR` |
| `601` | `PPLPPPDDPA` | `TLGPHLGLRG` | `VPQSTAASS` |  |  |

Unformatted sequence string: 629 residues (for pasting into other applications).

|  |  |  |  |
| --- | --- | --- | --- |
| Sort by | residue number | increasing mass | decreasing mass |
| Show | matched peptides only | predicted peptides also |  |
| Show | uncorrected delta | delta corrected for 13C |  |

| Query | Start | – | End | Observed | Mr(expt) | Mr(calc) | ppm | M | Score | Expect | Rank | U | Peptide |
| --- | --- | --- | --- | --- | --- | --- | --- | --- | --- | --- | --- | --- | --- |
| 4637 | 84 | – | 102 | 466.8299 | 2329.1131 | 2329.1087 | 1.90 | 2 | 49 | 1.3e-05 | 1Score **> 13** indicates **identity** | U | K.EKKVLNHGYDDDNHDYIVR.S |
| 4638 | 84 | – | 102 | 777.7117 | 2330.1133 | 2329.1087 | 431 | 2 | 75 | 3.1e-08 | 1Score **> 13** indicates **identity** | U | K.EKKVLNHGYDDDNHDYIVR.S |
| 4639 | 84 | – | 102 | 583.5361 | 2330.1153 | 2329.1087 | 432 | 2 | 82 | 6.6e-09 | 1Score **> 13** indicates **identity** | U | K.EKKVLNHGYDDDNHDYIVR.S |
| 4334 | 86 | – | 102 | 1036.9941 | 2071.9736 | 2071.9712 | 1.20 | 1 | 93 | 5.4e-10 | 1Score **> 13** indicates **identity** | U | K.KVLNHGYDDDNHDYIVR.S |
| 4335 | 86 | – | 102 | 1036.9968 | 2071.9790 | 2071.9712 | 3.81 | 1 | 106 | 2.4e-11 | 1Score **> 13** indicates **identity** | U | K.KVLNHGYDDDNHDYIVR.S |
| 4336 | 86 | – | 102 | 691.9940 | 2072.9602 | 2071.9712 | 477 | 1 | 69 | 1.2e-07 | 1Score **> 13** indicates **identity** | U | K.KVLNHGYDDDNHDYIVR.S |
| 4337 | 86 | – | 102 | 519.2491 | 2072.9673 | 2071.9712 | 481 | 1 | 52 | 7e-06 | 1Score **> 13** indicates **identity** | U | K.KVLNHGYDDDNHDYIVR.S |
| 4338 | 86 | – | 102 | 1037.4922 | 2072.9698 | 2071.9712 | 482 | 1 | 67 | 1.8e-07 | 1Score **> 13** indicates **identity** | U | K.KVLNHGYDDDNHDYIVR.S |
| 4339 | 86 | – | 102 | 415.6014 | 2072.9706 | 2071.9712 | 482 | 1 | 15 | 0.029 | 1Score **> 13** indicates **identity** | U | K.KVLNHGYDDDNHDYIVR.S |
| 4340 | 86 | – | 102 | 691.9976 | 2072.9710 | 2071.9712 | 483 | 1 | 46 | 2.7e-05 | 1Score **> 13** indicates **identity** | U | K.KVLNHGYDDDNHDYIVR.S |
| 4343 | 86 | – | 102 | 519.2517 | 2072.9777 | 2071.9712 | 486 | 1 | 23 | 0.0045 | 1Score **> 13** indicates **identity** | U | K.KVLNHGYDDDNHDYIVR.S |
| 4344 | 86 | – | 102 | 692.0001 | 2072.9785 | 2071.9712 | 486 | 1 | 68 | 1.5e-07 | 1Score **> 13** indicates **identity** | U | K.KVLNHGYDDDNHDYIVR.S |
| 4345 | 86 | – | 102 | 519.2519 | 2072.9785 | 2071.9712 | 486 | 1 | 77 | 2.1e-08 | 1Score **> 13** indicates **identity** | U | K.KVLNHGYDDDNHDYIVR.S |
| 4346 | 86 | – | 102 | 519.2521 | 2072.9793 | 2071.9712 | 487 | 1 | 56 | 2.6e-06 | 1Score **> 13** indicates **identity** | U | K.KVLNHGYDDDNHDYIVR.S |
| 4347 | 86 | – | 102 | 415.6032 | 2072.9796 | 2071.9712 | 487 | 1 | 13 | 0.049 | 1Score **> 13** indicates **identity** | U | K.KVLNHGYDDDNHDYIVR.S |
| 4847 | 86 | – | 106 | 626.5499 | 2502.1705 | 2501.1684 | 401 | 2 | 39 | 0.00011 | 1Score **> 13** indicates **identity** | U | K.KVLNHGYDDDNHDYIVRSGER.W |
| 4848 | 86 | – | 106 | 835.0656 | 2502.1750 | 2501.1684 | 402 | 2 | 26 | 0.0026 | 1Score **> 13** indicates **identity** | U | K.KVLNHGYDDDNHDYIVRSGER.W |
| 4849 | 86 | – | 106 | 501.4423 | 2502.1751 | 2501.1684 | 403 | 2 | 42 | 6.6e-05 | 1Score **> 13** indicates **identity** | U | K.KVLNHGYDDDNHDYIVRSGER.W |
| 4850 | 86 | – | 106 | 626.5511 | 2502.1753 | 2501.1684 | 403 | 2 | 29 | 0.0012 | 1Score **> 13** indicates **identity** | U | K.KVLNHGYDDDNHDYIVRSGER.W |
| 4112 | 87 | – | 102 | 972.9460 | 1943.8774 | 1943.8762 | 0.64 | 0 | 79 | 1.3e-08 | 1Score **> 13** indicates **identity** | U | K.VLNHGYDDDNHDYIVR.S |
| 4113 | 87 | – | 102 | 972.9479 | 1943.8812 | 1943.8762 | 2.59 | 0 | 86 | 2.3e-09 | 1Score **> 13** indicates **identity** | U | K.VLNHGYDDDNHDYIVR.S |
| 4114 | 87 | – | 102 | 972.9481 | 1943.8816 | 1943.8762 | 2.80 | 0 | 89 | 1.2e-09 | 1Score **> 13** indicates **identity** | U | K.VLNHGYDDDNHDYIVR.S |
| 4115 | 87 | – | 102 | 486.9777 | 1943.8817 | 1943.8762 | 2.82 | 0 | 57 | 1.9e-06 | 1Score **> 13** indicates **identity** | U | K.VLNHGYDDDNHDYIVR.S |
| 4116 | 87 | – | 102 | 648.9690 | 1943.8852 | 1943.8762 | 4.61 | 0 | 18 | 0.018 | 1Score **> 13** indicates **identity** | U | K.VLNHGYDDDNHDYIVR.S |
| 4117 | 87 | – | 102 | 648.9752 | 1943.9038 | 1943.8762 | 14.2 | 0 | 37 | 0.00021 | 1Score **> 13** indicates **identity** | U | K.VLNHGYDDDNHDYIVR.S |
| 4118 | 87 | – | 102 | 649.2960 | 1944.8662 | 1943.8762 | 509 | 0 | 58 | 1.4e-06 | 1Score **> 13** indicates **identity** | U | K.VLNHGYDDDNHDYIVR.S |
| 4119 | 87 | – | 102 | 487.2242 | 1944.8677 | 1943.8762 | 510 | 0 | 20 | 0.01 | 1Score **> 13** indicates **identity** | U | K.VLNHGYDDDNHDYIVR.S |
| 4120 | 87 | – | 102 | 649.2971 | 1944.8695 | 1943.8762 | 511 | 0 | 45 | 3.3e-05 | 1Score **> 13** indicates **identity** | U | K.VLNHGYDDDNHDYIVR.S |
| 4121 | 87 | – | 102 | 649.2977 | 1944.8713 | 1943.8762 | 512 | 0 | 50 | 8.9e-06 | 1Score **> 13** indicates **identity** | U | K.VLNHGYDDDNHDYIVR.S |
| 4122 | 87 | – | 102 | 487.2273 | 1944.8801 | 1943.8762 | 516 | 0 | 46 | 2.7e-05 | 1Score **> 13** indicates **identity** | U | K.VLNHGYDDDNHDYIVR.S |
| 4123 | 87 | – | 102 | 487.2273 | 1944.8801 | 1943.8762 | 516 | 0 | 42 | 5.9e-05 | 1Score **> 13** indicates **identity** | U | K.VLNHGYDDDNHDYIVR.S |
| 4124 | 87 | – | 102 | 649.3010 | 1944.8812 | 1943.8762 | 517 | 0 | 41 | 7.3e-05 | 1Score **> 13** indicates **identity** | U | K.VLNHGYDDDNHDYIVR.S |
| 4125 | 87 | – | 102 | 649.3011 | 1944.8815 | 1943.8762 | 517 | 0 | 58 | 1.7e-06 | 1Score **> 13** indicates **identity** | U | K.VLNHGYDDDNHDYIVR.S |
| 4126 | 87 | – | 102 | 487.2277 | 1944.8817 | 1943.8762 | 517 | 0 | 44 | 4.3e-05 | 1Score **> 13** indicates **identity** | U | K.VLNHGYDDDNHDYIVR.S |
| 4127 | 87 | – | 102 | 487.2279 | 1944.8825 | 1943.8762 | 518 | 0 | 31 | 0.00087 | 1Score **> 13** indicates **identity** | U | K.VLNHGYDDDNHDYIVR.S |
| 4128 | 87 | – | 102 | 649.3015 | 1944.8827 | 1943.8762 | 518 | 0 | 58 | 1.4e-06 | 1Score **> 13** indicates **identity** | U | K.VLNHGYDDDNHDYIVR.S |
| 4129 | 87 | – | 102 | 973.4489 | 1944.8832 | 1943.8762 | 518 | 0 | 63 | 4.5e-07 | 1Score **> 13** indicates **identity** | U | K.VLNHGYDDDNHDYIVR.S |
| 4130 | 87 | – | 102 | 487.2282 | 1944.8837 | 1943.8762 | 518 | 0 | 19 | 0.013 | 1Score **> 13** indicates **identity** | U | K.VLNHGYDDDNHDYIVR.S |
| 4679 | 87 | – | 106 | 792.0312 | 2373.0718 | 2373.0734 | -0.68 | 1 | 30 | 0.001 | 1Score **> 13** indicates **identity** | U | K.VLNHGYDDDNHDYIVRSGER.W |
| 4683 | 87 | – | 106 | 792.3636 | 2374.0690 | 2373.0734 | 420 | 1 | 35 | 0.00034 | 1Score **> 13** indicates **identity** | U | K.VLNHGYDDDNHDYIVRSGER.W |
| 4684 | 87 | – | 106 | 475.8221 | 2374.0741 | 2373.0734 | 422 | 1 | 33 | 0.00046 | 1Score **> 13** indicates **identity** | U | K.VLNHGYDDDNHDYIVRSGER.W |
| 4685 | 87 | – | 106 | 1188.0468 | 2374.0790 | 2373.0734 | 424 | 1 | 39 | 0.00012 | 1Score **> 13** indicates **identity** | U | K.VLNHGYDDDNHDYIVRSGER.W |
| 4686 | 87 | – | 106 | 594.5276 | 2374.0813 | 2373.0734 | 425 | 1 | 36 | 0.00027 | 1Score **> 13** indicates **identity** | U | K.VLNHGYDDDNHDYIVRSGER.W |
| 4687 | 87 | – | 106 | 1188.0498 | 2374.0850 | 2373.0734 | 426 | 1 | 33 | 0.00047 | 1Score **> 13** indicates **identity** | U | K.VLNHGYDDDNHDYIVRSGER.W |
| 4299 | 103 | – | 119 | 513.7711 | 2051.0553 | 2050.0483 | 491 | 2 | 14 | 0.042 | 1Score **> 13** indicates **identity** | U | R.SGERWLERYEIDSLIGK.G |
| 4300 | 103 | – | 119 | 1026.5353 | 2051.0560 | 2050.0483 | 492 | 2 | 23 | 0.0051 | 1Score **> 13** indicates **identity** | U | R.SGERWLERYEIDSLIGK.G |
| 4301 | 103 | – | 119 | 513.7714 | 2051.0565 | 2050.0483 | 492 | 2 | 17 | 0.019 | 1Score **> 13** indicates **identity** | U | R.SGERWLERYEIDSLIGK.G |
| 3452 | 107 | – | 119 | 811.4346 | 1620.8546 | 1620.8511 | 2.18 | 1 | 52 | 7.1e-06 | 1Score **> 13** indicates **identity** | U | R.WLERYEIDSLIGK.G |
| 3454 | 107 | – | 119 | 811.4358 | 1620.8570 | 1620.8511 | 3.66 | 1 | 50 | 1.1e-05 | 1Score **> 13** indicates **identity** | U | R.WLERYEIDSLIGK.G |
| 3456 | 107 | – | 119 | 811.9363 | 1621.8580 | 1620.8511 | 621 | 1 | 47 | 1.9e-05 | 1Score **> 13** indicates **identity** | U | R.WLERYEIDSLIGK.G |
| 3457 | 107 | – | 119 | 811.9364 | 1621.8582 | 1620.8511 | 621 | 1 | 43 | 5.3e-05 | 1Score **> 13** indicates **identity** | U | R.WLERYEIDSLIGK.G |
| 4754 | 107 | – | 127 | 606.8278 | 2423.2821 | 2423.2849 | -1.15 | 2 | 21 | 0.0078 | 1Score **> 13** indicates **identity** | U | R.WLERYEIDSLIGKGSFGQVVK.A |
| 4755 | 107 | – | 127 | 808.7716 | 2423.2930 | 2423.2849 | 3.34 | 2 | 50 | 9.4e-06 | 1Score **> 13** indicates **identity** | U | R.WLERYEIDSLIGKGSFGQVVK.A |
| 4756 | 107 | – | 127 | 1212.6548 | 2423.2950 | 2423.2849 | 4.20 | 2 | 83 | 4.7e-09 | 1Score **> 13** indicates **identity** | U | R.WLERYEIDSLIGKGSFGQVVK.A |
| 4758 | 107 | – | 127 | 607.0812 | 2424.2957 | 2423.2849 | 417 | 2 | 38 | 0.00016 | 1Score **> 13** indicates **identity** | U | R.WLERYEIDSLIGKGSFGQVVK.A |
| 4759 | 107 | – | 127 | 809.1059 | 2424.2959 | 2423.2849 | 417 | 2 | 55 | 3.1e-06 | 1Score **> 13** indicates **identity** | U | R.WLERYEIDSLIGKGSFGQVVK.A |
| 4760 | 107 | – | 127 | 607.0813 | 2424.2961 | 2423.2849 | 417 | 2 | 39 | 0.00013 | 1Score **> 13** indicates **identity** | U | R.WLERYEIDSLIGKGSFGQVVK.A |
| 4761 | 107 | – | 127 | 607.0823 | 2424.3001 | 2423.2849 | 419 | 2 | 33 | 0.00048 | 1Score **> 13** indicates **identity** | U | R.WLERYEIDSLIGKGSFGQVVK.A |
| 4762 | 107 | – | 127 | 1213.1576 | 2424.3006 | 2423.2849 | 419 | 2 | 58 | 1.5e-06 | 1Score **> 13** indicates **identity** | U | R.WLERYEIDSLIGKGSFGQVVK.A |
| 2529 | 111 | – | 119 | 1037.5522 | 1036.5449 | 1036.5441 | 0.84 | 0 | 35 | 0.00032 | 1Score **> 13** indicates **identity** | U | R.YEIDSLIGK.G |
| 2530 | 111 | – | 119 | 1037.5543 | 1036.5470 | 1036.5441 | 2.87 | 0 | 35 | 0.00031 | 1Score **> 13** indicates **identity** | U | R.YEIDSLIGK.G |
| 2531 | 111 | – | 119 | 519.2808 | 1036.5470 | 1036.5441 | 2.89 | 0 | 30 | 0.001 | 1Score **> 13** indicates **identity** | U | R.YEIDSLIGK.G |
| 3903 | 111 | – | 127 | 920.4988 | 1838.9830 | 1838.9778 | 2.85 | 1 | 67 | 1.9e-07 | 1Score **> 13** indicates **identity** | U | R.YEIDSLIGKGSFGQVVK.A |
| 3904 | 111 | – | 127 | 920.4999 | 1838.9852 | 1838.9778 | 4.04 | 1 | 75 | 3.3e-08 | 1Score **> 13** indicates **identity** | U | R.YEIDSLIGKGSFGQVVK.A |
| 3905 | 111 | – | 127 | 614.0032 | 1838.9878 | 1838.9778 | 5.41 | 1 | 47 | 1.9e-05 | 1Score **> 13** indicates **identity** | U | R.YEIDSLIGKGSFGQVVK.A |
| 3907 | 111 | – | 127 | 921.0026 | 1839.9906 | 1838.9778 | 551 | 1 | 67 | 2.2e-07 | 1Score **> 13** indicates **identity** | U | R.YEIDSLIGKGSFGQVVK.A |
| 4624 | 120 | – | 140 | 580.5604 | 2318.2125 | 2317.2066 | 434 | 1 | 21 | 0.0077 | 1Score **> 13** indicates **identity** | U | K.GSFGQVVKAYDHQTQELVAIK.I |
| 4625 | 120 | – | 140 | 773.7467 | 2318.2183 | 2317.2066 | 437 | 1 | 33 | 0.00051 | 1Score **> 13** indicates **identity** | U | K.GSFGQVVKAYDHQTQELVAIK.I |
| 3257 | 128 | – | 140 | 505.9297 | 1514.7673 | 1514.7729 | -3.71 | 0 | 28 | 0.0015 | 1Score **> 13** indicates **identity** | U | K.AYDHQTQELVAIK.I |
| 3258 | 128 | – | 140 | 758.3958 | 1514.7770 | 1514.7729 | 2.75 | 0 | 64 | 4.2e-07 | 1Score **> 13** indicates **identity** | U | K.AYDHQTQELVAIK.I |
| 3259 | 128 | – | 140 | 505.9330 | 1514.7772 | 1514.7729 | 2.83 | 0 | 26 | 0.0028 | 1Score **> 13** indicates **identity** | U | K.AYDHQTQELVAIK.I |
| 3260 | 128 | – | 140 | 758.3959 | 1514.7772 | 1514.7729 | 2.88 | 0 | 55 | 3e-06 | 1Score **> 13** indicates **identity** | U | K.AYDHQTQELVAIK.I |
| 3992 | 128 | – | 143 | 624.3544 | 1870.0414 | 1869.0360 | 538 | 1 | 34 | 0.0004 | 1Score **> 13** indicates **identity** | U | K.AYDHQTQELVAIKIIK.N |
| 3994 | 128 | – | 143 | 936.0300 | 1870.0454 | 1869.0360 | 540 | 1 | 47 | 1.9e-05 | 1Score **> 13** indicates **identity** | U | K.AYDHQTQELVAIKIIK.N |
| 3995 | 128 | – | 143 | 624.3558 | 1870.0456 | 1869.0360 | 540 | 1 | 49 | 1.4e-05 | 1Score **> 13** indicates **identity** | U | K.AYDHQTQELVAIKIIK.N |
| 3996 | 128 | – | 143 | 936.0307 | 1870.0468 | 1869.0360 | 541 | 1 | 52 | 6.2e-06 | 1Score **> 13** indicates **identity** | U | K.AYDHQTQELVAIKIIK.N |
| 4436 | 128 | – | 145 | 705.0679 | 2112.1819 | 2111.1738 | 477 | 2 | 68 | 1.6e-07 | 1Score **> 13** indicates **identity** | U | K.AYDHQTQELVAIKIIKNK.K |
| 3569 | 144 | – | 157 | 418.9940 | 1671.9469 | 1671.9420 | 2.94 | 2 | 30 | 0.00091 | 1Score **> 13** indicates **identity** | U | K.NKKAFLNQAQIELR.L |
| 3570 | 144 | – | 157 | 837.4808 | 1672.9470 | 1671.9420 | 601 | 2 | 63 | 5.6e-07 | 1Score **> 13** indicates **identity** | U | K.NKKAFLNQAQIELR.L |
| 3571 | 144 | – | 157 | 558.6577 | 1672.9513 | 1671.9420 | 604 | 2 | 59 | 1.2e-06 | 1Score **> 13** indicates **identity** | U | K.NKKAFLNQAQIELR.L |
| 3088 | 146 | – | 157 | 715.9097 | 1429.8048 | 1429.8041 | 0.52 | 1 | 88 | 1.6e-09 | 1Score **> 13** indicates **identity** | U | K.KAFLNQAQIELR.L |
| 3089 | 146 | – | 157 | 715.9106 | 1429.8066 | 1429.8041 | 1.78 | 1 | 88 | 1.6e-09 | 1Score **> 13** indicates **identity** | U | K.KAFLNQAQIELR.L |
| 3090 | 146 | – | 157 | 477.6100 | 1429.8082 | 1429.8041 | 2.84 | 1 | 48 | 1.5e-05 | 1Score **> 13** indicates **identity** | U | K.KAFLNQAQIELR.L |
| 3091 | 146 | – | 157 | 715.9125 | 1429.8104 | 1429.8041 | 4.43 | 1 | 68 | 1.5e-07 | 1Score **> 13** indicates **identity** | U | K.KAFLNQAQIELR.L |
| 3094 | 146 | – | 157 | 477.9377 | 1430.7913 | 1429.8041 | 690 | 1 | 22 | 0.0059 | 1Score **> 13** indicates **identity** | U | K.KAFLNQAQIELR.L |
| 2892 | 147 | – | 157 | 651.8635 | 1301.7124 | 1301.7092 | 2.53 | 0 | 58 | 1.5e-06 | 1Score **> 13** indicates **identity** | U | K.AFLNQAQIELR.L |
| 2893 | 147 | – | 157 | 651.8636 | 1301.7126 | 1301.7092 | 2.69 | 0 | 72 | 5.9e-08 | 1Score **> 13** indicates **identity** | U | K.AFLNQAQIELR.L |
| 2894 | 147 | – | 157 | 651.8636 | 1301.7126 | 1301.7092 | 2.69 | 0 | 82 | 6.4e-09 | 1Score **> 13** indicates **identity** | U | K.AFLNQAQIELR.L |
| 2896 | 147 | – | 157 | 434.9119 | 1301.7139 | 1301.7092 | 3.63 | 0 | 61 | 7.4e-07 | 1Score **> 13** indicates **identity** | U | K.AFLNQAQIELR.L |
| 2899 | 147 | – | 157 | 652.3556 | 1302.6966 | 1301.7092 | 759 | 0 | 46 | 2.7e-05 | 1Score **> 13** indicates **identity** | U | K.AFLNQAQIELR.L |
| 2900 | 147 | – | 157 | 652.3567 | 1302.6988 | 1301.7092 | 760 | 0 | 46 | 2.6e-05 | 1Score **> 13** indicates **identity** | U | K.AFLNQAQIELR.L |
| 2901 | 147 | – | 157 | 652.3655 | 1302.7164 | 1301.7092 | 774 | 0 | 58 | 1.7e-06 | 1Score **> 13** indicates **identity** | U | K.AFLNQAQIELR.L |
| 3411 | 158 | – | 170 | 801.3890 | 1600.7634 | 1600.7589 | 2.85 | 0 | 54 | 4.1e-06 | 1Score **> 13** indicates **identity** | U | R.LLELMNQHDTEMK.Y |
| 3413 | 158 | – | 170 | 534.9304 | 1601.7694 | 1600.7589 | 631 | 0 | 40 | 0.0001 | 1Score **> 13** indicates **identity** | U | R.LLELMNQHDTEMK.Y |
| 3414 | 158 | – | 170 | 801.8930 | 1601.7714 | 1600.7589 | 633 | 0 | 42 | 6e-05 | 1Score **> 13** indicates **identity** | U | R.LLELMNQHDTEMK.Y |
| 4873 | 158 | – | 177 | 504.6646 | 2518.2866 | 2517.2759 | 401 | 1 | 31 | 0.00073 | 1Score **> 13** indicates **identity** | U | R.LLELMNQHDTEMKYYIVHLK.R |
| 4874 | 158 | – | 177 | 630.5797 | 2518.2897 | 2517.2759 | 403 | 1 | 25 | 0.0031 | 1Score **> 13** indicates **identity** | U | R.LLELMNQHDTEMKYYIVHLK.R |
| 5031 | 158 | – | 178 | 669.6042 | 2674.3877 | 2673.3770 | 378 | 2 | 81 | 7.9e-09 | 1Score **> 13** indicates **identity** | U | R.LLELMNQHDTEMKYYIVHLKR.H |
| 5032 | 158 | – | 178 | 535.8851 | 2674.3891 | 2673.3770 | 379 | 2 | 66 | 2.4e-07 | 1Score **> 13** indicates **identity** | U | R.LLELMNQHDTEMKYYIVHLKR.H |
| 5040 | 158 | – | 178 | 539.0845 | 2690.3861 | 2689.3720 | 377 | 2 | 31 | 0.00072 | 1Score **> 13** indicates **identity** | U | R.LLELMNQHDTEMKYYIVHLKR.H  + Oxidation (M) |
| 3256 | 203 | – | 215 | 379.4614 | 1513.8165 | 1513.8114 | 3.40 | 1 | 20 | 0.009 | 1Score **> 13** indicates **identity** | U | R.NTHFRGVSLNLTR.K |
| 3526 | 203 | – | 216 | 411.4850 | 1641.9109 | 1641.9063 | 2.79 | 2 | 20 | 0.011 | 1Score **> 13** indicates **identity** | U | R.NTHFRGVSLNLTRK.L |
| 4270 | 252 | – | 269 | 675.0256 | 2022.0550 | 2021.0476 | 498 | 2 | 60 | 1e-06 | 1Score **> 13** indicates **identity** | U | K.RSAIKIVDFGSSCQLGQR.I |
| 3964 | 253 | – | 269 | 933.4833 | 1864.9520 | 1864.9465 | 2.96 | 1 | 108 | 1.6e-11 | 1Score **> 13** indicates **identity** | U | R.SAIKIVDFGSSCQLGQR.I |
| 3965 | 253 | – | 269 | 933.4833 | 1864.9520 | 1864.9465 | 2.96 | 1 | 102 | 6.8e-11 | 1Score **> 13** indicates **identity** | U | R.SAIKIVDFGSSCQLGQR.I |
| 3971 | 253 | – | 269 | 622.9924 | 1865.9554 | 1864.9465 | 541 | 1 | 79 | 1.4e-08 | 1Score **> 13** indicates **identity** | U | R.SAIKIVDFGSSCQLGQR.I |
| 3972 | 253 | – | 269 | 622.9930 | 1865.9572 | 1864.9465 | 542 | 1 | 80 | 1e-08 | 1Score **> 13** indicates **identity** | U | R.SAIKIVDFGSSCQLGQR.I |
| 5180 | 253 | – | 277 | 730.3834 | 2917.5045 | 2916.4916 | 347 | 2 | 40 | 0.00011 | 1Score **> 13** indicates **identity** | U | R.SAIKIVDFGSSCQLGQRIYQYIQSR.F |
| 5181 | 253 | – | 277 | 973.5089 | 2917.5049 | 2916.4916 | 347 | 2 | 39 | 0.00013 | 1Score **> 13** indicates **identity** | U | R.SAIKIVDFGSSCQLGQRIYQYIQSR.F |
| 5227 | 253 | – | 277 | 750.3751 | 2997.4713 | 2996.4579 | 338 | 2 | 24 | 0.004 | 1Score **> 13** indicates **identity** | U | R.SAIKIVDFGSSCQLGQRIYQYIQSR.F  + Phospho (Y) |
| 3155 | 257 | – | 269 | 733.8577 | 1465.7008 | 1465.6984 | 1.68 | 0 | 58 | 1.5e-06 | 1Score **> 13** indicates **identity** | U | K.IVDFGSSCQLGQR.I |
| 3156 | 257 | – | 269 | 733.8585 | 1465.7024 | 1465.6984 | 2.78 | 0 | 85 | 3.4e-09 | 1Score **> 13** indicates **identity** | U | K.IVDFGSSCQLGQR.I |
| 3157 | 257 | – | 269 | 733.8586 | 1465.7026 | 1465.6984 | 2.91 | 0 | 79 | 1.3e-08 | 1Score **> 13** indicates **identity** | U | K.IVDFGSSCQLGQR.I |
| 3158 | 257 | – | 269 | 489.5750 | 1465.7032 | 1465.6984 | 3.27 | 0 | 47 | 2e-05 | 1Score **> 13** indicates **identity** | U | K.IVDFGSSCQLGQR.I |
| 4867 | 257 | – | 277 | 630.3210 | 2517.2549 | 2517.2434 | 4.55 | 1 | 32 | 0.00064 | 1Score **> 13** indicates **identity** | U | K.IVDFGSSCQLGQRIYQYIQSR.F |
| 4870 | 257 | – | 277 | 1260.1342 | 2518.2538 | 2517.2434 | 401 | 1 | 35 | 0.00029 | 1Score **> 13** indicates **identity** | U | K.IVDFGSSCQLGQRIYQYIQSR.F |
| 4871 | 257 | – | 277 | 1260.1351 | 2518.2556 | 2517.2434 | 402 | 1 | 30 | 0.00092 | 1Score **> 13** indicates **identity** | U | K.IVDFGSSCQLGQRIYQYIQSR.F |
| 4872 | 257 | – | 277 | 840.4261 | 2518.2565 | 2517.2434 | 402 | 1 | 43 | 4.8e-05 | 1Score **> 13** indicates **identity** | U | K.IVDFGSSCQLGQRIYQYIQSR.F |
| 4970 | 257 | – | 277 | 866.7443 | 2597.2111 | 2597.2098 | 0.50 | 1 | 16 | 0.027 | 1Score **> 13** indicates **identity** | U | K.IVDFGSSCQLGQRIYQYIQSR.F  + Phospho (Y) |
| 3305 | 270 | – | 280 | 768.9033 | 1535.7920 | 1535.7885 | 2.33 | 1 | 25 | 0.003 | 1Score **> 13** indicates **identity** | U | R.IYQYIQSRFYR.S |
| 3306 | 270 | – | 280 | 512.9396 | 1535.7970 | 1535.7885 | 5.54 | 1 | 17 | 0.021 | 1Score **> 13** indicates **identity** | U | R.IYQYIQSRFYR.S |
| 3310 | 270 | – | 280 | 769.4058 | 1536.7970 | 1535.7885 | 657 | 1 | 14 | 0.038 | 1Score **> 13** indicates **identity** | U | R.IYQYIQSRFYR.S |
| 3938 | 325 | – | 342 | 621.3538 | 1861.0396 | 1861.0383 | 0.69 | 0 | 56 | 2.3e-06 | 1Score **> 13** indicates **identity** | U | R.IVEVLGIPPAAMLDQAPK.A |
| 3939 | 325 | – | 342 | 621.3545 | 1861.0417 | 1861.0383 | 1.82 | 0 | 62 | 6.9e-07 | 1Score **> 13** indicates **identity** | U | R.IVEVLGIPPAAMLDQAPK.A |
| 3940 | 325 | – | 342 | 466.2678 | 1861.0421 | 1861.0383 | 2.05 | 0 | 58 | 1.4e-06 | 1Score **> 13** indicates **identity** | U | R.IVEVLGIPPAAMLDQAPK.A |
| 3941 | 325 | – | 342 | 931.5286 | 1861.0426 | 1861.0383 | 2.34 | 0 | 53 | 4.6e-06 | 1Score **> 13** indicates **identity** | U | R.IVEVLGIPPAAMLDQAPK.A |
| 3942 | 325 | – | 342 | 621.3553 | 1861.0441 | 1861.0383 | 3.11 | 0 | 58 | 1.7e-06 | 1Score **> 13** indicates **identity** | U | R.IVEVLGIPPAAMLDQAPK.A |
| 3943 | 325 | – | 342 | 931.5295 | 1861.0444 | 1861.0383 | 3.31 | 0 | 48 | 1.5e-05 | 1Score **> 13** indicates **identity** | U | R.IVEVLGIPPAAMLDQAPK.A |
| 3944 | 325 | – | 342 | 931.5299 | 1861.0452 | 1861.0383 | 3.74 | 0 | 64 | 3.8e-07 | 1Score **> 13** indicates **identity** | U | R.IVEVLGIPPAAMLDQAPK.A |
| 3945 | 325 | – | 342 | 621.3561 | 1861.0465 | 1861.0383 | 4.40 | 0 | 51 | 8.7e-06 | 1Score **> 13** indicates **identity** | U | R.IVEVLGIPPAAMLDQAPK.A |
| 3946 | 325 | – | 342 | 931.5318 | 1861.0490 | 1861.0383 | 5.78 | 0 | 54 | 3.9e-06 | 1Score **> 13** indicates **identity** | U | R.IVEVLGIPPAAMLDQAPK.A |
| 3949 | 325 | – | 342 | 932.0283 | 1862.0420 | 1861.0383 | 539 | 0 | 28 | 0.0016 | 1Score **> 13** indicates **identity** | U | R.IVEVLGIPPAAMLDQAPK.A |
| 3950 | 325 | – | 342 | 932.0295 | 1862.0444 | 1861.0383 | 541 | 0 | 36 | 0.00025 | 1Score **> 13** indicates **identity** | U | R.IVEVLGIPPAAMLDQAPK.A |
| 3951 | 325 | – | 342 | 621.6888 | 1862.0446 | 1861.0383 | 541 | 0 | 53 | 5.1e-06 | 1Score **> 13** indicates **identity** | U | R.IVEVLGIPPAAMLDQAPK.A |
| 3952 | 325 | – | 342 | 932.0301 | 1862.0456 | 1861.0383 | 541 | 0 | 37 | 0.00018 | 1Score **> 13** indicates **identity** | U | R.IVEVLGIPPAAMLDQAPK.A |
| 3953 | 325 | – | 342 | 932.0303 | 1862.0460 | 1861.0383 | 542 | 0 | 39 | 0.00014 | 1Score **> 13** indicates **identity** | U | R.IVEVLGIPPAAMLDQAPK.A |
| 3954 | 325 | – | 342 | 621.6895 | 1862.0467 | 1861.0383 | 542 | 0 | 55 | 3.1e-06 | 1Score **> 13** indicates **identity** | U | R.IVEVLGIPPAAMLDQAPK.A |
| 3955 | 325 | – | 342 | 932.0308 | 1862.0470 | 1861.0383 | 542 | 0 | 38 | 0.00017 | 1Score **> 13** indicates **identity** | U | R.IVEVLGIPPAAMLDQAPK.A |
| 3956 | 325 | – | 342 | 621.6898 | 1862.0476 | 1861.0383 | 542 | 0 | 51 | 7.7e-06 | 1Score **> 13** indicates **identity** | U | R.IVEVLGIPPAAMLDQAPK.A |
| 3957 | 325 | – | 342 | 932.0314 | 1862.0482 | 1861.0383 | 543 | 0 | 37 | 0.00022 | 1Score **> 13** indicates **identity** | U | R.IVEVLGIPPAAMLDQAPK.A |
| 3958 | 325 | – | 342 | 932.0316 | 1862.0486 | 1861.0383 | 543 | 0 | 34 | 0.00043 | 1Score **> 13** indicates **identity** | U | R.IVEVLGIPPAAMLDQAPK.A |
| 3959 | 325 | – | 342 | 932.0325 | 1862.0504 | 1861.0383 | 544 | 0 | 43 | 5e-05 | 1Score **> 13** indicates **identity** | U | R.IVEVLGIPPAAMLDQAPK.A |
| 4005 | 325 | – | 342 | 470.2672 | 1877.0397 | 1877.0332 | 3.46 | 0 | 30 | 0.0011 | 1Score **> 13** indicates **identity** | U | R.IVEVLGIPPAAMLDQAPK.A  + Oxidation (M) |
| 4007 | 325 | – | 342 | 940.0284 | 1878.0422 | 1877.0332 | 538 | 0 | 38 | 0.00014 | 1Score **> 13** indicates **identity** | U | R.IVEVLGIPPAAMLDQAPK.A  + Oxidation (M) |
| 4008 | 325 | – | 342 | 940.0284 | 1878.0422 | 1877.0332 | 538 | 0 | 35 | 0.00035 | 1Score **> 13** indicates **identity** | U | R.IVEVLGIPPAAMLDQAPK.A  + Oxidation (M) |
| 4009 | 325 | – | 342 | 940.0286 | 1878.0426 | 1877.0332 | 538 | 0 | 34 | 0.00043 | 1Score **> 13** indicates **identity** | U | R.IVEVLGIPPAAMLDQAPK.A  + Oxidation (M) |
| 4010 | 325 | – | 342 | 940.0287 | 1878.0428 | 1877.0332 | 538 | 0 | 33 | 0.00047 | 1Score **> 13** indicates **identity** | U | R.IVEVLGIPPAAMLDQAPK.A  + Oxidation (M) |
| 4011 | 325 | – | 342 | 627.0216 | 1878.0430 | 1877.0332 | 538 | 0 | 45 | 3.4e-05 | 1Score **> 13** indicates **identity** | U | R.IVEVLGIPPAAMLDQAPK.A  + Oxidation (M) |
| 4012 | 325 | – | 342 | 627.0217 | 1878.0433 | 1877.0332 | 538 | 0 | 43 | 4.6e-05 | 1Score **> 13** indicates **identity** | U | R.IVEVLGIPPAAMLDQAPK.A  + Oxidation (M) |
| 4013 | 325 | – | 342 | 940.0291 | 1878.0436 | 1877.0332 | 538 | 0 | 33 | 0.00048 | 1Score **> 13** indicates **identity** | U | R.IVEVLGIPPAAMLDQAPK.A  + Oxidation (M) |
| 4014 | 325 | – | 342 | 940.0292 | 1878.0438 | 1877.0332 | 538 | 0 | 26 | 0.0025 | 1Score **> 13** indicates **identity** | U | R.IVEVLGIPPAAMLDQAPK.A  + Oxidation (M) |
| 4374 | 325 | – | 344 | 697.0314 | 2088.0724 | 2088.1765 | -49.9 | 1 | 28 | 0.0016 | 1Score **> 13** indicates **identity** | U | R.IVEVLGIPPAAMLDQAPKAR.K |
| 4375 | 325 | – | 344 | 1045.0975 | 2088.1804 | 2088.1765 | 1.89 | 1 | 117 | 2.1e-12 | 1Score **> 13** indicates **identity** | U | R.IVEVLGIPPAAMLDQAPKAR.K |
| 4380 | 325 | – | 344 | 1045.5983 | 2089.1820 | 2088.1765 | 482 | 1 | 105 | 3e-11 | 1Score **> 13** indicates **identity** | U | R.IVEVLGIPPAAMLDQAPKAR.K |
| 4381 | 325 | – | 344 | 523.3031 | 2089.1833 | 2088.1765 | 482 | 1 | 56 | 2.4e-06 | 1Score **> 13** indicates **identity** | U | R.IVEVLGIPPAAMLDQAPKAR.K |
| 4382 | 325 | – | 344 | 523.3036 | 2089.1853 | 2088.1765 | 483 | 1 | 50 | 1e-05 | 1Score **> 13** indicates **identity** | U | R.IVEVLGIPPAAMLDQAPKAR.K |
| 4383 | 325 | – | 344 | 1045.6001 | 2089.1856 | 2088.1765 | 483 | 1 | 98 | 1.6e-10 | 1Score **> 13** indicates **identity** | U | R.IVEVLGIPPAAMLDQAPKAR.K |
| 4384 | 325 | – | 344 | 1045.6001 | 2089.1856 | 2088.1765 | 483 | 1 | 116 | 2.6e-12 | 1Score **> 13** indicates **identity** | U | R.IVEVLGIPPAAMLDQAPKAR.K |
| 4385 | 325 | – | 344 | 697.4025 | 2089.1857 | 2088.1765 | 483 | 1 | 88 | 1.7e-09 | 1Score **> 13** indicates **identity** | U | R.IVEVLGIPPAAMLDQAPKAR.K |
| 4386 | 325 | – | 344 | 523.3038 | 2089.1861 | 2088.1765 | 483 | 1 | 60 | 1.1e-06 | 1Score **> 13** indicates **identity** | U | R.IVEVLGIPPAAMLDQAPKAR.K |
| 4387 | 325 | – | 344 | 697.4028 | 2089.1866 | 2088.1765 | 484 | 1 | 64 | 3.9e-07 | 1Score **> 13** indicates **identity** | U | R.IVEVLGIPPAAMLDQAPKAR.K |
| 4388 | 325 | – | 344 | 697.4030 | 2089.1872 | 2088.1765 | 484 | 1 | 77 | 2.2e-08 | 1Score **> 13** indicates **identity** | U | R.IVEVLGIPPAAMLDQAPKAR.K |
| 4389 | 325 | – | 344 | 697.4030 | 2089.1872 | 2088.1765 | 484 | 1 | 77 | 2.2e-08 | 1Score **> 13** indicates **identity** | U | R.IVEVLGIPPAAMLDQAPKAR.K |
| 4390 | 325 | – | 344 | 1045.6016 | 2089.1886 | 2088.1765 | 485 | 1 | 116 | 2.6e-12 | 1Score **> 13** indicates **identity** | U | R.IVEVLGIPPAAMLDQAPKAR.K |
| 4391 | 325 | – | 344 | 697.4036 | 2089.1890 | 2088.1765 | 485 | 1 | 56 | 2.3e-06 | 1Score **> 13** indicates **identity** | U | R.IVEVLGIPPAAMLDQAPKAR.K |
| 4392 | 325 | – | 344 | 1045.6019 | 2089.1892 | 2088.1765 | 485 | 1 | 113 | 5.2e-12 | 1Score **> 13** indicates **identity** | U | R.IVEVLGIPPAAMLDQAPKAR.K |
| 4420 | 325 | – | 344 | 702.7339 | 2105.1799 | 2104.1714 | 479 | 1 | 76 | 2.4e-08 | 1Score **> 13** indicates **identity** | U | R.IVEVLGIPPAAMLDQAPKAR.K  + Oxidation (M) |
| 4421 | 325 | – | 344 | 527.3027 | 2105.1817 | 2104.1714 | 480 | 1 | 81 | 8.9e-09 | 1Score **> 13** indicates **identity** | U | R.IVEVLGIPPAAMLDQAPKAR.K  + Oxidation (M) |
| 4422 | 325 | – | 344 | 1053.5986 | 2105.1826 | 2104.1714 | 481 | 1 | 68 | 1.6e-07 | 1Score **> 13** indicates **identity** | U | R.IVEVLGIPPAAMLDQAPKAR.K  + Oxidation (M) |
| 4423 | 325 | – | 344 | 702.7358 | 2105.1856 | 2104.1714 | 482 | 1 | 100 | 9.9e-11 | 1Score **> 13** indicates **identity** | U | R.IVEVLGIPPAAMLDQAPKAR.K  + Oxidation (M) |
| 4523 | 325 | – | 345 | 555.0764 | 2216.2765 | 2216.2715 | 2.27 | 2 | 59 | 1.2e-06 | 1Score **> 13** indicates **identity** | U | R.IVEVLGIPPAAMLDQAPKARK.Y |
| 4524 | 325 | – | 345 | 555.3271 | 2217.2793 | 2216.2715 | 455 | 2 | 47 | 2e-05 | 1Score **> 13** indicates **identity** | U | R.IVEVLGIPPAAMLDQAPKARK.Y |
| 4525 | 325 | – | 345 | 740.1011 | 2217.2815 | 2216.2715 | 456 | 2 | 50 | 9.2e-06 | 1Score **> 13** indicates **identity** | U | R.IVEVLGIPPAAMLDQAPKARK.Y |
| 4526 | 325 | – | 345 | 555.3279 | 2217.2825 | 2216.2715 | 456 | 2 | 38 | 0.00014 | 1Score **> 13** indicates **identity** | U | R.IVEVLGIPPAAMLDQAPKARK.Y |
| 4527 | 325 | – | 345 | 740.1029 | 2217.2869 | 2216.2715 | 458 | 2 | 42 | 7e-05 | 1Score **> 13** indicates **identity** | U | R.IVEVLGIPPAAMLDQAPKARK.Y |
| 4536 | 325 | – | 345 | 559.3262 | 2233.2757 | 2232.2664 | 452 | 2 | 53 | 5e-06 | 1Score **> 13** indicates **identity** | U | R.IVEVLGIPPAAMLDQAPKARK.Y  + Oxidation (M) |
| 3583 | 345 | – | 358 | 560.6396 | 1678.8970 | 1678.8943 | 1.56 | 2 | 19 | 0.013 | 1Score **> 13** indicates **identity** | U | R.KYFERLPGGGWTLR.R |
| 3584 | 345 | – | 358 | 840.4561 | 1678.8976 | 1678.8943 | 1.97 | 2 | 36 | 0.00028 | 1Score **> 13** indicates **identity** | U | R.KYFERLPGGGWTLR.R |
| 3585 | 345 | – | 358 | 560.6400 | 1678.8982 | 1678.8943 | 2.28 | 2 | 20 | 0.0089 | 1Score **> 13** indicates **identity** | U | R.KYFERLPGGGWTLR.R |
| 3589 | 345 | – | 358 | 840.4579 | 1678.9012 | 1678.8943 | 4.11 | 2 | 41 | 8.4e-05 | 1Score **> 13** indicates **identity** | U | R.KYFERLPGGGWTLR.R |
| 3591 | 345 | – | 358 | 840.9585 | 1679.9024 | 1678.8943 | 600 | 2 | 28 | 0.0016 | 1Score **> 13** indicates **identity** | U | R.KYFERLPGGGWTLR.R |
| 3592 | 345 | – | 358 | 840.9585 | 1679.9024 | 1678.8943 | 600 | 2 | 30 | 0.0011 | 1Score **> 13** indicates **identity** | U | R.KYFERLPGGGWTLR.R |
| 3328 | 346 | – | 358 | 517.9421 | 1550.8045 | 1550.7994 | 3.28 | 1 | 26 | 0.0026 | 1Score **> 13** indicates **identity** | U | K.YFERLPGGGWTLR.R |
| 3329 | 346 | – | 358 | 776.4099 | 1550.8052 | 1550.7994 | 3.78 | 1 | 44 | 3.7e-05 | 1Score **> 13** indicates **identity** | U | K.YFERLPGGGWTLR.R |
| 3330 | 346 | – | 358 | 518.2753 | 1551.8041 | 1550.7994 | 648 | 1 | 21 | 0.0079 | 1Score **> 13** indicates **identity** | U | K.YFERLPGGGWTLR.R |
| 3331 | 346 | – | 358 | 776.9112 | 1551.8078 | 1550.7994 | 650 | 1 | 35 | 0.00033 | 1Score **> 13** indicates **identity** | U | K.YFERLPGGGWTLR.R |
| 3631 | 346 | – | 359 | 569.9728 | 1706.8966 | 1706.9005 | -2.30 | 2 | 22 | 0.0066 | 1Score **> 13** indicates **identity** | U | K.YFERLPGGGWTLRR.T |
| 3637 | 346 | – | 359 | 854.4604 | 1706.9062 | 1706.9005 | 3.37 | 2 | 27 | 0.002 | 1Score **> 13** indicates **identity** | U | K.YFERLPGGGWTLRR.T |
| 3641 | 346 | – | 359 | 854.4609 | 1706.9072 | 1706.9005 | 3.96 | 2 | 27 | 0.0021 | 1Score **> 13** indicates **identity** | U | K.YFERLPGGGWTLRR.T |
| 3779 | 346 | – | 359 | 596.6326 | 1786.8760 | 1786.8668 | 5.12 | 2 | 15 | 0.034 | 1Score **> 13** indicates **identity** | U | K.YFERLPGGGWTLRR.T  + Phospho (ST) |
| 2404 | 350 | – | 358 | 478.7712 | 955.5278 | 955.5240 | 4.07 | 0 | 52 | 6.7e-06 | 1Score **> 13** indicates **identity** | U | R.LPGGGWTLR.R |
| 2405 | 350 | – | 358 | 478.7712 | 955.5278 | 955.5240 | 4.07 | 0 | 52 | 6.7e-06 | 1Score **> 13** indicates **identity** | U | R.LPGGGWTLR.R |
| 2406 | 350 | – | 358 | 478.7713 | 955.5280 | 955.5240 | 4.28 | 0 | 48 | 1.7e-05 | 1Score **> 13** indicates **identity** | U | R.LPGGGWTLR.R |
| 2407 | 350 | – | 358 | 478.7721 | 955.5296 | 955.5240 | 5.96 | 0 | 50 | 1e-05 | 1Score **> 13** indicates **identity** | U | R.LPGGGWTLR.R |
| 2638 | 350 | – | 359 | 371.5499 | 1111.6279 | 1111.6251 | 2.53 | 1 | 16 | 0.022 | 1Score **> 13** indicates **identity** | U | R.LPGGGWTLRR.T |
| 2639 | 350 | – | 359 | 371.5499 | 1111.6279 | 1111.6251 | 2.53 | 1 | 19 | 0.012 | 1Score **> 13** indicates **identity** | U | R.LPGGGWTLRR.T |
| 2640 | 350 | – | 359 | 556.8214 | 1111.6282 | 1111.6251 | 2.87 | 1 | 22 | 0.0064 | 1Score **> 13** indicates **identity** | U | R.LPGGGWTLRR.T |
| 2641 | 350 | – | 359 | 556.8214 | 1111.6282 | 1111.6251 | 2.87 | 1 | 21 | 0.0073 | 1Score **> 13** indicates **identity** | U | R.LPGGGWTLRR.T |
| 2643 | 350 | – | 359 | 556.8218 | 1111.6290 | 1111.6251 | 3.59 | 1 | 17 | 0.018 | 1Score **> 13** indicates **identity** | U | R.LPGGGWTLRR.T |
| 4837 | 366 | – | 389 | 1249.6482 | 2497.2818 | 2497.2786 | 1.30 | 2 | 85 | 3.4e-09 | 1Score **> 13** indicates **identity** | U | K.DYQGPGTRRLQEVLGVQTGGPGGR.R |
| 4838 | 366 | – | 389 | 833.4353 | 2497.2841 | 2497.2786 | 2.19 | 2 | 51 | 8.9e-06 | 1Score **> 13** indicates **identity** | U | K.DYQGPGTRRLQEVLGVQTGGPGGR.R |
| 4839 | 366 | – | 389 | 1249.6510 | 2497.2874 | 2497.2786 | 3.54 | 2 | 63 | 4.7e-07 | 1Score **> 13** indicates **identity** | U | K.DYQGPGTRRLQEVLGVQTGGPGGR.R |
| 4841 | 366 | – | 389 | 833.7698 | 2498.2876 | 2497.2786 | 404 | 2 | 32 | 0.00057 | 1Score **> 13** indicates **identity** | U | K.DYQGPGTRRLQEVLGVQTGGPGGR.R |
| 4842 | 366 | – | 389 | 625.5793 | 2498.2881 | 2497.2786 | 404 | 2 | 14 | 0.036 | 1Score **> 13** indicates **identity** | U | K.DYQGPGTRRLQEVLGVQTGGPGGR.R |
| 4843 | 366 | – | 389 | 625.5797 | 2498.2897 | 2497.2786 | 405 | 2 | 14 | 0.042 | 1Score **> 13** indicates **identity** | U | K.DYQGPGTRRLQEVLGVQTGGPGGR.R |
| 3461 | 374 | – | 389 | 812.4507 | 1622.8868 | 1622.8853 | 0.97 | 1 | 64 | 3.6e-07 | 1Score **> 13** indicates **identity** | U | R.RLQEVLGVQTGGPGGR.R |
| 3462 | 374 | – | 389 | 541.9696 | 1622.8870 | 1622.8853 | 1.05 | 1 | 42 | 6.4e-05 | 1Score **> 13** indicates **identity** | U | R.RLQEVLGVQTGGPGGR.R |
| 3463 | 374 | – | 389 | 812.4509 | 1622.8872 | 1622.8853 | 1.22 | 1 | 67 | 1.8e-07 | 1Score **> 13** indicates **identity** | U | R.RLQEVLGVQTGGPGGR.R |
| 3464 | 374 | – | 389 | 541.9697 | 1622.8873 | 1622.8853 | 1.23 | 1 | 48 | 1.6e-05 | 1Score **> 13** indicates **identity** | U | R.RLQEVLGVQTGGPGGR.R |
| 3465 | 374 | – | 389 | 541.9697 | 1622.8873 | 1622.8853 | 1.23 | 1 | 43 | 5.1e-05 | 1Score **> 13** indicates **identity** | U | R.RLQEVLGVQTGGPGGR.R |
| 3466 | 374 | – | 389 | 406.7291 | 1622.8873 | 1622.8853 | 1.25 | 1 | 52 | 5.7e-06 | 1Score **> 13** indicates **identity** | U | R.RLQEVLGVQTGGPGGR.R |
| 3468 | 374 | – | 389 | 812.4515 | 1622.8884 | 1622.8853 | 1.96 | 1 | 74 | 4.2e-08 | 1Score **> 13** indicates **identity** | U | R.RLQEVLGVQTGGPGGR.R |
| 3469 | 374 | – | 389 | 812.4517 | 1622.8888 | 1622.8853 | 2.21 | 1 | 81 | 7.6e-09 | 1Score **> 13** indicates **identity** | U | R.RLQEVLGVQTGGPGGR.R |
| 3470 | 374 | – | 389 | 812.4517 | 1622.8888 | 1622.8853 | 2.21 | 1 | 69 | 1.3e-07 | 1Score **> 13** indicates **identity** | U | R.RLQEVLGVQTGGPGGR.R |
| 3471 | 374 | – | 389 | 812.4518 | 1622.8890 | 1622.8853 | 2.33 | 1 | 77 | 2e-08 | 1Score **> 13** indicates **identity** | U | R.RLQEVLGVQTGGPGGR.R |
| 3472 | 374 | – | 389 | 541.9703 | 1622.8891 | 1622.8853 | 2.34 | 1 | 53 | 5e-06 | 1Score **> 13** indicates **identity** | U | R.RLQEVLGVQTGGPGGR.R |
| 3473 | 374 | – | 389 | 541.9704 | 1622.8894 | 1622.8853 | 2.53 | 1 | 34 | 0.00039 | 1Score **> 13** indicates **identity** | U | R.RLQEVLGVQTGGPGGR.R |
| 3477 | 374 | – | 389 | 541.9705 | 1622.8897 | 1622.8853 | 2.71 | 1 | 46 | 2.5e-05 | 1Score **> 13** indicates **identity** | U | R.RLQEVLGVQTGGPGGR.R |
| 3479 | 374 | – | 389 | 541.9707 | 1622.8903 | 1622.8853 | 3.08 | 1 | 47 | 2.2e-05 | 1Score **> 13** indicates **identity** | U | R.RLQEVLGVQTGGPGGR.R |
| 3480 | 374 | – | 389 | 541.9708 | 1622.8906 | 1622.8853 | 3.27 | 1 | 52 | 6.5e-06 | 1Score **> 13** indicates **identity** | U | R.RLQEVLGVQTGGPGGR.R |
| 3481 | 374 | – | 389 | 541.9708 | 1622.8906 | 1622.8853 | 3.27 | 1 | 35 | 0.0003 | 1Score **> 13** indicates **identity** | U | R.RLQEVLGVQTGGPGGR.R |
| 3482 | 374 | – | 389 | 541.9709 | 1622.8909 | 1622.8853 | 3.45 | 1 | 44 | 4.1e-05 | 1Score **> 13** indicates **identity** | U | R.RLQEVLGVQTGGPGGR.R |
| 3483 | 374 | – | 389 | 812.4530 | 1622.8914 | 1622.8853 | 3.81 | 1 | 65 | 3.2e-07 | 1Score **> 13** indicates **identity** | U | R.RLQEVLGVQTGGPGGR.R |
| 3484 | 374 | – | 389 | 812.4530 | 1622.8914 | 1622.8853 | 3.81 | 1 | 65 | 3.3e-07 | 1Score **> 13** indicates **identity** | U | R.RLQEVLGVQTGGPGGR.R |
| 3486 | 374 | – | 389 | 541.9713 | 1622.8921 | 1622.8853 | 4.19 | 1 | 33 | 0.00047 | 1Score **> 13** indicates **identity** | U | R.RLQEVLGVQTGGPGGR.R |
| 3488 | 374 | – | 389 | 542.0063 | 1622.9971 | 1622.8853 | 68.9 | 1 | 15 | 0.032 | 1Score **> 13** indicates **identity** | U | R.RLQEVLGVQTGGPGGR.R |
| 3490 | 374 | – | 389 | 542.3043 | 1623.8911 | 1622.8853 | 620 | 1 | 37 | 0.00018 | 1Score **> 13** indicates **identity** | U | R.RLQEVLGVQTGGPGGR.R |
| 3491 | 374 | – | 389 | 406.9806 | 1623.8933 | 1622.8853 | 621 | 1 | 58 | 1.5e-06 | 1Score **> 13** indicates **identity** | U | R.RLQEVLGVQTGGPGGR.R |
| 3492 | 374 | – | 389 | 542.3052 | 1623.8938 | 1622.8853 | 621 | 1 | 35 | 0.00033 | 1Score **> 13** indicates **identity** | U | R.RLQEVLGVQTGGPGGR.R |
| 3628 | 374 | – | 389 | 568.9604 | 1703.8594 | 1702.8516 | 592 | 1 | 44 | 3.8e-05 | 1Score **> 13** indicates **identity** | U | R.RLQEVLGVQTGGPGGR.R  + Phospho (ST) |
| 3750 | 374 | – | 390 | 890.5005 | 1778.9864 | 1778.9864 | 0.042 | 2 | 23 | 0.0047 | 1Score **> 13** indicates **identity** | U | R.RLQEVLGVQTGGPGGRR.A |
| 3751 | 374 | – | 390 | 890.5014 | 1778.9882 | 1778.9864 | 1.05 | 2 | 59 | 1.3e-06 | 1Score **> 13** indicates **identity** | U | R.RLQEVLGVQTGGPGGRR.A |
| 3752 | 374 | – | 390 | 890.5015 | 1778.9884 | 1778.9864 | 1.17 | 2 | 71 | 7.9e-08 | 1Score **> 13** indicates **identity** | U | R.RLQEVLGVQTGGPGGRR.A |
| 3753 | 374 | – | 390 | 356.8051 | 1778.9891 | 1778.9864 | 1.54 | 2 | 42 | 6.4e-05 | 1Score **> 13** indicates **identity** | U | R.RLQEVLGVQTGGPGGRR.A |
| 3754 | 374 | – | 390 | 890.5019 | 1778.9892 | 1778.9864 | 1.62 | 2 | 71 | 7.2e-08 | 1Score **> 13** indicates **identity** | U | R.RLQEVLGVQTGGPGGRR.A |
| 3755 | 374 | – | 390 | 445.7546 | 1778.9893 | 1778.9864 | 1.64 | 2 | 54 | 4.1e-06 | 1Score **> 13** indicates **identity** | U | R.RLQEVLGVQTGGPGGRR.A |
| 3756 | 374 | – | 390 | 594.0038 | 1778.9896 | 1778.9864 | 1.80 | 2 | 54 | 4.1e-06 | 1Score **> 13** indicates **identity** | U | R.RLQEVLGVQTGGPGGRR.A |
| 3757 | 374 | – | 390 | 445.7547 | 1778.9897 | 1778.9864 | 1.87 | 2 | 52 | 6e-06 | 1Score **> 13** indicates **identity** | U | R.RLQEVLGVQTGGPGGRR.A |
| 3758 | 374 | – | 390 | 594.0039 | 1778.9899 | 1778.9864 | 1.97 | 2 | 66 | 2.6e-07 | 1Score **> 13** indicates **identity** | U | R.RLQEVLGVQTGGPGGRR.A |
| 3759 | 374 | – | 390 | 445.7548 | 1778.9901 | 1778.9864 | 2.09 | 2 | 48 | 1.7e-05 | 1Score **> 13** indicates **identity** | U | R.RLQEVLGVQTGGPGGRR.A |
| 3760 | 374 | – | 390 | 445.7548 | 1778.9901 | 1778.9864 | 2.09 | 2 | 26 | 0.0026 | 1Score **> 13** indicates **identity** | U | R.RLQEVLGVQTGGPGGRR.A |
| 3761 | 374 | – | 390 | 594.0040 | 1778.9902 | 1778.9864 | 2.13 | 2 | 47 | 2e-05 | 1Score **> 13** indicates **identity** | U | R.RLQEVLGVQTGGPGGRR.A |
| 3762 | 374 | – | 390 | 890.5024 | 1778.9902 | 1778.9864 | 2.18 | 2 | 59 | 1.1e-06 | 1Score **> 13** indicates **identity** | U | R.RLQEVLGVQTGGPGGRR.A |
| 3763 | 374 | – | 390 | 445.7549 | 1778.9905 | 1778.9864 | 2.32 | 2 | 45 | 2.9e-05 | 1Score **> 13** indicates **identity** | U | R.RLQEVLGVQTGGPGGRR.A |
| 3764 | 374 | – | 390 | 594.0042 | 1778.9908 | 1778.9864 | 2.47 | 2 | 44 | 4e-05 | 1Score **> 13** indicates **identity** | U | R.RLQEVLGVQTGGPGGRR.A |
| 3765 | 374 | – | 390 | 594.0043 | 1778.9911 | 1778.9864 | 2.64 | 2 | 54 | 3.9e-06 | 1Score **> 13** indicates **identity** | U | R.RLQEVLGVQTGGPGGRR.A |
| 3766 | 374 | – | 390 | 594.0045 | 1778.9917 | 1778.9864 | 2.98 | 2 | 60 | 1e-06 | 1Score **> 13** indicates **identity** | U | R.RLQEVLGVQTGGPGGRR.A |
| 3767 | 374 | – | 390 | 594.0050 | 1778.9932 | 1778.9864 | 3.82 | 2 | 44 | 3.9e-05 | 1Score **> 13** indicates **identity** | U | R.RLQEVLGVQTGGPGGRR.A |
| 3768 | 374 | – | 390 | 594.0051 | 1778.9935 | 1778.9864 | 3.99 | 2 | 48 | 1.7e-05 | 1Score **> 13** indicates **identity** | U | R.RLQEVLGVQTGGPGGRR.A |
| 3769 | 374 | – | 390 | 594.0066 | 1778.9980 | 1778.9864 | 6.52 | 2 | 52 | 5.6e-06 | 1Score **> 13** indicates **identity** | U | R.RLQEVLGVQTGGPGGRR.A |
| 3770 | 374 | – | 390 | 594.3098 | 1779.9076 | 1778.9864 | 518 | 2 | 35 | 0.00033 | 1Score **> 13** indicates **identity** | U | R.RLQEVLGVQTGGPGGRR.A |
| 3771 | 374 | – | 390 | 446.0055 | 1779.9929 | 1778.9864 | 566 | 2 | 31 | 0.0008 | 1Score **> 13** indicates **identity** | U | R.RLQEVLGVQTGGPGGRR.A |
| 3772 | 374 | – | 390 | 594.3387 | 1779.9943 | 1778.9864 | 567 | 2 | 53 | 4.7e-06 | 1Score **> 13** indicates **identity** | U | R.RLQEVLGVQTGGPGGRR.A |
| 3773 | 374 | – | 390 | 594.3398 | 1779.9976 | 1778.9864 | 568 | 2 | 34 | 0.00037 | 1Score **> 13** indicates **identity** | U | R.RLQEVLGVQTGGPGGRR.A |
| 3774 | 374 | – | 390 | 446.0069 | 1779.9985 | 1778.9864 | 569 | 2 | 41 | 8.7e-05 | 1Score **> 13** indicates **identity** | U | R.RLQEVLGVQTGGPGGRR.A |
| 3937 | 374 | – | 390 | 620.9939 | 1859.9599 | 1858.9527 | 542 | 2 | 48 | 1.6e-05 | 1Score **> 13** indicates **identity** | U | R.RLQEVLGVQTGGPGGRR.A  + Phospho (ST) |
| 3159 | 375 | – | 389 | 1467.7911 | 1466.7838 | 1466.7842 | -0.23 | 0 | 81 | 7.2e-09 | 1Score **> 13** indicates **identity** | U | R.LQEVLGVQTGGPGGR.R |
| 3160 | 375 | – | 389 | 489.9355 | 1466.7847 | 1466.7842 | 0.35 | 0 | 39 | 0.00011 | 1Score **> 13** indicates **identity** | U | R.LQEVLGVQTGGPGGR.R |
| 3161 | 375 | – | 389 | 734.4006 | 1466.7866 | 1466.7842 | 1.69 | 0 | 59 | 1.3e-06 | 1Score **> 13** indicates **identity** | U | R.LQEVLGVQTGGPGGR.R |
| 3162 | 375 | – | 389 | 734.4008 | 1466.7870 | 1466.7842 | 1.97 | 0 | 74 | 3.6e-08 | 1Score **> 13** indicates **identity** | U | R.LQEVLGVQTGGPGGR.R |
| 3163 | 375 | – | 389 | 734.4008 | 1466.7870 | 1466.7842 | 1.97 | 0 | 74 | 3.8e-08 | 1Score **> 13** indicates **identity** | U | R.LQEVLGVQTGGPGGR.R |
| 3164 | 375 | – | 389 | 489.9365 | 1466.7877 | 1466.7842 | 2.39 | 0 | 56 | 2.3e-06 | 1Score **> 13** indicates **identity** | U | R.LQEVLGVQTGGPGGR.R |
| 3165 | 375 | – | 389 | 734.4012 | 1466.7878 | 1466.7842 | 2.51 | 0 | 77 | 2.1e-08 | 1Score **> 13** indicates **identity** | U | R.LQEVLGVQTGGPGGR.R |
| 3166 | 375 | – | 389 | 734.4013 | 1466.7880 | 1466.7842 | 2.65 | 0 | 69 | 1.3e-07 | 1Score **> 13** indicates **identity** | U | R.LQEVLGVQTGGPGGR.R |
| 3167 | 375 | – | 389 | 734.4013 | 1466.7880 | 1466.7842 | 2.65 | 0 | 70 | 1e-07 | 1Score **> 13** indicates **identity** | U | R.LQEVLGVQTGGPGGR.R |
| 3168 | 375 | – | 389 | 1467.7954 | 1466.7881 | 1466.7842 | 2.70 | 0 | 81 | 7.9e-09 | 1Score **> 13** indicates **identity** | U | R.LQEVLGVQTGGPGGR.R |
| 3169 | 375 | – | 389 | 734.4016 | 1466.7886 | 1466.7842 | 3.06 | 0 | 76 | 2.4e-08 | 1Score **> 13** indicates **identity** | U | R.LQEVLGVQTGGPGGR.R |
| 3170 | 375 | – | 389 | 734.4017 | 1466.7888 | 1466.7842 | 3.19 | 0 | 71 | 7.9e-08 | 1Score **> 13** indicates **identity** | U | R.LQEVLGVQTGGPGGR.R |
| 3171 | 375 | – | 389 | 734.4017 | 1466.7888 | 1466.7842 | 3.19 | 0 | 75 | 3.4e-08 | 1Score **> 13** indicates **identity** | U | R.LQEVLGVQTGGPGGR.R |
| 3172 | 375 | – | 389 | 489.9369 | 1466.7889 | 1466.7842 | 3.21 | 0 | 37 | 0.00021 | 1Score **> 13** indicates **identity** | U | R.LQEVLGVQTGGPGGR.R |
| 3173 | 375 | – | 389 | 734.4020 | 1466.7894 | 1466.7842 | 3.60 | 0 | 63 | 4.7e-07 | 1Score **> 13** indicates **identity** | U | R.LQEVLGVQTGGPGGR.R |
| 3174 | 375 | – | 389 | 734.4023 | 1466.7900 | 1466.7842 | 4.01 | 0 | 76 | 2.7e-08 | 1Score **> 13** indicates **identity** | U | R.LQEVLGVQTGGPGGR.R |
| 3175 | 375 | – | 389 | 734.4023 | 1466.7900 | 1466.7842 | 4.01 | 0 | 74 | 3.6e-08 | 1Score **> 13** indicates **identity** | U | R.LQEVLGVQTGGPGGR.R |
| 3176 | 375 | – | 389 | 734.4023 | 1466.7900 | 1466.7842 | 4.01 | 0 | 73 | 5.2e-08 | 1Score **> 13** indicates **identity** | U | R.LQEVLGVQTGGPGGR.R |
| 3177 | 375 | – | 389 | 734.9036 | 1467.7926 | 1466.7842 | 688 | 0 | 64 | 4.5e-07 | 1Score **> 13** indicates **identity** | U | R.LQEVLGVQTGGPGGR.R |
| 3178 | 375 | – | 389 | 734.9036 | 1467.7926 | 1466.7842 | 688 | 0 | 46 | 2.4e-05 | 1Score **> 13** indicates **identity** | U | R.LQEVLGVQTGGPGGR.R |
| 3460 | 375 | – | 390 | 406.7289 | 1622.8865 | 1622.8853 | 0.76 | 1 | 41 | 7.2e-05 | 1Score **> 13** indicates **identity** | U | R.LQEVLGVQTGGPGGRR.A |
| 3467 | 375 | – | 390 | 541.9700 | 1622.8882 | 1622.8853 | 1.79 | 1 | 38 | 0.00016 | 1Score **> 13** indicates **identity** | U | R.LQEVLGVQTGGPGGRR.A |
| 3474 | 375 | – | 390 | 812.4520 | 1622.8894 | 1622.8853 | 2.58 | 1 | 44 | 3.9e-05 | 1Score **> 13** indicates **identity** | U | R.LQEVLGVQTGGPGGRR.A |
| 3475 | 375 | – | 390 | 812.4521 | 1622.8896 | 1622.8853 | 2.70 | 1 | 46 | 2.8e-05 | 1Score **> 13** indicates **identity** | U | R.LQEVLGVQTGGPGGRR.A |
| 3476 | 375 | – | 390 | 812.4521 | 1622.8896 | 1622.8853 | 2.70 | 1 | 52 | 7e-06 | 1Score **> 13** indicates **identity** | U | R.LQEVLGVQTGGPGGRR.A |
| 3478 | 375 | – | 390 | 541.9706 | 1622.8900 | 1622.8853 | 2.90 | 1 | 39 | 0.00014 | 1Score **> 13** indicates **identity** | U | R.LQEVLGVQTGGPGGRR.A |
| 3485 | 375 | – | 390 | 812.4532 | 1622.8918 | 1622.8853 | 4.05 | 1 | 58 | 1.6e-06 | 1Score **> 13** indicates **identity** | U | R.LQEVLGVQTGGPGGRR.A |
| 3487 | 375 | – | 390 | 541.9713 | 1622.8921 | 1622.8853 | 4.19 | 1 | 41 | 8.9e-05 | 1Score **> 13** indicates **identity** | U | R.LQEVLGVQTGGPGGRR.A |
| 3627 | 375 | – | 390 | 568.6245 | 1702.8517 | 1702.8516 | 0.046 | 1 | 50 | 1e-05 | 1Score **> 13** indicates **identity** | U | R.LQEVLGVQTGGPGGRR.A  + Phospho (ST) |
| 5212 | 375 | – | 403 | 595.9133 | 2974.5301 | 2973.5169 | 341 | 2 | 14 | 0.043 | 1Score **> 13** indicates **identity** | U | R.LQEVLGVQTGGPGGRRAGEPGHSPADYLR.F |
| 5262 | 375 | – | 403 | 764.6306 | 3054.4933 | 3053.4832 | 331 | 2 | 17 | 0.02 | 1Score **> 13** indicates **identity** | U | R.LQEVLGVQTGGPGGRRAGEPGHSPADYLR.F  + Phospho (ST) |
| 3278 | 390 | – | 403 | 509.2573 | 1524.7501 | 1524.7433 | 4.43 | 1 | 19 | 0.014 | 1Score **> 13** indicates **identity** | U | R.RAGEPGHSPADYLR.F |
| 3280 | 390 | – | 403 | 382.4444 | 1525.7485 | 1524.7433 | 659 | 1 | 66 | 2.6e-07 | 1Score **> 13** indicates **identity** | U | R.RAGEPGHSPADYLR.F |
| 4702 | 390 | – | 410 | 480.2556 | 2396.2416 | 2396.2349 | 2.80 | 2 | 23 | 0.0055 | 1Score **> 13** indicates **identity** | U | R.RAGEPGHSPADYLRFQDLVLR.M |
| 4703 | 390 | – | 410 | 600.0677 | 2396.2417 | 2396.2349 | 2.83 | 2 | 18 | 0.014 | 1Score **> 13** indicates **identity** | U | R.RAGEPGHSPADYLRFQDLVLR.M |
| 4704 | 390 | – | 410 | 799.7548 | 2396.2426 | 2396.2349 | 3.20 | 2 | 32 | 0.00066 | 1Score **> 13** indicates **identity** | U | R.RAGEPGHSPADYLRFQDLVLR.M |
| 4705 | 390 | – | 410 | 600.0684 | 2396.2445 | 2396.2349 | 4.00 | 2 | 16 | 0.024 | 1Score **> 13** indicates **identity** | U | R.RAGEPGHSPADYLRFQDLVLR.M |
| 4706 | 390 | – | 410 | 1199.1304 | 2396.2462 | 2396.2349 | 4.73 | 2 | 41 | 7.1e-05 | 1Score **> 13** indicates **identity** | U | R.RAGEPGHSPADYLRFQDLVLR.M |
| 4708 | 390 | – | 410 | 600.3175 | 2397.2409 | 2396.2349 | 420 | 2 | 14 | 0.044 | 1Score **> 13** indicates **identity** | U | R.RAGEPGHSPADYLRFQDLVLR.M |
| 4709 | 390 | – | 410 | 480.4556 | 2397.2416 | 2396.2349 | 420 | 2 | 26 | 0.0023 | 1Score **> 13** indicates **identity** | U | R.RAGEPGHSPADYLRFQDLVLR.M |
| 4710 | 390 | – | 410 | 800.0883 | 2397.2431 | 2396.2349 | 421 | 2 | 31 | 0.00084 | 1Score **> 13** indicates **identity** | U | R.RAGEPGHSPADYLRFQDLVLR.M |
| 4711 | 390 | – | 410 | 480.4560 | 2397.2436 | 2396.2349 | 421 | 2 | 22 | 0.0068 | 1Score **> 13** indicates **identity** | U | R.RAGEPGHSPADYLRFQDLVLR.M |
| 4712 | 390 | – | 410 | 800.0887 | 2397.2443 | 2396.2349 | 421 | 2 | 23 | 0.0049 | 1Score **> 13** indicates **identity** | U | R.RAGEPGHSPADYLRFQDLVLR.M |
| 4713 | 390 | – | 410 | 600.3184 | 2397.2445 | 2396.2349 | 421 | 2 | 13 | 0.05 | 1Score **> 13** indicates **identity** | U | R.RAGEPGHSPADYLRFQDLVLR.M |
| 4714 | 390 | – | 410 | 480.4562 | 2397.2446 | 2396.2349 | 421 | 2 | 24 | 0.0039 | 1Score **> 13** indicates **identity** | U | R.RAGEPGHSPADYLRFQDLVLR.M |
| 4716 | 390 | – | 410 | 600.3187 | 2397.2457 | 2396.2349 | 422 | 2 | 18 | 0.014 | 1Score **> 13** indicates **identity** | U | R.RAGEPGHSPADYLRFQDLVLR.M |
| 4718 | 390 | – | 410 | 1199.6315 | 2397.2484 | 2396.2349 | 423 | 2 | 28 | 0.0017 | 1Score **> 13** indicates **identity** | U | R.RAGEPGHSPADYLRFQDLVLR.M |
| 4806 | 390 | – | 410 | 620.0591 | 2476.2073 | 2476.2012 | 2.45 | 2 | 14 | 0.036 | 1Score **> 13** indicates **identity** | U | R.RAGEPGHSPADYLRFQDLVLR.M  + Phospho (ST) |
| 4807 | 390 | – | 410 | 496.4491 | 2477.2091 | 2476.2012 | 407 | 2 | 19 | 0.014 | 1Score **> 13** indicates **identity** | U | R.RAGEPGHSPADYLRFQDLVLR.M  + Phospho (ST) |
| 4808 | 390 | – | 410 | 620.3096 | 2477.2093 | 2476.2012 | 407 | 2 | 22 | 0.0061 | 1Score **> 13** indicates **identity** | U | R.RAGEPGHSPADYLRFQDLVLR.M  + Phospho (ST) |
| 4809 | 390 | – | 410 | 620.3102 | 2477.2117 | 2476.2012 | 408 | 2 | 19 | 0.012 | 1Score **> 13** indicates **identity** | U | R.RAGEPGHSPADYLRFQDLVLR.M  + Phospho (ST) |
| 4810 | 390 | – | 410 | 1239.6136 | 2477.2126 | 2476.2012 | 408 | 2 | 21 | 0.0088 | 1Score **> 13** indicates **identity** | U | R.RAGEPGHSPADYLRFQDLVLR.M  + Phospho (Y) |
| 4811 | 390 | – | 410 | 826.7450 | 2477.2132 | 2476.2012 | 409 | 2 | 15 | 0.032 | 1Score **> 13** indicates **identity** | U | R.RAGEPGHSPADYLRFQDLVLR.M  + Phospho (ST) |
| 4812 | 390 | – | 410 | 826.7452 | 2477.2138 | 2476.2012 | 409 | 2 | 27 | 0.0022 | 1Score **> 13** indicates **identity** | U | R.RAGEPGHSPADYLRFQDLVLR.M  + Phospho (ST) |
| 2987 | 391 | – | 403 | 685.3260 | 1368.6374 | 1368.6422 | -3.48 | 0 | 45 | 3.1e-05 | 1Score **> 13** indicates **identity** | U | R.AGEPGHSPADYLR.F |
| 2988 | 391 | – | 403 | 685.3304 | 1368.6462 | 1368.6422 | 2.95 | 0 | 74 | 4e-08 | 1Score **> 13** indicates **identity** | U | R.AGEPGHSPADYLR.F |
| 2989 | 391 | – | 403 | 457.2227 | 1368.6463 | 1368.6422 | 2.97 | 0 | 69 | 1.2e-07 | 1Score **> 13** indicates **identity** | U | R.AGEPGHSPADYLR.F |
| 2990 | 391 | – | 403 | 457.2227 | 1368.6463 | 1368.6422 | 2.97 | 0 | 17 | 0.022 | 1Score **> 13** indicates **identity** | U | R.AGEPGHSPADYLR.F |
| 2991 | 391 | – | 403 | 457.2228 | 1368.6466 | 1368.6422 | 3.19 | 0 | 73 | 4.6e-08 | 1Score **> 13** indicates **identity** | U | R.AGEPGHSPADYLR.F |
| 2994 | 391 | – | 403 | 685.8325 | 1369.6504 | 1368.6422 | 737 | 0 | 39 | 0.00013 | 1Score **> 13** indicates **identity** | U | R.AGEPGHSPADYLR.F |
| 3128 | 391 | – | 403 | 725.3127 | 1448.6108 | 1448.6085 | 1.60 | 0 | 28 | 0.0015 | 1Score **> 13** indicates **identity** | U | R.AGEPGHSPADYLR.F  + Phospho (ST) |
| 3129 | 391 | – | 403 | 483.8779 | 1448.6119 | 1448.6085 | 2.30 | 0 | 46 | 2.6e-05 | 1Score **> 13** indicates **identity** | U | R.AGEPGHSPADYLR.F  + Phospho (ST) |
| 3130 | 391 | – | 403 | 483.8781 | 1448.6125 | 1448.6085 | 2.72 | 0 | 45 | 3e-05 | 1Score **> 13** indicates **identity** | U | R.AGEPGHSPADYLR.F  + Phospho (ST) |
| 3131 | 391 | – | 403 | 725.3136 | 1448.6126 | 1448.6085 | 2.84 | 0 | 35 | 0.00029 | 1Score **> 13** indicates **identity** | U | R.AGEPGHSPADYLR.F  + Phospho (ST) |
| 4542 | 391 | – | 410 | 561.0353 | 2240.1121 | 2240.1338 | -9.69 | 1 | 23 | 0.0045 | 1Score **> 13** indicates **identity** | U | R.AGEPGHSPADYLRFQDLVLR.M |
| 4543 | 391 | – | 410 | 561.0392 | 2240.1277 | 2240.1338 | -2.73 | 1 | 17 | 0.021 | 1Score **> 13** indicates **identity** | U | R.AGEPGHSPADYLRFQDLVLR.M |
| 4544 | 391 | – | 410 | 561.0410 | 2240.1349 | 2240.1338 | 0.49 | 1 | 13 | 0.049 | 1Score **> 13** indicates **identity** | U | R.AGEPGHSPADYLRFQDLVLR.M |
| 4545 | 391 | – | 410 | 561.0411 | 2240.1353 | 2240.1338 | 0.67 | 1 | 15 | 0.034 | 1Score **> 13** indicates **identity** | U | R.AGEPGHSPADYLRFQDLVLR.M |
| 4546 | 391 | – | 410 | 561.0413 | 2240.1361 | 2240.1338 | 1.02 | 1 | 24 | 0.004 | 1Score **> 13** indicates **identity** | U | R.AGEPGHSPADYLRFQDLVLR.M |
| 4547 | 391 | – | 410 | 747.7197 | 2240.1373 | 2240.1338 | 1.55 | 1 | 19 | 0.014 | 1Score **> 13** indicates **identity** | U | R.AGEPGHSPADYLRFQDLVLR.M |
| 4548 | 391 | – | 410 | 1121.0764 | 2240.1382 | 2240.1338 | 1.98 | 1 | 28 | 0.0016 | 1Score **> 13** indicates **identity** | U | R.AGEPGHSPADYLRFQDLVLR.M |
| 4549 | 391 | – | 410 | 561.0422 | 2240.1397 | 2240.1338 | 2.63 | 1 | 33 | 0.00048 | 1Score **> 13** indicates **identity** | U | R.AGEPGHSPADYLRFQDLVLR.M |
| 4550 | 391 | – | 410 | 1121.0786 | 2240.1426 | 2240.1338 | 3.95 | 1 | 19 | 0.012 | 1Score **> 13** indicates **identity** | U | R.AGEPGHSPADYLRFQDLVLR.M |
| 4551 | 391 | – | 410 | 1121.0800 | 2240.1454 | 2240.1338 | 5.20 | 1 | 27 | 0.0021 | 1Score **> 13** indicates **identity** | U | R.AGEPGHSPADYLRFQDLVLR.M |
| 4556 | 391 | – | 410 | 561.2920 | 2241.1389 | 2240.1338 | 449 | 1 | 25 | 0.0034 | 1Score **> 13** indicates **identity** | U | R.AGEPGHSPADYLRFQDLVLR.M |
| 4557 | 391 | – | 410 | 561.2926 | 2241.1413 | 2240.1338 | 450 | 1 | 23 | 0.0045 | 1Score **> 13** indicates **identity** | U | R.AGEPGHSPADYLRFQDLVLR.M |
| 4558 | 391 | – | 410 | 748.0545 | 2241.1417 | 2240.1338 | 450 | 1 | 21 | 0.0071 | 1Score **> 13** indicates **identity** | U | R.AGEPGHSPADYLRFQDLVLR.M |
| 4559 | 391 | – | 410 | 561.2927 | 2241.1417 | 2240.1338 | 450 | 1 | 23 | 0.0047 | 1Score **> 13** indicates **identity** | U | R.AGEPGHSPADYLRFQDLVLR.M |
| 4560 | 391 | – | 410 | 748.0546 | 2241.1420 | 2240.1338 | 450 | 1 | 25 | 0.0029 | 1Score **> 13** indicates **identity** | U | R.AGEPGHSPADYLRFQDLVLR.M |
| 4561 | 391 | – | 410 | 748.0549 | 2241.1429 | 2240.1338 | 450 | 1 | 16 | 0.023 | 1Score **> 13** indicates **identity** | U | R.AGEPGHSPADYLRFQDLVLR.M |
| 4626 | 391 | – | 410 | 1161.0610 | 2320.1074 | 2320.1001 | 3.16 | 1 | 32 | 0.00062 | 1Score **> 13** indicates **identity** | U | R.AGEPGHSPADYLRFQDLVLR.M  + Phospho (ST) |
| 4627 | 391 | – | 410 | 774.7084 | 2321.1034 | 2320.1001 | 432 | 1 | 22 | 0.0069 | 1Score **> 13** indicates **identity** | U | R.AGEPGHSPADYLRFQDLVLR.M  + Phospho (ST) |
| 4628 | 391 | – | 410 | 774.7102 | 2321.1088 | 2320.1001 | 435 | 1 | 16 | 0.028 | 1Score **> 13** indicates **identity** | U | R.AGEPGHSPADYLRFQDLVLR.M  + Phospho (ST) |
| 4630 | 391 | – | 410 | 581.2849 | 2321.1105 | 2320.1001 | 435 | 1 | 25 | 0.0029 | 1Score **> 13** indicates **identity** | U | R.AGEPGHSPADYLRFQDLVLR.M  + Phospho (ST) |
| 4631 | 391 | – | 410 | 1161.5626 | 2321.1106 | 2320.1001 | 436 | 1 | 18 | 0.016 | 1Score **> 13** indicates **identity** | U | R.AGEPGHSPADYLRFQDLVLR.M  + Phospho (ST) |
| 2288 | 404 | – | 410 | 445.7586 | 889.5026 | 889.5022 | 0.56 | 0 | 19 | 0.014 | 1Score **> 13** indicates **identity** | U | R.FQDLVLR.M |
| 2289 | 404 | – | 410 | 890.5104 | 889.5031 | 889.5022 | 1.09 | 0 | 32 | 0.00056 | 1Score **> 13** indicates **identity** | U | R.FQDLVLR.M |
| 2290 | 404 | – | 410 | 445.7589 | 889.5032 | 889.5022 | 1.23 | 0 | 31 | 0.00083 | 1Score **> 13** indicates **identity** | U | R.FQDLVLR.M |
| 2291 | 404 | – | 410 | 445.7590 | 889.5034 | 889.5022 | 1.46 | 0 | 22 | 0.0067 | 1Score **> 13** indicates **identity** | U | R.FQDLVLR.M |
| 2292 | 404 | – | 410 | 445.7591 | 889.5036 | 889.5022 | 1.68 | 0 | 37 | 0.00021 | 1Score **> 13** indicates **identity** | U | R.FQDLVLR.M |
| 2293 | 404 | – | 410 | 890.5110 | 889.5037 | 889.5022 | 1.77 | 0 | 35 | 0.00031 | 1Score **> 13** indicates **identity** | U | R.FQDLVLR.M |
| 2294 | 404 | – | 410 | 445.7592 | 889.5038 | 889.5022 | 1.91 | 0 | 31 | 0.00071 | 1Score **> 13** indicates **identity** | U | R.FQDLVLR.M |
| 2295 | 404 | – | 410 | 890.5118 | 889.5045 | 889.5022 | 2.67 | 0 | 34 | 0.00041 | 1Score **> 13** indicates **identity** | U | R.FQDLVLR.M |
| 2296 | 404 | – | 410 | 445.7596 | 889.5046 | 889.5022 | 2.81 | 0 | 27 | 0.0018 | 1Score **> 13** indicates **identity** | U | R.FQDLVLR.M |
| 2297 | 404 | – | 410 | 445.7597 | 889.5048 | 889.5022 | 3.03 | 0 | 30 | 0.00098 | 1Score **> 13** indicates **identity** | U | R.FQDLVLR.M |
| 2298 | 404 | – | 410 | 445.7597 | 889.5048 | 889.5022 | 3.03 | 0 | 31 | 0.00077 | 1Score **> 13** indicates **identity** | U | R.FQDLVLR.M |
| 2299 | 404 | – | 410 | 445.7597 | 889.5048 | 889.5022 | 3.03 | 0 | 23 | 0.0047 | 1Score **> 13** indicates **identity** | U | R.FQDLVLR.M |
| 2300 | 404 | – | 410 | 445.7598 | 889.5050 | 889.5022 | 3.26 | 0 | 30 | 0.00095 | 1Score **> 13** indicates **identity** | U | R.FQDLVLR.M |
| 2301 | 404 | – | 410 | 445.7598 | 889.5050 | 889.5022 | 3.26 | 0 | 41 | 7.1e-05 | 1Score **> 13** indicates **identity** | U | R.FQDLVLR.M |
| 2302 | 404 | – | 410 | 445.7599 | 889.5052 | 889.5022 | 3.48 | 0 | 28 | 0.0017 | 1Score **> 13** indicates **identity** | U | R.FQDLVLR.M |
| 2303 | 404 | – | 410 | 445.7599 | 889.5052 | 889.5022 | 3.48 | 0 | 27 | 0.0022 | 1Score **> 13** indicates **identity** | U | R.FQDLVLR.M |
| 2304 | 404 | – | 410 | 445.7599 | 889.5052 | 889.5022 | 3.48 | 0 | 24 | 0.0042 | 1Score **> 13** indicates **identity** | U | R.FQDLVLR.M |
| 2305 | 404 | – | 410 | 890.5126 | 889.5053 | 889.5022 | 3.57 | 0 | 32 | 0.00057 | 1Score **> 13** indicates **identity** | U | R.FQDLVLR.M |
| 2306 | 404 | – | 410 | 445.7600 | 889.5054 | 889.5022 | 3.71 | 0 | 41 | 7.7e-05 | 1Score **> 13** indicates **identity** | U | R.FQDLVLR.M |
| 2307 | 404 | – | 410 | 445.7601 | 889.5056 | 889.5022 | 3.93 | 0 | 27 | 0.0021 | 1Score **> 13** indicates **identity** | U | R.FQDLVLR.M |
| 2308 | 404 | – | 410 | 445.7603 | 889.5060 | 889.5022 | 4.38 | 0 | 27 | 0.0021 | 1Score **> 13** indicates **identity** | U | R.FQDLVLR.M |
| 2309 | 404 | – | 410 | 446.2608 | 890.5070 | 889.5022 | 1130 | 0 | 21 | 0.0077 | 1Score **> 13** indicates **identity** | U | R.FQDLVLR.M |
| 4145 | 404 | – | 419 | 651.0096 | 1950.0070 | 1950.0033 | 1.89 | 1 | 30 | 0.00093 | 1Score **> 13** indicates **identity** | U | R.FQDLVLRMLEYEPAAR.I |
| 4146 | 404 | – | 419 | 976.0117 | 1950.0088 | 1950.0033 | 2.85 | 1 | 67 | 2e-07 | 1Score **> 13** indicates **identity** | U | R.FQDLVLRMLEYEPAAR.I |
| 4147 | 404 | – | 419 | 488.5095 | 1950.0089 | 1950.0033 | 2.88 | 1 | 28 | 0.0018 | 1Score **> 13** indicates **identity** | U | R.FQDLVLRMLEYEPAAR.I |
| 4148 | 404 | – | 419 | 651.0106 | 1950.0100 | 1950.0033 | 3.43 | 1 | 31 | 0.00077 | 1Score **> 13** indicates **identity** | U | R.FQDLVLRMLEYEPAAR.I |
| 4152 | 404 | – | 419 | 976.5127 | 1951.0108 | 1950.0033 | 517 | 1 | 56 | 2.7e-06 | 1Score **> 13** indicates **identity** | U | R.FQDLVLRMLEYEPAAR.I |
| 4153 | 404 | – | 419 | 976.5128 | 1951.0110 | 1950.0033 | 517 | 1 | 38 | 0.00016 | 1Score **> 13** indicates **identity** | U | R.FQDLVLRMLEYEPAAR.I |
| 4154 | 404 | – | 419 | 651.3453 | 1951.0141 | 1950.0033 | 518 | 1 | 31 | 0.00077 | 1Score **> 13** indicates **identity** | U | R.FQDLVLRMLEYEPAAR.I |
| 4185 | 404 | – | 419 | 984.0084 | 1966.0022 | 1965.9982 | 2.06 | 1 | 52 | 5.7e-06 | 1Score **> 13** indicates **identity** | U | R.FQDLVLRMLEYEPAAR.I  + Oxidation (M) |
| 4186 | 404 | – | 419 | 984.0098 | 1966.0050 | 1965.9982 | 3.48 | 1 | 54 | 4.4e-06 | 1Score **> 13** indicates **identity** | U | R.FQDLVLRMLEYEPAAR.I  + Oxidation (M) |
| 4187 | 404 | – | 419 | 656.3423 | 1966.0051 | 1965.9982 | 3.50 | 1 | 29 | 0.0012 | 1Score **> 13** indicates **identity** | U | R.FQDLVLRMLEYEPAAR.I  + Oxidation (M) |
| 4189 | 404 | – | 419 | 984.5108 | 1967.0070 | 1965.9982 | 513 | 1 | 31 | 0.00089 | 1Score **> 13** indicates **identity** | U | R.FQDLVLRMLEYEPAAR.I  + Oxidation (M) |
| 5410 | 404 | – | 432 | 844.7034 | 3374.7845 | 3373.7757 | 299 | 2 | 32 | 0.00058 | 1Score **> 13** indicates **identity** | U | R.FQDLVLRMLEYEPAARISPLGALQHGFFR.R |
| 5461 | 404 | – | 432 | 1157.9252 | 3470.7538 | 3469.7370 | 293 | 2 | 31 | 0.00077 | 1Score **> 13** indicates **identity** | U | R.FQDLVLRMLEYEPAARISPLGALQHGFFR.R  + Oxidation (M); Phospho (ST) |
| 5462 | 404 | – | 432 | 868.6965 | 3470.7569 | 3469.7370 | 294 | 2 | 33 | 0.00054 | 1Score **> 13** indicates **identity** | U | R.FQDLVLRMLEYEPAARISPLGALQHGFFR.R  + Oxidation (M); Phospho (ST) |
| 4851 | 411 | – | 432 | 835.1045 | 2502.2917 | 2502.2841 | 3.01 | 1 | 39 | 0.00013 | 1Score **> 13** indicates **identity** | U | R.MLEYEPAARISPLGALQHGFFR.R |
| 4854 | 411 | – | 432 | 626.8309 | 2503.2945 | 2502.2841 | 404 | 1 | 37 | 0.00021 | 1Score **> 13** indicates **identity** | U | R.MLEYEPAARISPLGALQHGFFR.R |
| 4942 | 411 | – | 432 | 862.0911 | 2583.2515 | 2582.2505 | 388 | 1 | 31 | 0.00077 | 1Score **> 13** indicates **identity** | U | R.MLEYEPAARISPLGALQHGFFR.R  + Phospho (ST) |
| 4943 | 411 | – | 432 | 517.6589 | 2583.2581 | 2582.2505 | 390 | 1 | 36 | 0.00028 | 1Score **> 13** indicates **identity** | U | R.MLEYEPAARISPLGALQHGFFR.R  + Phospho (ST) |
| 4944 | 411 | – | 432 | 862.0940 | 2583.2602 | 2582.2505 | 391 | 1 | 37 | 0.00019 | 1Score **> 13** indicates **identity** | U | R.MLEYEPAARISPLGALQHGFFR.R  + Phospho (ST) |
| 4945 | 411 | – | 432 | 862.0941 | 2583.2605 | 2582.2505 | 391 | 1 | 30 | 0.0009 | 1Score **> 13** indicates **identity** | U | R.MLEYEPAARISPLGALQHGFFR.R  + Phospho (ST) |
| 4946 | 411 | – | 432 | 862.0942 | 2583.2608 | 2582.2505 | 391 | 1 | 28 | 0.0016 | 1Score **> 13** indicates **identity** | U | R.MLEYEPAARISPLGALQHGFFR.R  + Phospho (ST) |
| 4947 | 411 | – | 432 | 646.8225 | 2583.2609 | 2582.2505 | 391 | 1 | 38 | 0.00014 | 1Score **> 13** indicates **identity** | U | R.MLEYEPAARISPLGALQHGFFR.R  + Phospho (ST) |
| 4948 | 411 | – | 432 | 862.0943 | 2583.2611 | 2582.2505 | 391 | 1 | 37 | 0.00022 | 1Score **> 13** indicates **identity** | U | R.MLEYEPAARISPLGALQHGFFR.R  + Phospho (ST) |
| 4949 | 411 | – | 432 | 646.8226 | 2583.2613 | 2582.2505 | 391 | 1 | 33 | 0.0005 | 1Score **> 13** indicates **identity** | U | R.MLEYEPAARISPLGALQHGFFR.R  + Phospho (ST) |
| 4950 | 411 | – | 432 | 646.8227 | 2583.2617 | 2582.2505 | 392 | 1 | 35 | 0.00035 | 1Score **> 13** indicates **identity** | U | R.MLEYEPAARISPLGALQHGFFR.R  + Phospho (ST) |
| 4951 | 411 | – | 432 | 1292.6382 | 2583.2618 | 2582.2505 | 392 | 1 | 35 | 0.00034 | 1Score **> 13** indicates **identity** | U | R.MLEYEPAARISPLGALQHGFFR.R  + Phospho (ST) |
| 4952 | 411 | – | 432 | 646.8229 | 2583.2625 | 2582.2505 | 392 | 1 | 33 | 0.00056 | 1Score **> 13** indicates **identity** | U | R.MLEYEPAARISPLGALQHGFFR.R  + Phospho (ST) |
| 4953 | 411 | – | 432 | 862.0949 | 2583.2629 | 2582.2505 | 392 | 1 | 32 | 0.0006 | 1Score **> 13** indicates **identity** | U | R.MLEYEPAARISPLGALQHGFFR.R  + Phospho (ST) |
| 4954 | 411 | – | 432 | 646.8230 | 2583.2629 | 2582.2505 | 392 | 1 | 35 | 0.00031 | 1Score **> 13** indicates **identity** | U | R.MLEYEPAARISPLGALQHGFFR.R  + Phospho (ST) |
| 4955 | 411 | – | 432 | 1292.6396 | 2583.2646 | 2582.2505 | 393 | 1 | 38 | 0.00016 | 1Score **> 13** indicates **identity** | U | R.MLEYEPAARISPLGALQHGFFR.R  + Phospho (ST) |
| 4956 | 411 | – | 432 | 862.0959 | 2583.2659 | 2582.2505 | 393 | 1 | 26 | 0.0023 | 1Score **> 13** indicates **identity** | U | R.MLEYEPAARISPLGALQHGFFR.R  + Phospho (ST) |
| 4971 | 411 | – | 432 | 867.4261 | 2599.2565 | 2598.2454 | 389 | 1 | 31 | 0.00083 | 1Score **> 13** indicates **identity** | U | R.MLEYEPAARISPLGALQHGFFR.R  + Oxidation (M); Phospho (ST) |
| 4972 | 411 | – | 432 | 867.4271 | 2599.2595 | 2598.2454 | 390 | 1 | 36 | 0.00023 | 1Score **> 13** indicates **identity** | U | R.MLEYEPAARISPLGALQHGFFR.R  + Oxidation (M); Phospho (ST) |
| 5071 | 411 | – | 433 | 548.6790 | 2738.3586 | 2738.3516 | 2.57 | 2 | 14 | 0.044 | 1Score **> 13** indicates **identity** | U | R.MLEYEPAARISPLGALQHGFFRR.T  + Phospho (ST) |
| 5072 | 411 | – | 433 | 457.5671 | 2739.3589 | 2738.3516 | 368 | 2 | 30 | 0.00091 | 1Score **> 13** indicates **identity** | U | R.MLEYEPAARISPLGALQHGFFRR.T  + Phospho (ST) |
| 5078 | 411 | – | 433 | 548.8799 | 2739.3631 | 2738.3516 | 369 | 2 | 22 | 0.007 | 1Score **> 13** indicates **identity** | U | R.MLEYEPAARISPLGALQHGFFRR.T  + Phospho (ST) |
| 5097 | 411 | – | 433 | 552.0781 | 2755.3541 | 2754.3465 | 366 | 2 | 24 | 0.0041 | 1Score **> 13** indicates **identity** | U | R.MLEYEPAARISPLGALQHGFFRR.T  + Oxidation (M); Phospho (ST) |
| 3108 | 420 | – | 432 | 721.8996 | 1441.7846 | 1441.7830 | 1.13 | 0 | 85 | 2.9e-09 | 1Score **> 13** indicates **identity** | U | R.ISPLGALQHGFFR.R |
| 3109 | 420 | – | 432 | 481.6026 | 1441.7860 | 1441.7830 | 2.05 | 0 | 65 | 2.9e-07 | 1Score **> 13** indicates **identity** | U | R.ISPLGALQHGFFR.R |
| 3110 | 420 | – | 432 | 481.6028 | 1441.7866 | 1441.7830 | 2.46 | 0 | 57 | 1.9e-06 | 1Score **> 13** indicates **identity** | U | R.ISPLGALQHGFFR.R |
| 3111 | 420 | – | 432 | 721.9006 | 1441.7866 | 1441.7830 | 2.52 | 0 | 84 | 3.7e-09 | 1Score **> 13** indicates **identity** | U | R.ISPLGALQHGFFR.R |
| 3112 | 420 | – | 432 | 721.9008 | 1441.7870 | 1441.7830 | 2.79 | 0 | 84 | 3.7e-09 | 1Score **> 13** indicates **identity** | U | R.ISPLGALQHGFFR.R |
| 3113 | 420 | – | 432 | 481.6030 | 1441.7872 | 1441.7830 | 2.88 | 0 | 58 | 1.5e-06 | 1Score **> 13** indicates **identity** | U | R.ISPLGALQHGFFR.R |
| 3114 | 420 | – | 432 | 721.9011 | 1441.7876 | 1441.7830 | 3.21 | 0 | 71 | 8.4e-08 | 1Score **> 13** indicates **identity** | U | R.ISPLGALQHGFFR.R |
| 3115 | 420 | – | 432 | 481.6032 | 1441.7878 | 1441.7830 | 3.30 | 0 | 58 | 1.5e-06 | 1Score **> 13** indicates **identity** | U | R.ISPLGALQHGFFR.R |
| 3116 | 420 | – | 432 | 481.6032 | 1441.7878 | 1441.7830 | 3.30 | 0 | 45 | 3.1e-05 | 1Score **> 13** indicates **identity** | U | R.ISPLGALQHGFFR.R |
| 3117 | 420 | – | 432 | 481.6032 | 1441.7878 | 1441.7830 | 3.30 | 0 | 46 | 2.4e-05 | 1Score **> 13** indicates **identity** | U | R.ISPLGALQHGFFR.R |
| 3118 | 420 | – | 432 | 721.9012 | 1441.7878 | 1441.7830 | 3.35 | 0 | 79 | 1.4e-08 | 1Score **> 13** indicates **identity** | U | R.ISPLGALQHGFFR.R |
| 3119 | 420 | – | 432 | 721.9015 | 1441.7884 | 1441.7830 | 3.76 | 0 | 87 | 1.8e-09 | 1Score **> 13** indicates **identity** | U | R.ISPLGALQHGFFR.R |
| 3120 | 420 | – | 432 | 721.9055 | 1441.7964 | 1441.7830 | 9.31 | 0 | 55 | 3.3e-06 | 1Score **> 13** indicates **identity** | U | R.ISPLGALQHGFFR.R |
| 3123 | 420 | – | 432 | 481.9368 | 1442.7886 | 1441.7830 | 697 | 0 | 60 | 9e-07 | 1Score **> 13** indicates **identity** | U | R.ISPLGALQHGFFR.R |
| 3124 | 420 | – | 432 | 481.9371 | 1442.7895 | 1441.7830 | 698 | 0 | 41 | 7.7e-05 | 1Score **> 13** indicates **identity** | U | R.ISPLGALQHGFFR.R |
| 3274 | 420 | – | 432 | 761.8839 | 1521.7532 | 1521.7493 | 2.57 | 0 | 69 | 1.1e-07 | 1Score **> 13** indicates **identity** | U | R.ISPLGALQHGFFR.R  + Phospho (ST) |
| 3275 | 420 | – | 432 | 761.8850 | 1521.7554 | 1521.7493 | 4.01 | 0 | 61 | 7.5e-07 | 1Score **> 13** indicates **identity** | U | R.ISPLGALQHGFFR.R  + Phospho (ST) |
| 3383 | 420 | – | 433 | 799.9506 | 1597.8866 | 1597.8841 | 1.58 | 1 | 33 | 0.00051 | 1Score **> 13** indicates **identity** | U | R.ISPLGALQHGFFRR.T |
| 3384 | 420 | – | 433 | 533.6365 | 1597.8877 | 1597.8841 | 2.22 | 1 | 26 | 0.0026 | 1Score **> 13** indicates **identity** | U | R.ISPLGALQHGFFRR.T |
| 3385 | 420 | – | 433 | 400.4792 | 1597.8877 | 1597.8841 | 2.23 | 1 | 21 | 0.0085 | 1Score **> 13** indicates **identity** | U | R.ISPLGALQHGFFRR.T |
| 3386 | 420 | – | 433 | 400.4792 | 1597.8877 | 1597.8841 | 2.23 | 1 | 21 | 0.0074 | 1Score **> 13** indicates **identity** | U | R.ISPLGALQHGFFRR.T |
| 3387 | 420 | – | 433 | 400.4793 | 1597.8881 | 1597.8841 | 2.49 | 1 | 27 | 0.0018 | 1Score **> 13** indicates **identity** | U | R.ISPLGALQHGFFRR.T |
| 3388 | 420 | – | 433 | 400.4793 | 1597.8881 | 1597.8841 | 2.49 | 1 | 17 | 0.019 | 1Score **> 13** indicates **identity** | U | R.ISPLGALQHGFFRR.T |
| 3389 | 420 | – | 433 | 799.9514 | 1597.8882 | 1597.8841 | 2.58 | 1 | 31 | 0.00077 | 1Score **> 13** indicates **identity** | U | R.ISPLGALQHGFFRR.T |
| 3390 | 420 | – | 433 | 400.4794 | 1597.8885 | 1597.8841 | 2.74 | 1 | 25 | 0.0034 | 1Score **> 13** indicates **identity** | U | R.ISPLGALQHGFFRR.T |
| 3391 | 420 | – | 433 | 533.6368 | 1597.8886 | 1597.8841 | 2.78 | 1 | 31 | 0.00081 | 1Score **> 13** indicates **identity** | U | R.ISPLGALQHGFFRR.T |
| 3392 | 420 | – | 433 | 533.6368 | 1597.8886 | 1597.8841 | 2.78 | 1 | 20 | 0.011 | 1Score **> 13** indicates **identity** | U | R.ISPLGALQHGFFRR.T |
| 3393 | 420 | – | 433 | 400.4795 | 1597.8889 | 1597.8841 | 2.99 | 1 | 22 | 0.0068 | 1Score **> 13** indicates **identity** | U | R.ISPLGALQHGFFRR.T |
| 3394 | 420 | – | 433 | 400.4795 | 1597.8889 | 1597.8841 | 2.99 | 1 | 21 | 0.0084 | 1Score **> 13** indicates **identity** | U | R.ISPLGALQHGFFRR.T |
| 3395 | 420 | – | 433 | 400.4795 | 1597.8889 | 1597.8841 | 2.99 | 1 | 21 | 0.0072 | 1Score **> 13** indicates **identity** | U | R.ISPLGALQHGFFRR.T |
| 3396 | 420 | – | 433 | 533.6370 | 1597.8892 | 1597.8841 | 3.16 | 1 | 20 | 0.01 | 1Score **> 13** indicates **identity** | U | R.ISPLGALQHGFFRR.T |
| 3397 | 420 | – | 433 | 400.4796 | 1597.8893 | 1597.8841 | 3.24 | 1 | 20 | 0.01 | 1Score **> 13** indicates **identity** | U | R.ISPLGALQHGFFRR.T |
| 3398 | 420 | – | 433 | 400.4796 | 1597.8893 | 1597.8841 | 3.24 | 1 | 27 | 0.0022 | 1Score **> 13** indicates **identity** | U | R.ISPLGALQHGFFRR.T |
| 3399 | 420 | – | 433 | 400.4796 | 1597.8893 | 1597.8841 | 3.24 | 1 | 20 | 0.0096 | 1Score **> 13** indicates **identity** | U | R.ISPLGALQHGFFRR.T |
| 3400 | 420 | – | 433 | 400.4796 | 1597.8893 | 1597.8841 | 3.24 | 1 | 23 | 0.0053 | 1Score **> 13** indicates **identity** | U | R.ISPLGALQHGFFRR.T |
| 3401 | 420 | – | 433 | 533.6378 | 1597.8916 | 1597.8841 | 4.66 | 1 | 25 | 0.003 | 1Score **> 13** indicates **identity** | U | R.ISPLGALQHGFFRR.T |
| 3402 | 420 | – | 433 | 533.9690 | 1598.8852 | 1597.8841 | 626 | 1 | 18 | 0.016 | 1Score **> 13** indicates **identity** | U | R.ISPLGALQHGFFRR.T |
| 3403 | 420 | – | 433 | 533.9698 | 1598.8876 | 1597.8841 | 628 | 1 | 17 | 0.022 | 1Score **> 13** indicates **identity** | U | R.ISPLGALQHGFFRR.T |
| 3404 | 420 | – | 433 | 400.7299 | 1598.8905 | 1597.8841 | 630 | 1 | 18 | 0.014 | 1Score **> 13** indicates **identity** | U | R.ISPLGALQHGFFRR.T |
| 3405 | 420 | – | 433 | 400.7301 | 1598.8913 | 1597.8841 | 630 | 1 | 23 | 0.0049 | 1Score **> 13** indicates **identity** | U | R.ISPLGALQHGFFRR.T |
| 3406 | 420 | – | 433 | 400.7305 | 1598.8929 | 1597.8841 | 631 | 1 | 17 | 0.021 | 1Score **> 13** indicates **identity** | U | R.ISPLGALQHGFFRR.T |
| 3579 | 420 | – | 433 | 839.9349 | 1677.8552 | 1677.8504 | 2.86 | 1 | 15 | 0.029 | 1Score **> 13** indicates **identity** | U | R.ISPLGALQHGFFRR.T  + Phospho (ST) |
| 3580 | 420 | – | 433 | 839.9371 | 1677.8596 | 1677.8504 | 5.48 | 1 | 23 | 0.0056 | 1Score **> 13** indicates **identity** | U | R.ISPLGALQHGFFRR.T  + Phospho (ST) |

---

**Mascot:** http://www.matrixscience.com/


Phospho (Y) (+79.9663)
