## Supplementary figures and images for "Structural perspective on the design of selective DYRK1B inhibitors"

### 88x31_logo_white.gif

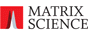

### external_arrow.png

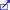

### mass_error.pl.pobrane

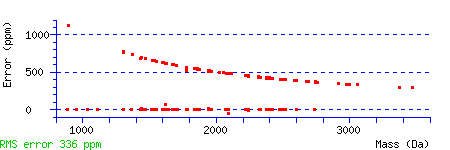
